# Naturomimetic conditions expose an unusual oxygen–iron regulatory network in *Rhodopseudomonas* and related *α-proteobacteria*

**DOI:** 10.64898/2026.06.27.734994

**Authors:** Brian Gallagher, Tahina Ranaivoarisoa, Prem Prabhakar, Jingtao Li, Anjali Rajkumar, Dinesh Gupta, Josh Kim, Arpita Bose

## Abstract

*Rhodopseudomonas palustris* (*Rp*.) is a metabolically versatile environmental bacterium that flourishes across gradients of iron, oxygen, and light. The varying metabolic landscapes *Rp*. exploits have pressured it to evolve hierarchical regulatory networks, enabling tuneable expression in response to multiple input signals. However, many studies of regulation in this organism have used standard laboratory conditions: incandescent illumination and single-substrate growth, which do not fully engage the complexity of its regulatory networks and lack relevance to natural environments. *Rp*. can generate energy through light-powered iron oxidation via the PioABC system, and swathes of its anaerobic metabolism employ proteins using ferrous cofactors produced from bioavailable Fe(II). *Rp*. lacks the Fe(II)-sensing regulators found in related lineages, leaving it unclear how it balances expression of anaerobic metabolic pathways with sensed iron levels. Using what we term *naturomimetic conditions*, including growth with simultaneous provision of an organic carbon source and Fe(II), we uncovered a surprising link between oxygen-responsive and iron-responsive regulatory networks in *Rp*. FixK and AadR mediate the regulatory shift from aerobic to anaerobic metabolism, but, as we demonstrate here, they also orchestrate the response to Fe(II) by modulating a secondary network of Fur-family regulators. Δ*aadR* and Δ*fixK* strains were both defective in iron metabolism, and Δ*aadR* showed perturbed expression of all four Fur-family regulators, sometimes in entirely opposite directions. Our findings establish that iron regulation in *Rp*. occurs via a regulatory network integrating multiple environmental signals. Phylogenetic analyses suggest conservation of network components across several related evolutionary lineages, including *Bradyrhizobium*.

**Graphical Abstract:** 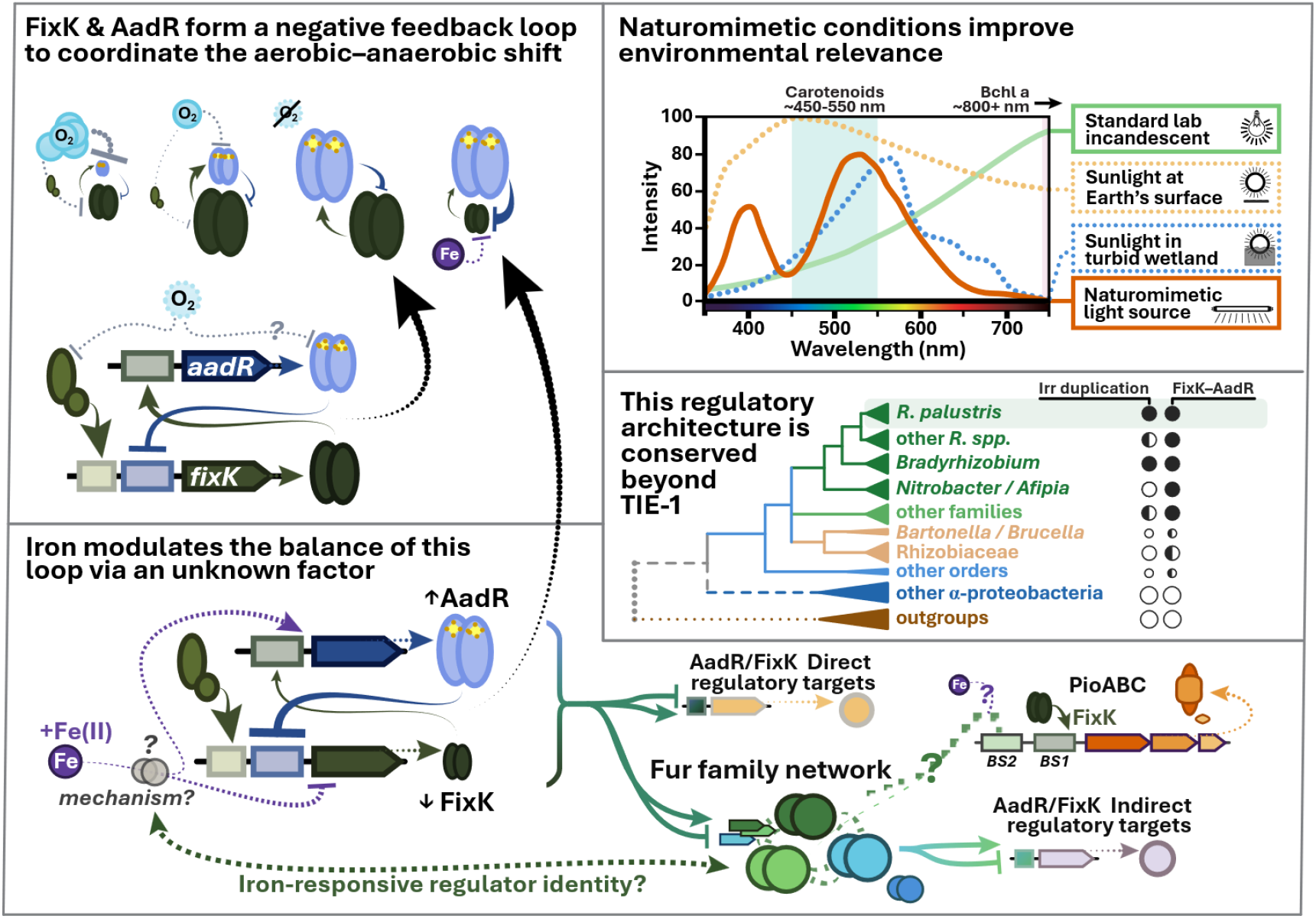

## Introduction

*Rhodopseudomonas palustris* is a ubiquitous and free-living phototrophic purple non-sulphur bacterium renowned for its metabolic versatility across varying environments.^1^ *R. palustris* inhabits wetland sediments and water columns,^2,3^ microbial mats,^4^ and wastewaters,^5^ where it is exposed to a variety of environmental gradients. As a “metabolic opportunist”,^1^ it uses various sources of carbon, electrons, and energy. It grows aerobically or anaerobically, and along the microaerobic transition, performing photoheterotrophy (on a variety of organic carbon sources, including aromatic ones), photoautotrophy, chemoheterotrophy, and/or nitrogen fixation as needed.^1,6,7^ To ensure fitness by expressing only what the environment demands, *R. palustris* uses a complex network of transcriptional regulators, often several regulatory layers deep.^1,8^ Despite the promise of its varied biotechnological and ecological capabilities (e.g., as a biofertilizer, for bioremediation, etc.), applications of *R. palustris* have been challenged by its elaborate regulatory network.^9^ Disentangling this complexity would unlock a powerful chassis for biotechnology and grant insights into how metabolically-versatile purple phototrophic bacteria sense and respond to their natural environments.

To ensure realistic and practical conclusions, regulatory studies should emulate the conditions the organism would experience in real-world conditions.^10^ Thus, we first developed a set of what we term *naturomimetic conditions* (i.e., mimicking nature) based on meta-analysis of the literature. The light source and concentrations of iron and organic carbon were carefully chosen to better simulate a real wetland, instead of optimizing for robust laboratory growth. Sensing and responding to these environmental factors underpins the metabolic choices *R. palustris* makes: for example, when provided both acetate and chelated Fe(II), it typically consumes the acetate before it begins utilising the iron.^11,12^ We included this condition to simulate the simultaneous substrate availability of natural environments, allowing us to interrogate the regulatory mechanisms underpinning this observation. Finally, we inoculated most anaerobic growth experiments from aerobically grown cultures, enforcing the aerobic-to-anaerobic regulatory shift to see how it intersects with the rest of the regulatory network.

The aerobic-to-anaerobic regulatory shift in *R. palustris* is mediated by the CRP/FNR-family transcriptional regulators FixK and AadR.^13^ Two oxygen-sensing schemes operate in *α-proteobacteria*: the ancestral one uses an [Fe-S]-bearing FNR that directly senses O_2_, while the scheme found in *R. palustris* and relatives involves a FixL(-J)-FixK_(2)_ cascade terminating with AadR (FNR/FixK_1_).^14–19^ The FixL-J two-component sensor activates expression of *fixK* as oxygen decreases, at which point FixK activates expression at numerous other loci, including *aadR*.^13^ AadR represses further *fixK* transcription in a negative feedback loop to complete the shift to an anaerobic regulatory program.^13,15^ The two FNR-family regulators are paralogues, and the similarity of their DNA-binding domains has precluded determination of distinct regulons for each (**Table ST1**), but Rey and Harwood (2010) identified dozens of loci in *R. palustris* strain CGA009 that are directly controlled by one or both regulators.^13^ They observed that AadR’s sphere of influence extends beyond a role in anaerobic aromatic degradation, broadly touching many anaerobic metabolic pathways.^13,20,21^ Deletion of *aadR* derepressed ferric iron transport genes, putative Fur-family regulator *irrA*, and what was later discovered to be the *fez* operon encoding the iron-storage ferrosome.^13,22^ Thus, we hypothesized that AadR (and by extension, FixK) may play a role in iron-responsive regulation.

Iron-responsive regulation has undergone substantial lineage-specific remodelling in *α*-proteobacteria.^23–29^ In taxonomic order Hyphomicrobiales, the ancestral Fur (ferric uptake regulator) has functionally diverged to become the manganese uptake regulator Mur, while IscR (Iron-sulphur cluster regulator) has been lost entirely.^14,23,29–31^ In the *Rhizobiaceae* family, the acquisition of the regulator RirA (*Rhizobiaceae* iron regulator) has filled the resulting regulatory gap as a direct Fe(II)–binding regulator.^29,30^ The sister family *Nitrobacteraceae*, including *R. palustris*, lacks RirA entirely.^30^ Members of this family are thought to manage iron-responsive regulation exclusively through the heme-sensing Irr, predicted to bind sites at many loci controlled by RirA in relatives with that regulator.^23,25,28^ Irr activates genes involved with Fe^3+^ uptake when iron is limited, and is deactivated or marked for degradation when binding haem.^14,32^ *R. palustris* possesses two Irr paralogues, like most *Nitrobacteraceae*, named *irrA1* and *irrA2* here (**Fig. S1**).^23,28^ IrrA1 retains the conserved high-affinity haem recognition motif, while IrrA2 does not, leaving its role less certain.^28^ Iron, often in the form of chelated Fe(II), is important for the anaerobic metabolism of *R. palustris*, but as it lacks a dedicated Fe(II) sensor such as Fur or RirA, the method by which it coordinates iron-responsive gene expression in response to Fe(II) remains an open question.

The *pioABC* operon provides a compelling case study for the question of iron-responsive regulation in *R. palustris*. One of its most distinctive metabolic capabilities is photoferrotrophy: a form of photoautotrophy involving carbon dioxide fixation, light as an energy source, and electrons for reducing power sourced from the oxidation of reduced iron (Fe(II)).^33–35^ In *R. palustris* strain TIE-1 (TIE-1), the PioABC protein complex facilitates this process, named for the phototrophic oxidation of iron it enables.^4,34–36^ While many *pioABC*- encoding strains can oxidise Fe(II) when growing heterotrophically on an organic carbon source, the ability to grow photoferrotrophically is not equally widespread.^34,35,37^ The *pioABC-*homologous *mtrCAB* in *Shewanella oneidensis* is regulated by Fur, but no Fur-family regulators have yet been implicated in *pioABC* regulation in *R. palustris*.^38,39^ The *pioABC* promoter (P*pio*) instead contains two confirmed transcription factor binding sites with the AT-rich palindromic motifs usually bound by CRP/FNR-family regulators, henceforth termed Binding Sites 1 (BS1) and 2 (BS2).^34^ P*pio-*BS1 lies at −44.5 relative to the transcription start site and is bound by FixK protein *in vitro*, while P*pio-*BS2 lies upstream of it (at −137.5 relative to TSS) and is not bound by FixK (**Fig. S2**).^34^ Deletion of BS2 reduced expression by 60-80% and abolished the iron-responsiveness of regulation by the promoter, suggesting that the unknown BS2-binding factor is either iron-responsive or under the control of an iron-responsive regulator.^34^ Intriguingly, FixK-driven activation of *pioABC* transcription occurs specifically under photoferrotrophy, implicating a second regulator (likely one capable of sensing iron), but its identity has not yet been determined.^34^

In this work, we compared growth and transcriptional regulation in wild-type (WT) TIE-1 and in regulator deletion (Δ) mutants. We performed experiments under naturomimetic growth conditions, carefully tuning lighting, carbon sources, oxygen, and iron availability to better reflect the conditions *R. palustris* experiences in nature to reveal growth differences and regulatory connections that might otherwise be missed under routine laboratory growth conditions. We show that AadR coordinates iron-responsive gene regulation as an additional regulatory layer linked with Fur-family regulators, and that the pathways differentially regulated in its absence are significantly enriched for genes encoding both iron-requiring and iron-related proteins. The Δ*aadR* strain was severely defective in iron oxidation, and it refused to grow entirely under photoferrotrophic conditions. The effects of AadR on *pioABC* expression are real but indirect — narrowing the search for the unknown *Ppio-*BS2-binding factor. Finally, we examine the dispersion of this regulatory network architecture across α-proteobacteria, finding several related lineages in which it is likely conserved.

## Methods

### Strains, mutants, and plasmids

Wild-type TIE-1 was originally isolated by Jiao *et al*. (2005).^40^ Markerless deletion mutants (Δ*fixK*, Δ*badM*) were generated from this background using methods described previously, using suicide plasmid pJQ200KS (**Fig. S20A-F**).^41,42^ Combination mutants were constructed from the single-mutant backgrounds. Mutants were confirmed by PCR and whole-genome sequencing (Plasmidsaurus). Earlier work in *R. palustris* strain CGA009 failed to construct a clean Δ*aadR* mutant, despite observing no aerobic growth defect,^20^ and our own repeated failures to create a markerless deletion strain in TIE-1 compelled deletion of *aadR* via insertion of a Pc promoter-driven *kanR* cassette, cloned between 1000 bp *aadR* flanking regions into pJQ200KS (**Fig. S20G-I**; HiFi assembly, New England Biolabs, Inc.). The ribosome binding site of the gene downstream of *aadR* (and its continued transcription) were confirmed to be preserved in Δ*aadR*. Gene neighbourhoods of all regulators of interest in wild-type and mutant TIE-1 are explicitly discussed in **Fig. S20**. Interestingly, a CGA009 transposon insertion library held no strains with a random insertion breaking *aadR*, even as insertions into the neighbouring genes yielded dozens (*n* > 120) (**Fig. S17, SI Dataset S05**).^43,44^ Together, these observations suggest strong counter-selection at the *aadR* locus in *R. palustris*. Complementation plasmids were constructed in the pSRKGm expression vector (under a modified P*lac* inducible promoter),^45^ and were transformed into *R. palustris* via conjugation with *E. coli* WM6026 (**Figs. S4, S7–S8**).^46^ A complete list of strains, plasmids, and primers used in this work may be found in **SI Tables ST4**-**5**.

### Microbial culture conditions

Aerobic cultures were grown in yeast-peptone succinate medium (YPS: 3 g/L yeast extract, 3 g/L peptone, 1 mM succinate) buffered with 10 mM MOPS [3-(*N*-morpholino)propanesulphonic acid], pH 7.0, 30°C, shaking at 250 rpm. Anaerobic phototrophic growth was performed under our naturomimetic conditions (**SI 1.1-1.3**). Briefly, strains were grown in Balch tubes or 100-mL serum bottles (for iron-oxidizing conditions), at 30°C, under fluorescent lighting, and in bicarbonate-buffered freshwater (FW) medium prepared as previously described.^33,34,47^ FW was amended with 10 mM sodium acetate (FWAc), 10 mM sodium malate (FWMal), 1 mM sodium benzoate (FWB), or 10 mM sodium succinate (FW-S), from filter-sterilised anaerobic 1 M stocks. Iron-supplemented conditions (FWAc+Fe, FWMal+Fe, FW+Fe) were amended with 5 mM FeCl_2_ and equimolar NTA from anaerobic stocks. Pre-growth for photoheterotrophy was performed aerobically in YPS to better capture the aerobic-anaerobic metabolic shift phenotype. Pre-growth for photoferrotrophy was performed on hydrogen, using a mixed gas (80:20 H_2_:CO_2_) headspace at 34.5 kPa. Purity of Δ*aadR* strains was maintained with 20 μg/mL kanamycin selection, which was omitted during growth experiments. Complement strains were grown with 400 μg/mL gentamicin and 10 mM IPTG (Isopropyl β-D-1-thiogalactopyranoside) to ensure retention of complementation vectors and expression of complemented genes, respectively.

### Growth curves and iron oxidation curves

Growth was monitored by OD_600_ (Spectronic 200, Thermo Fisher Scientific). Growth curves were modelled using a modified Gompertz equation, with doubling times calculated as described in **SI 1.4**.^48^ Statistical comparisons of growth parameters used two-way ANOVA with Tukey’s HSD (R v4.5.1). Growth data represent averages of biological replicates (*n*=3-6) with 95% confidence intervals (CI). Fe(II) concentrations were measured by ferrozine assay: 100-µL anaerobic samples were acidified in 1M HCl, reacted with ferrozine reagent, and measured at 562 nm (BioTek, Agilent).^49^ Concentrations were calculated from a standard curve, normalised to abiotic controls, and averaged across technical (*n*=2) and biological (*n*=2–3) replicates with standard error propagated. Extended details are in **SI 1.4**.

### RNA extraction and sequencing

Cells were harvested at mid-exponential phase (0.5 < OD_660_ < 1.0) from *n*=2 biological replicates per strain per condition, centrifuged in RNA-Later buffer, and stored at −80°C. RNA was extracted using the RNeasy Mini Kit (Qiagen). Library preparation and RNA-seq were performed by the Genome Technology Access Center at the McDonnell Genome Institute (GTAC@MGI). Quality was assessed by TapeStation (Agilent), libraries were prepared with ribosomal depletion (FastSelect Bacteria), and sequencing was performed on an Illumina NovaSeq X Plus (300 cycles, ∼15M targeted reads/sample). Demultiplexed raw reads were processed on KBase: adapter-trimmed and quality-filtered (JGI RQCFilter, BBTools v38.22), aligned to the TIE-1 genome (RefSeq:NC_011004) with HISAT2 v2.1.0 (with splicing/intron settings disabled), and assembled with StringTie v2.1.5. Transcripts per million (TPM) and raw count matrices were exported.

### Differential expression and principal component analysis (PCA)

Differential expression was assessed on KBase using the DESeq2 app, and DE matrices were exported.^50,51^ DESeq2 was chosen for its strong performance in calculating reliable and stable dispersion estimates even from biological duplicates, ensuring stringent statistical validity of the findings and preventing inflated false positive rates.^50,52^ The high-magnitude, high-confidence effects reported in this study obviate detection of small effects and low-abundance transcripts granted by inclusion of more replicates.^52^ The meta-analyses with independent datasets and statistical robustness of contrasts central to the conclusions indicate that no qPCR confirmation is required.^53^ Pairwise comparisons were performed with a single-factor design (∼condition); significance was assessed by Wald test with Benjamini-Hochberg (BH) false discovery rate (FDR) correction (reported as *q*-values). For WT-vs-mutant comparisons, positive log_2_FC indicates higher expression in wild-type. Read counts were normalised for principal component analysis (PCA) and z-scoring using variance-stabilising transformation (VST; pyDESeq2 v0.5.2) on raw counts,^54^ standardised to zero mean/unit variance (scikit-learn v1.7.2).^55^ Pearson correlations with environmental factors were computed across all 28 samples (SciPy v1.15.3).^56^ Displacements were calculated as Euclidean distance between wild-type and mutant centroids in PC1-PC2 space.

### Genome and proteome annotation

The TIE-1 proteome was annotated for iron-related genes using FeGenie v1.0 with its full iron HMM library,^57^ and for iron-requiring genes using HMMER v3.4 with 20 curated Pfam profiles representing Fe-S clusters, haem, radical SAM, and related iron-binding domains.^58^ Predictions were reconciled with MetalPredator metal-binding domain calls and supplemented with manual curation using known iron-coordinating proteins cited in the literature.^59^ These gene sets (435 iron-related, 174 iron-requiring) were tested for enrichment in the WT/Δ*aadR* and WT/Δ*badM* differential expression datasets using one-sided Fisher’s exact tests and Mann-Whitney U tests on |log_2_FC| distributions (DEGs had |log_2_FC| > 1 and *q* < 0.05). Test comparisons were restricted to genes differentially expressed between aerobic and basal-iron anaerobic conditions. BH-FDR correction was applied across all tests. Full analysis details in **Dataset S03**.

### Data Availability Statement

A supporting information appendix containing additional figures, tables, and methods is attached. Raw RNA sequencing reads are in NCBI as an SRA (BioProject: PRJNA1492185). Analytical code is attached and is available as a GitShare repository. Processed datasets from this study including gene lists, annotations, and analysis results are in **SI Datasets S01-S09**.

## Results

### Naturomimetic growth reveals oxygen- and iron-dependent phenotypes in mutant strains

For a real-world understanding of microbial phototrophic growth, we analysed how the variables light, iron, and oxygen could be tuned to best represent natural environments, which we term naturomimetic conditions (**SI 1**). The attenuated, yellow-shifted light that penetrates into a wetland water column or sediment is better mimicked by certain fluorescent lights than it is by the incandescent tungsten bulb lamps commonly used for phototrophic growth in laboratories (**SI 1.1, Fig. S3**).^60,61^ Oxygen, iron, and carbon sources were also used in environmentally-relevant forms and concentrations.

While no significant growth defect was observed for any mutant strain under aerobic conditions (YPS, **Fig. S4**), the Δ*aadR*Δ*fixK* double mutant was synthetically lethal under anaerobic phototrophic conditions, despite both Δ*aadR* and Δ*fixK* single mutants being capable of anaerobic growth (**Figs. S5–S8**). Δ*fixK* showed defective growth during photoheterotrophy, as expected based on past work (**Table ST2, Fig. S9**).^34^ Δ*aadR* showed the expected defect on benzoate but it surprisingly also showed a significant defect on succinate photoheterotrophy (**Figs. S6–S7**), contrary to what was reported in CGA009.^13,20^ Knocking out either *fixK* or *aadR* significantly reduced expression of light-harvesting genes under our conditions, implying that the growth defects caused by these mutations may be exacerbated by the reduced photon flux of naturomimetic lighting compared with the conventional incandescent illumination used in past work (**SI 1.1, Fig. S9**).^13,34^

Iron-supplemented (**“+Fe”**) conditions had 5 mM FeCl_2_ with equimolar nitrilotriacetic acid (NTA), emulating the millimolar-scale Fe(II) in groundwater-fed wetland sediments,^62–64^ while basal iron (**“–Fe”**) conditions contained only ∼4 μM Fe as a trace element (**SI 1.2**).^47^ As predicted from previous studies, TIE-1 grew photoheterotrophically using the acetate before it began performing measurable iron oxidation, permitting collection of RNA during the photoheterotrophic phase of FWAc+Fe growth (as evidenced by ferrozine assay results; **Fig 1A, C**; 0.5 < OD_660_ < 1.0 at harvest; **SI 1.3**).^11,12^ Thus, the transcriptome reported for FWAc+Fe in this study directly reflects the response to supplemental Fe(II) levels, not to iron oxidation metabolism. This avoids a confounding limitation inherent in past methods, which only directly compared photoheterotrophy with photoferrotrophy.

**Figure 1.**
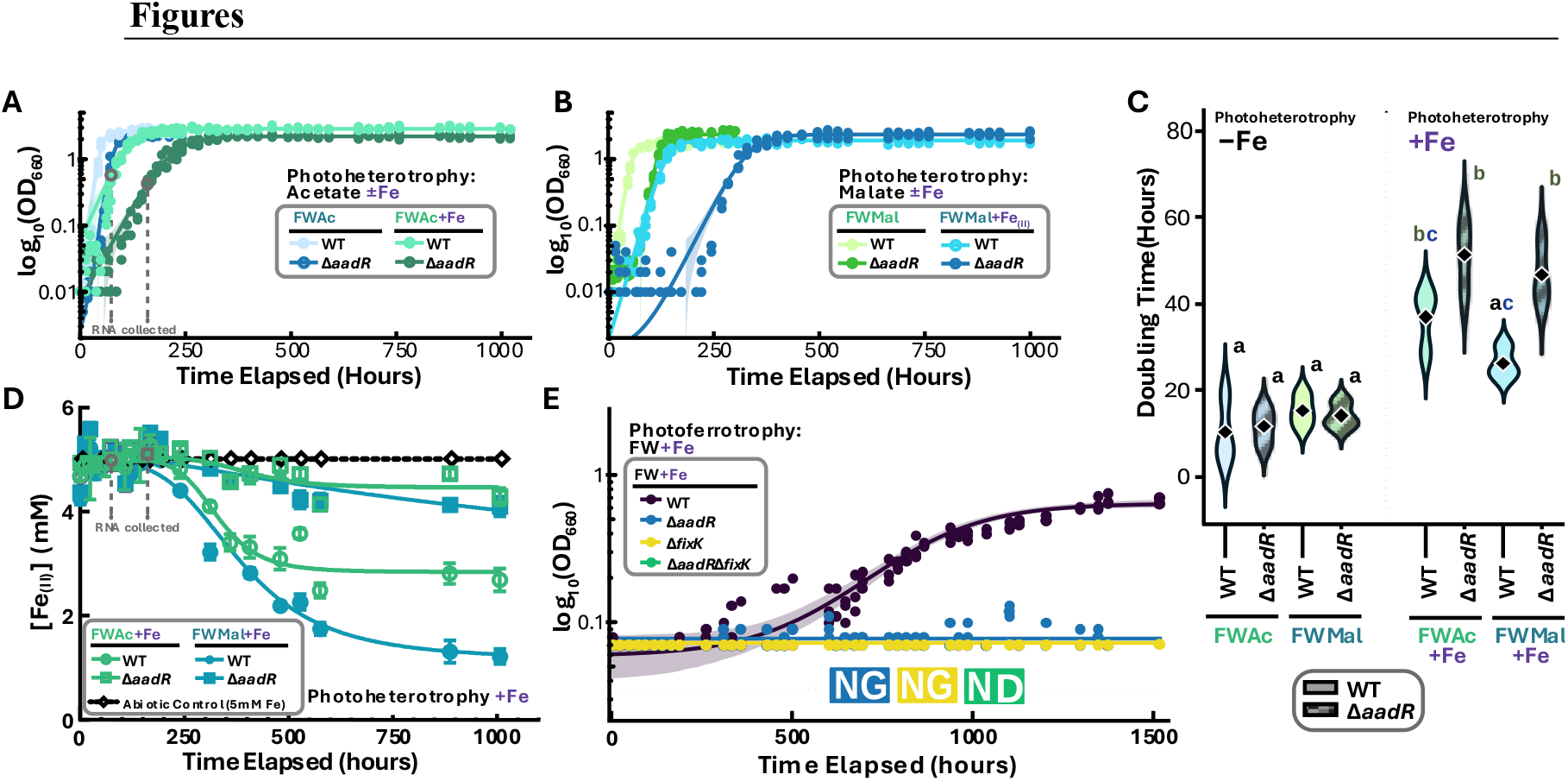
AadR and FixK are important for growth of TIE-1 under iron-supplemented conditions. Growth curves of wild-type and Δ*aadR* TIE-1 on **(A)** FWAcetate or **(B)** FWMalate. Δ*aadR* showed a slight defect in lag time, but no significant defect in doubling time relative to wild-type under photoheterotrophy with basal iron (–Fe) on either carbon source. Growth curves for each strain on each carbon source are also reported for supplemental iron (+Fe, +5 mM) conditions. **(C)** Doubling time for each strain on each carbon source under either –Fe or +Fe concentrations. CLD Letters represent the results of a two-way ANOVA (Tukey-adjusted) on calculated doubling time. **(D)** Oxidation curves for the +Fe growth experiments, measured via the ferrozine assay (see Methods) and normalised to the abiotic control. Fe oxidation curves were calculated as asymmetric sigmoidal 5-parameter lines fit to the data, with iron oxidation rate and final amount of iron oxidised both significantly decreased in Δ*aadR* relative to wild-type (**Table ST3**). For wildtype, significantly less iron was oxidised by the end of growth with acetate compared to with malate, though the rate at which it was oxidised did not significantly differ. **(E)** Growth curve for FW+Fe(II) photoferrotrophy, with a FW-Hydrogen photoautotrophic pre-growth. The Δ*aadR*Δ*fixK* double mutant did not grow on the pre-growth (“NG” = no growth), so its performance on photoferrotrophy was not determined (“ND” = not determined). While Δ*aadR* and Δ*fixK* grew in the pre-growth, neither began iron-oxidising photoferrotrophic growth.

+Fe photoheterotrophy revealed more severe phenotypes than displayed under –Fe growth. Δ*aadR* showed a significant growth defect on both FWAc+Fe and FWMal+Fe, despite growing comparably to wild-type on both carbon sources with basal iron levels (–Fe) (**Fig. 1A, B, C**). Δ*aadR* also oxidised significantly less Fe(II) than wild-type, with less than 1 mM oxidised over the course of the experiment during growth on either carbon source, compared to ∼2.5 mM oxidised by WT on acetate and ∼4.5 mM on malate (**Fig. 1D**). Δ*fixK* failed to grow at all under +Fe conditions (**Fig. 1E, S5, S8**), contrasting the delayed but successful photoferrotrophy it exhibited under high-intensity incandescent light.^34^ Neither single mutant grew on photoferrotrophy (FW+Fe) (**Fig. 1E, S8**). These iron-dependent growth defects, particularly the conditional Δ*aadR* defect on acetate and malate (absent under –Fe but pronounced under +Fe, **Fig. 1C**), motivated our transcriptomic investigation of a hypothetical iron-related regulatory role of AadR.

### Δ*aadR* disrupts the FixK−AadR balance, perturbing Fur-family transcriptional regulators

During photoferrotrophic growth, the balance between *fixK* and *aadR* transcript levels shifts in favour of *aadR* (**Fig. 2A**). Activation of *aadR* and repression of *fixK* under photoferrotrophy was replicated in the independent Guzman *et al*. (2019) dataset.^65^ In photoheterotrophy with supplemental Fe(II), wild-type TIE-1 continued to show repressed transcription of *fixK* (log_2_FC = –0.73, *q =* 0.002) and activation *aadR* (log_2_FC = +1.40, q< 10^−15^) (**Fig. 2B**). Critically, this iron-mediated repression of *fixK* was abolished in Δ*aadR* (log_2_FC = +0.20, *not significant*), supporting the possibility that this repression occurs through the established mechanism of AadR’s direct repression of *fixK* in a negative feedback loop in response to iron, not photoferrotrophy.^13^

**Figure 2.**
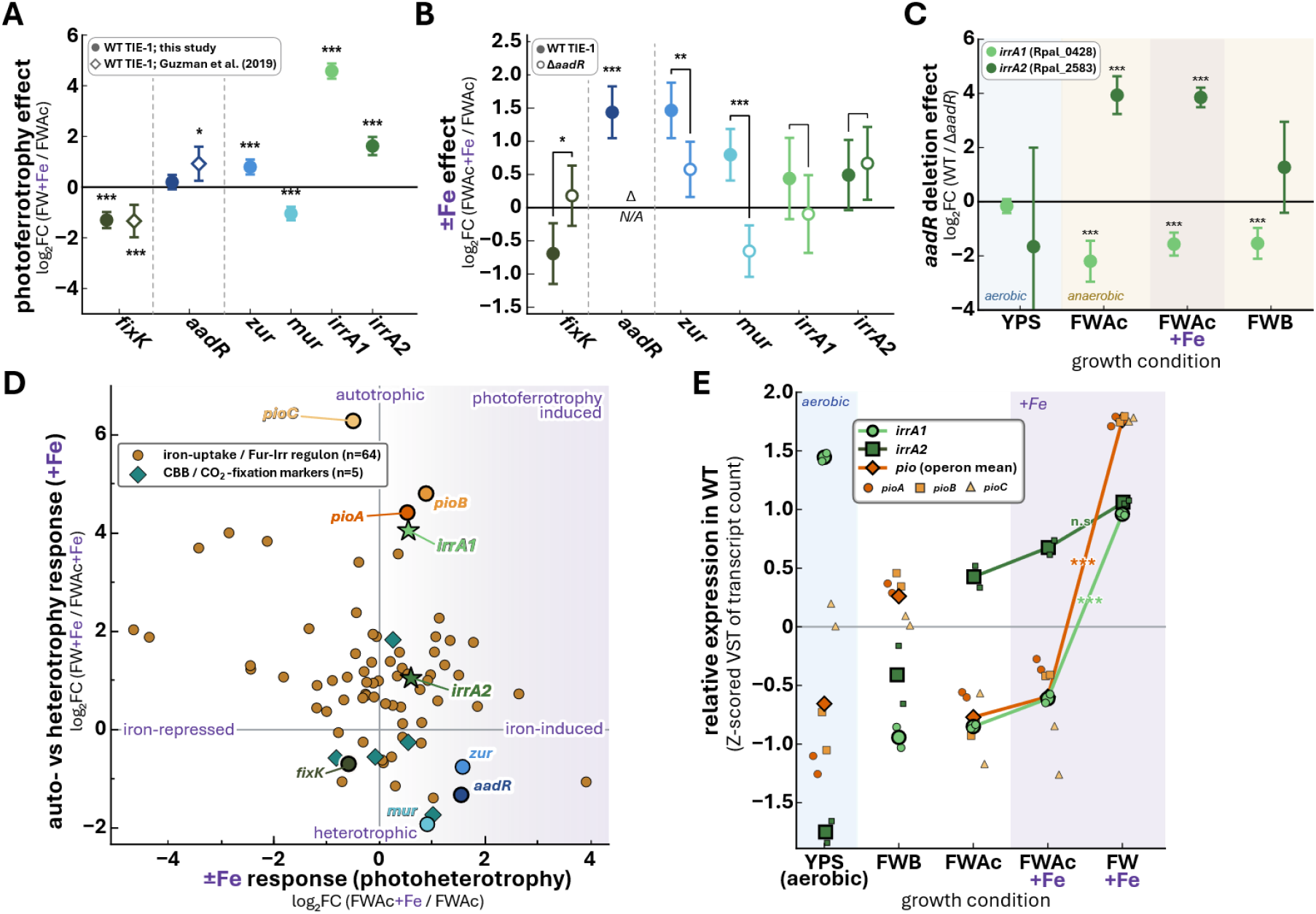
Iron levels mediate FixK–AadR feedback loop and its putative regulatory targets. **(A)** Photoferrotrophy induced repression of *fixK* and activation of *aadR*. Log_2_-adjusted fold change (log_2_FC) between FW+Fe and FWAc is shown for each transcript of interest. Expression data from the same conditional comparison released by Guzman *et al*. (2019; **SI Dataset S02**; open diamonds) were processed alongside this study’s transcriptomic data (filled circles) (45). Stars indicate significance of differential expression (BH-adjusted Wald test, DESeq2). **(B)** Fe(II) significantly shifted expression of the regulators of interest independently of photoferrotrophic metabolism. Furthermore, deletion of *aadR* led to significant shifts in the iron-responsivity of the regulators: induction of *irrA1* by +Fe was ablated in Δ*aadR*, while induction of *mur* became repression of it. Stars indicate a statistically significant difference in ±Fe differential expression between WT and Δ*aadR* (two-way ANOVA). **(C)** Deletion of *aadR* leads to opposite expression patterns in *irrA1* and *irrA2* during anaerobic growth on acetate, regardless of ±Fe state. **(D)** Induction of *irrA1* and *irrA2* is characteristic of a photoferrotrophy-specific response, as their transcription is activated by +Fe and further enhanced by operating under the photoferrotrophic growth mode. Also, the Fe(II)-driven shift in *aadR* and *fixK* transcription occurs independently of photoferrotrophy. Furthermore, *pioABC* transcription is induced specifically by the photoferrotrophic growth mode, as +Fe photoheterotrophy only marginally increased its expression. Data are shown on a scatterplot with quadrants labelled by interpreted qualitative meaning. Background points represent iron-uptake-related genes (circles) predicted to be in the Irr/Fur regulons and CBB-associated genes (diamonds) associated with the autotrophic growth mode.^23,66^ **E**. Expression of both *irrA* paralogues tracks expression of *pioABC* during anaerobic photoheterotrophy. Averages of replicate points are displayed as larger shapes with thicker outlines. Relative expression was calculated by applying a Z-score calculation to variance-stabilised transformation (VST)-normalised RNA-seq data. Stars represent the significance of each transcript’s photoferrotrophy-linked nature (i.e., the difference between expression on FW+Fe relative to on the other conditions).

Transcription of Fur-family regulators changed in Δ*aadR* under anaerobic conditions (**Fig. 2B**). Notably, expression of *irrA2* was significantly greater in wild-type than Δ*aadR*, while *irrA1* showed the opposite pattern, which was replicated across all tested photoheterotrophic growth conditions (**Fig. 2C**). Transcriptomic data for Δ*aadR* on photoferrotrophy were unavailable due to the complete failure of the strain to grow on that condition. The *irrA2* gene shares an upstream intergenic region with the *fez* operon, has predicted tandem FNR sites in its promoter,^66^ and appears to be strongly activated by the FixK–AadR system under anaerobic conditions.^13^,^22^,^28^,^30^,^34^ The promoter of *irrA1* lacks a FNR site but holds a predicted Mur-binding site.^66^ In wild-type TIE-1, *irrA1* showed a striking 28-fold increase in expression under photoferrotrophy relative to photoheterotrophy (log_2_FC = +4.58, *q =* 3.9×10^-170^), far outstripping its response to Fe(II) during heterotrophic +Fe growth (**Fig. 2D, E**). This suggests that *irrA1* transcription responds to the metabolic context of photoferrotrophy rather than to iron concentration alone (as does *pioABC*).

Contrasting the lack of an AadR-mediated iron response in *irr*, iron-responsive differential expression of *mur* was AadR-dependent. In wild-type, *mur* was activated under FWAc+Fe (log_2_FC = +0.76, *q =* 3.2×10^−5^), but Δ*aadR* inverted this response, repressing it (log_2_FC = −0.64, *q =* 0.017). Transcription of *mur* is repressed specifically under photoferrotrophic conditions, or in absence of AadR, further implicating AadR as important for the photoferrotrophic regulatory programme. Activation of *zur* by +Fe occurred in wild-type, but this was significantly attenuated in Δ*aadR* (log_2_FC = +1.46 vs. +0.58; *q* = 0.008; strain * iron interaction; **Fig. 2B**). Whether direct or indirect, these findings position AadR upstream of at least three of the Fur-family transcription factors in a condition-dependent regulatory cascade, though its role for each differs in importance.

### Loss of AadR perturbs the global transcriptome more severely under growth conditions necessitating tighter control over iron-related or iron-requiring pathways

Principal component analysis (PCA) of mutant and wild-type transcriptomes revealed that photoheterotrophy-related gene expression was the primary driver of the differences observed between whole transcriptomes: PC1 explained 30.6% of variance (**Fig. 3A, B**). Iron was nearly as strong a driver, correlating heavily with PC2, which explained a quarter of the variation observed (PC2, 24.5%, [Fe] *r* = +0.89). The displacement of Δ*aadR* from wild-type was largest with benzoate growth (59 units), consistent with AadR’s characterised role regulating aromatic degradation (**Fig. 3C**).^20^ The strain’s second largest displacement was on FWAc+Fe (42 units) predominantly along the PC2 axis, and more than double the displacement shown on FWAc with basal iron (18 units; **Fig. 3C**). The disproportionate iron-related axis displacement suggests a substantial expansion of AadR’s regulatory role.

**Figure 3.**
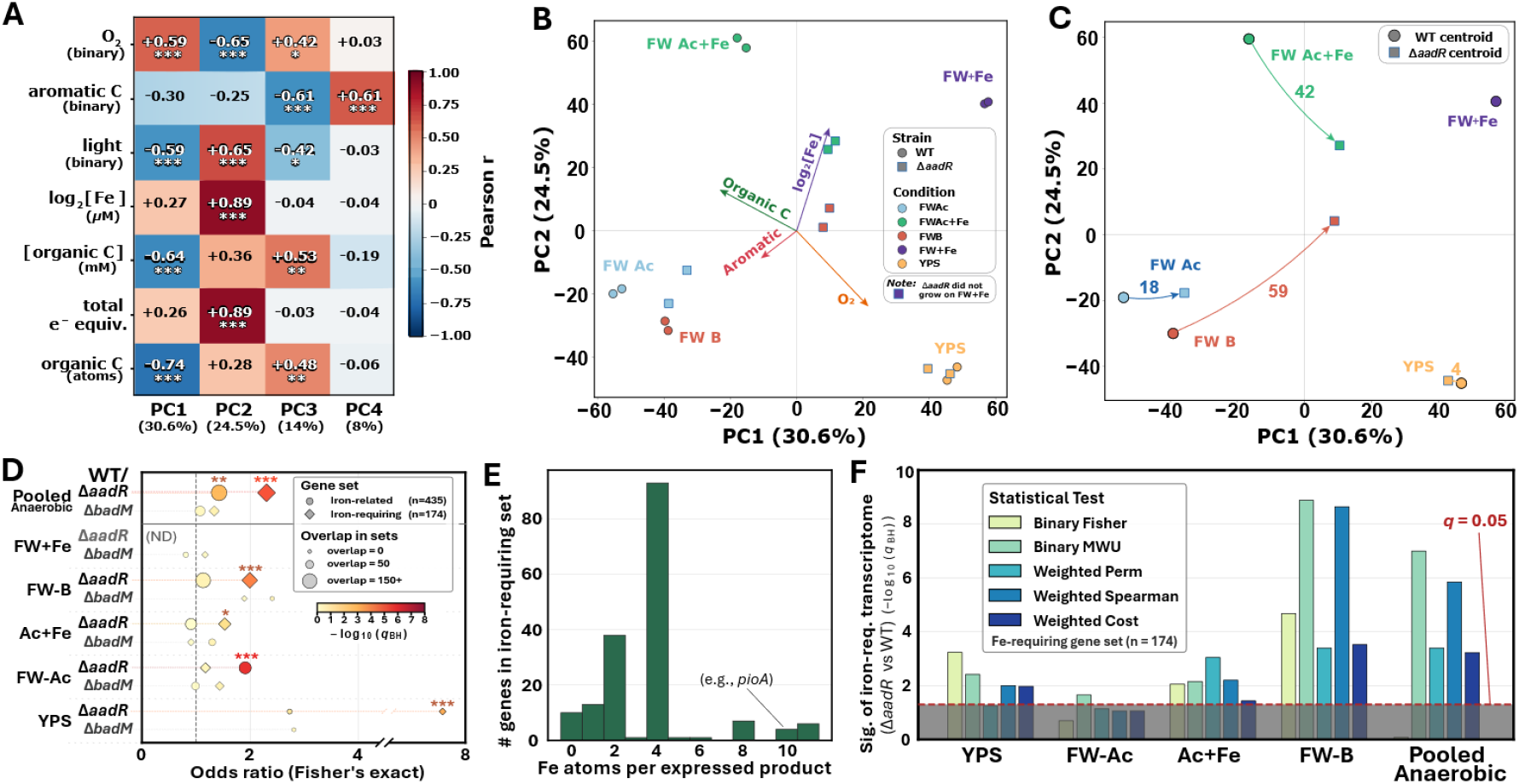
An environmental-driven stimulus hierarchy drives AadR-dependent changes in the global transcriptome, especially for iron-related and iron-requiring genes. **(A)** Correlation matrix, mapping environmental (growth condition) stimulus indices to principal components (PCs). PC1, explaining 30.6% of the variation, correlates positively with oxygen, and negatively with light and amount of organic carbon provided. PC2, explaining 24.5% of the variation, correlates very strongly (r=+0.89, p<0.001) with iron concentration, positively with light, and negatively with oxygen. **(B)** PCA biplot of PC1 vs PC2. Metadata vectors and biological replicate points for VST-adjusted whole transcriptomes are shown. **(C)** PCA displacement plot of PC1 vs PC2, showing how loss of *aadR* shifts the global transcriptome for each growth condition (centroids of replicate point averages are shown). No shift could be calculated for FW+Fe as the Δ*aadR* strain did not grow on that condition. **(D)** Odds ratio of Fisher’s exact test on the effect of knocking out *aadR* (or *badM*) on iron-requiring (*n*=174) and iron-related (*n*=435) DE gene sets, versus all other DE genes. The Δ*aadR* strain differentially regulates iron-related genes on FWAcetate, and iron-requiring genes on FWAc+Fe, FWB, and YPS. Stars mark significance compared to the background of all DE genes between aerobic and anaerobic conditions (log_2_FC > 1, q < 0.05). Bubbles are shaded by significance (–log_10_ *q* value (Benjamini-Hochberg FDR)) and sized by overlap (number of genes of each category in the differentially expressed set). **(E)** Distribution in the number of iron atoms required (by the holoenzyme product) to express each iron-requiring gene. The spike at four iron atoms is the abundance of [4Fe-4S] cluster proteins (**SI Dataset S03**). Genes with zero Fe weight here are ones encoding the subunit of a Fe-incorporating complex. **(F)** Statistical analysis of correlation between being iron-requiring or not (binary tests) or degree of iron requirement (calculated in **E**, weighted tests) and being significantly differentially regulated in Δ*aadR* vs wild-type TIE-1. Full results of the shown binary tests (Fisher and Mann-Whitney U) and weighted tests (PERMANOVA, Spearman’s rank correlation, and weighted average of iron cost) are in **SI Dataset S03**. A paired analysis was performed for comparison with the analytical control Δ*badM* (**Fig. S18**).

We next sought to determine whether AadR’s regulatory sphere of influence is significantly enriched for iron-related pathways. We annotated the TIE-1 protein-coding genome for “iron-related” and “iron-requiring” sets of genes and tested if they were over-represented in the set of differentially expressed (DE) genes between wild-type and Δ*aadR* (**Tables ST8-ST11; SI Dataset S03**).^57–59,67,68^ Genes associated with iron metabolism (such as siderophore production or haem biosynthesis) were classified as iron-related (*n*=435). Those whose products coordinate iron cofactors such as [Fe-S]-clusters or haem groups, (i.e., iron is required for their expression), were classified as iron-requiring (*n*=174). Test comparisons were restricted to genes differentially expressed between aerobic and basal-iron anaerobic conditions, to control for false positive signals caused by many anaerobic genes being iron-requiring. BH-FDR correction was applied across all tests.

The DE gene set was significantly enriched for these sets (pooled Fisher OR = 2.34, *q =* 2.6×10^-6^) (**Fig. 3D**). This enrichment persisted even after controlling for genes that are already DE between aerobic and anaerobic conditions, confirming that the iron-related signal is distinct, rather than just representative of AadR’s known regulation of anaerobic metabolism, which inherently demands iron cofactors to catalyse anoxic biochemical reactions (OR = 2.92, *q =* 2.0×10^-5^). After weighting these genes by the number of iron atoms expended to express them (**Fig. 3E**), iron-requiring genes continued to be over-represented (Mann-Whitney *q =* 1.5×10^- 10^) (**Fig. 3F**). Targets under the regulatory scope of AadR collectively contained an average of 45% more Fe atoms than random chance would predict. As an analysis specificity control, we used a Δ*badM* knockout mutant strain (**Fig. 3D**). Consistent with findings in CGA009, deletion of *badM* in TIE-1 constitutively derepressed the *badDEFG* operon, encoding the iron-requiring benzoyl-CoA reductase (**SI 2, Fig. S10– S13**).^6,8,20^ The set of DE genes between wild-type and Δ*badM* was not significantly enriched for iron-related genes, but it was for iron-requiring genes (*q =* 3×10^-4^). This enrichment became insignificant after excluding the *bad* genes (*q =* 0.16), while Δ*aadR*’s enriched set remained significant when excluding the same genes (**Fig. S18**).

### Characterising the roles of AadR, the P*pio*-BS2 binding site, and regulation of *pioABC*

P*pio-*BS2 likely binds an unknown cognate regulator. A P*pio*-Δ*BS2* reporter showed constitutively reduced expression (∼4-fold average reduction), and the iron-responsive activation of the promoter was abolished.^34^ Paradoxically, reporter activity increased in Δ*fixK* under non-photoferrotrophic conditions, despite the established role of FixK as an activator of P*pioABC*.^34^ Our finding that supplemental Fe(II) decreases *fixK* transcription implies that another factor is involved in the iron-responsive activation of *pioABC*, as the established mechanism of *pioABC* activation by FixK is insufficient to explain the observed pattern.

Wild-type TIE-1 showed modestly greater *pioA* transcription than Δ*aadR* when grown on FWAc (log_2_FC = +1.48, *q =* 2.9×10^−4^), but it showed no significant difference on FWAc+Fe (**Fig. 4A**). On FWB, only *pioC* expression was significantly affected (log_2_FC = −1.59, *q =* 1.1×10^−8^). These modest, inconsistent effects contrast sharply with the ∼4-fold changes in reporter expression in the P*pio*-Δ*BS2* reporter strain or the coordinated 6-18-fold expression changes in Δ*fixK*.^34^ Differences in *pioABC* regulation in Δ*aadR* are instead most parsimoniously explained by altered *fixK* dosage in response to the missing AadR. Transcription of *fixK* was 3- to 30-fold derepressed in Δ*aadR* across anaerobic growth conditions (**Fig. 4A**), which would perturb *pioABC* regulation by FixK via P*pio-*BS1 without requiring direct AadR binding at either site in the *pio* promoter. Supporting this interpretation, iron supplementation during growth on acetate significantly induced *pioA* and *pioB* in Δ*aadR* (log_2_FC = +1.59, +1.39; q < 10^−4^) but not in wild-type (**Fig. 4B**). This potentially reflects the loss of iron-driven repression of *fixK* that we observed in Δ*aadR*, and these data support a lack of direct *pioABC* regulation by AadR at either P*pio* binding site.

**Figure 4.**
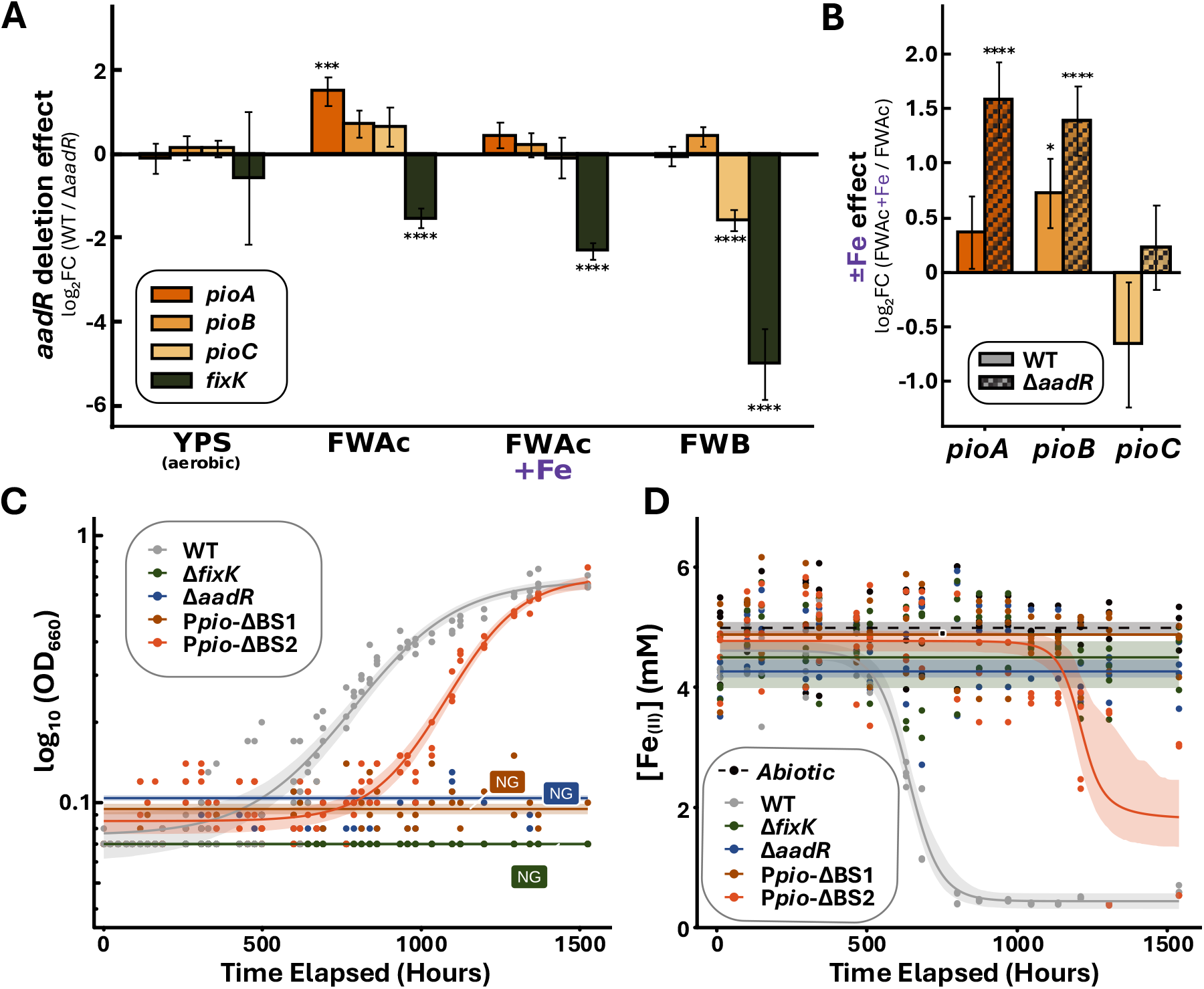
AadR is not a primary, direct regulator of *pioABC* iron-responsive expression. **(A)**. Log_2_ fold change (log_2_FC) in transcript between WT and Δ*aadR*. Relative transcription of *pioA, pioB, pioC*, and *fixK* was compared across four growth conditions: YPS (aerobic), FWAcetate±Fe, and FWBenzoate. Positive values indicate higher expression in wild-type. **(B)** Log2FC in transcript abundance between +Fe and –Fe photoheterotrophic conditions for *pioABC* transcripts, in wild-type (solid bars) and Δ*aadR* (hatched bars). **(C)** Growth curves of wild-type, Δ*aadR* and Δ*fixK* single mutants, and P*pio*-ΔBS1 and ΔBS2 deletion strains on photoferrotrophy (FW+Fe; *n* = 3 biological replicates). Only WT and (after a lag) P*pio*-Δ*BS2* grew. **(D)** Fe(II) oxidation curves of wild-type, single mutants, and P*pio*-ΔBS deletion strains on photoferrotrophy (FW+Fe). Only WT and (after a lag) Ppio-Δ*BS2* oxidised significant amounts of Fe(II). [Fe(II)] was measured using the ferrozine assay. Curves represent logistic growth and decay equations ((inverted) modified Gompertz) fit to replicate datapoints.

To directly assess P*pio*-BS2’s functional role in photoferrotrophy, we grew P*pio*-Δ*BS1* and P*pio*-Δ*BS2* strains on FW+Fe alongside wild-type and regulator mutants (**Fig. 4C**). Δ*BS1* entirely failed to grow, copying the phenotype of Δ*fixK* and confirming BS1 as the essential FixK-dependent activation site. Δ*BS2* grew and measurably oxidised Fe(II), but with substantially delayed onset and less complete oxidation of the provided iron (∼1.75 mM Fe(II) remaining vs. ∼0.25 mM) (**Fig. 4D**). The defective but not abolished photoferrotrophic capability of Δ*BS2* supports the previous finding that BS2 contributes to full *pio* activation under photoferrotrophy without being essential to growth under this condition.^34^

The unknown P*pio*-BS2 factor must satisfy two constraints: it must bind at or near BS2 to regulate *pioABC* expression, and its regulatory effect (or regulation of it) must be iron-responsive. We found that the FixK– AadR negative feedback loop plays a significant role in the regulation of *mur* and of both *irrA* paralogues in response to iron. An Irr-binding motif appears to overlap P*pio-*BS2 in the *pioABC* promoter, suggesting the possibility that a Fur-family regulator might bind at that site (FIMO on a matrix built from 122 Irr-regulated sites from across taxonomic order *Rhizobiales* (i.e., *Hyphomicrobiales*) to which *Rhodopseudomonas* belongs) vs the TIE-1 genome, MEME-suite, *p* = 4.73 × 10^−7^, *q =* 0.032; **Fig. S14**).^66,69,70^

### Components of the oxygen-iron regulatory network architecture are conserved beyond *Rhodopseudomonas palustris*

To place this regulatory architecture into evolutionary context, we mapped iron- and oxygen-responsive regulators across a 50-genome panel spanning the Hyphomicrobiales and select outgroups (**Fig. 5**). The common *α-*proteobacterial ancestor had a single Fur regulator for iron homeostasis, but its function in many descendent lineages (including Hyphomicrobiales) has shifted from sensing and responding to Fe(II) to Mn(II), being reclassified as Mur (**Fig. 5A**).^23^ The Fur/Mur–Irr network was ancient and near-universally conserved within this scope, while a dedicated Fe(II)-sensing RirA was only present within a single derived clade (the branch bearing *Rhizobiaceae* and relatives, eleven genomes; **Fig. 5B**). TIE-1 encodes a single Rrf2-family protein, BadM, which is a distant homologue of RirA and lacks its [Fe-S]-coordinating residues. Homologues of *irr* were classified as an *irrA1* based on two criteria: *irrA1* sits at a conserved ancestral locus next to the *fab* operon, and it carries a haem-regulatory motif (HRM; Cys-Pro-Trp).^28^ The HRM marked the ancestral copy in 12 of 12 GTDB *Xanthobacteraceae* (≈ NCBI *Nitrobacteraceae*) genomes, and it appeared in no genome outside that family (**Fig. S15**).

**Figure 5.**
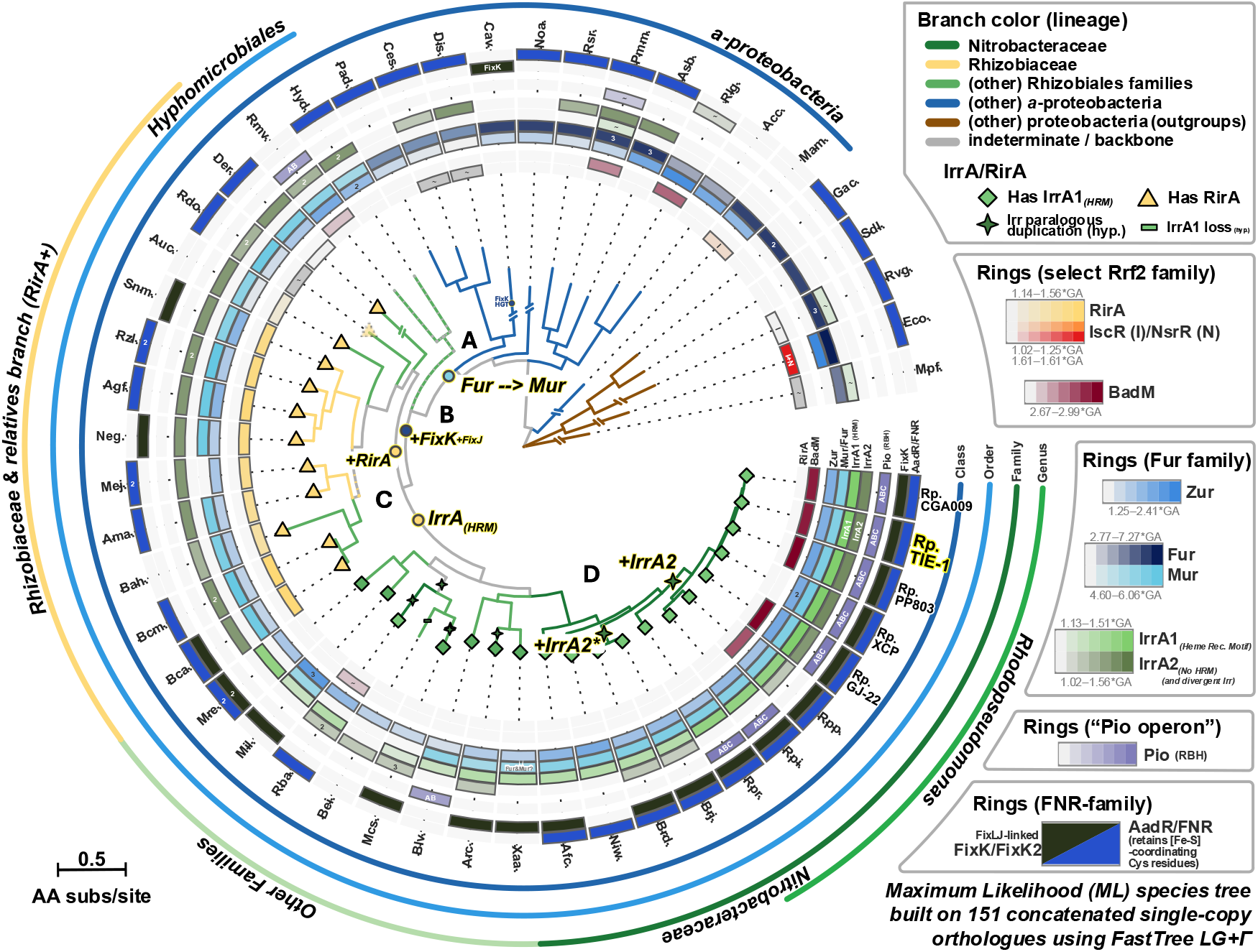
This regulatory network scheme is conserved across alphaproteobacterial lineages. Maximum-likelihood phylogenomic species tree built from 50 α-proteobacterial genomes (from the type strain set of Hördt *et al*., 2020), inferred from 151 concatenated single-copy orthologues using LG+Γ (FastTree); topology was congruent with a GTDB bac120 reference tree.^75–77^ Enclosing metadata rings show, per genome, the presence and HMM bitscore (relative to the family gathering-threshold, “*GA”) of Furfamily (Fur/Mur, Zur, IrrA1/IrrA2), select Rrf2-family (RirA, IscR/NsrR, BadM) and select CRP/FNR-family (FixL(-J)-linked FixK/FixK_2_, AadR/FNR/FixK_1_) regulators, and reciprocal best-hit BLAST (RBH) presence of the *pio* operon. Every claimed absence was confirmed at the nucleotide level by tBLASTn. Outer rings show taxonomic assignment. Tip symbols and branch label symbols are best estimates from tip presence/absence and homologue divergence, and mix inferences drawn from work performed here with findings from the literature. (**A)** A functional shift from the Fe^2+^-sensing by the ancestral Fur toward Mn^2+^/Mur was placed near the divergence of the Hyphomicrobiales, per Rodionov *et al*. (2006).^15^ **(B)** Emergence of the FixK–FixL/J oxygen-response architecture occurred early in the Hyphomicrobiales, with the FixK present in some Caulobacterales due to horizontal transfer from this order, per Tsoy *et al*. (2016).^15^ **(C)** Within the Hyphomicrobiales, the *Rhizobiaceae* lineage carries the Rrf2-family Fe(II) sensor RirA (11 genomes), whereas the *Nitrobacteraceae* and relatives (GTDB: *Xanthobacteraceae*) lineage instead bears a high-affinity haem-regulatory motif in IrrA1 (this study; motif determinants after Todd *et al*. (2006).^28^ **(D)** Independent, lineage-specific duplications of *irr* have frequently occurred across the *Nitrobacteraceae* and beyond. In *R. palustris*, the ancestral *fab*-locus copy ***irrA1*** (*Rpal_0428*, HRM^+^) is distinguished from the derived copy ***irrA2*** (*Rpal_2583*, HRM^−^). An analogous pair (labelled ‘*irrA2\**’) is present in *Bradyrhizobium*, resulting from an independent duplication event. *n* = 50 genomes. Full genome accessions in **Table ST11**; other supporting data in **Dataset S09**.

The gene encoding Irr looks to have undergone several independent gene replication events across Hyphomicrobiales (**Fig. 5C-D)**. Well-supported shallow duplications include the *R. palustris* IrrA2 and the genus-level *Bradyrhizobium* duplicate. IrrA2 is a species-level gain in *Rhodopseudomonas palustris*: other *Rhodopseudomonas* species, such as *R. infernalis* and *R. rhenobacensis*, have only IrrA1 and no apparent artifact or scar of an IrrA2 loss. Irr duplications are not exclusively confined to lineages lacking RirA, but they are widespread among them, making a compelling case for the importance of Irr when lacking RirA.

## Discussion

Our data suggest an expanded role for AadR as a central integrator of oxygen-responsive regulation with iron-responsive metabolism in TIE-1 (**Fig. 6A**). The difference in expression of *fixK* and *aadR* is further shifted by iron but this iron-driven repression of *fixK* is ablated in Δ*aadR* (**Fig. 6B**), supporting the interpretation that the two exist in a balance maintained by a negative feedback loop, rather than a unilateral hierarchy. Like *E. coli* FNR, AadR retains the conserved cysteine residues to coordinate an oxygen-sensing [4Fe-4S]-cluster, making it likely that it does so (**Fig. S16, SI 3**).^14,20^ Expression of iron-requiring and iron-related genes is perturbed with loss of AadR (**Fig. 6C**), and both single mutants show defective or inhibited growth in +Fe phototrophy, indicating an oxygen-iron regulatory network link. Our data also imply presence of an iron-responsive regulator or similar mechanism upstream of the FixK–AadR system, suggesting that this link between the oxygen-responsive and iron-responsive regulatory networks is likely bidirectional.

**Figure 6.**
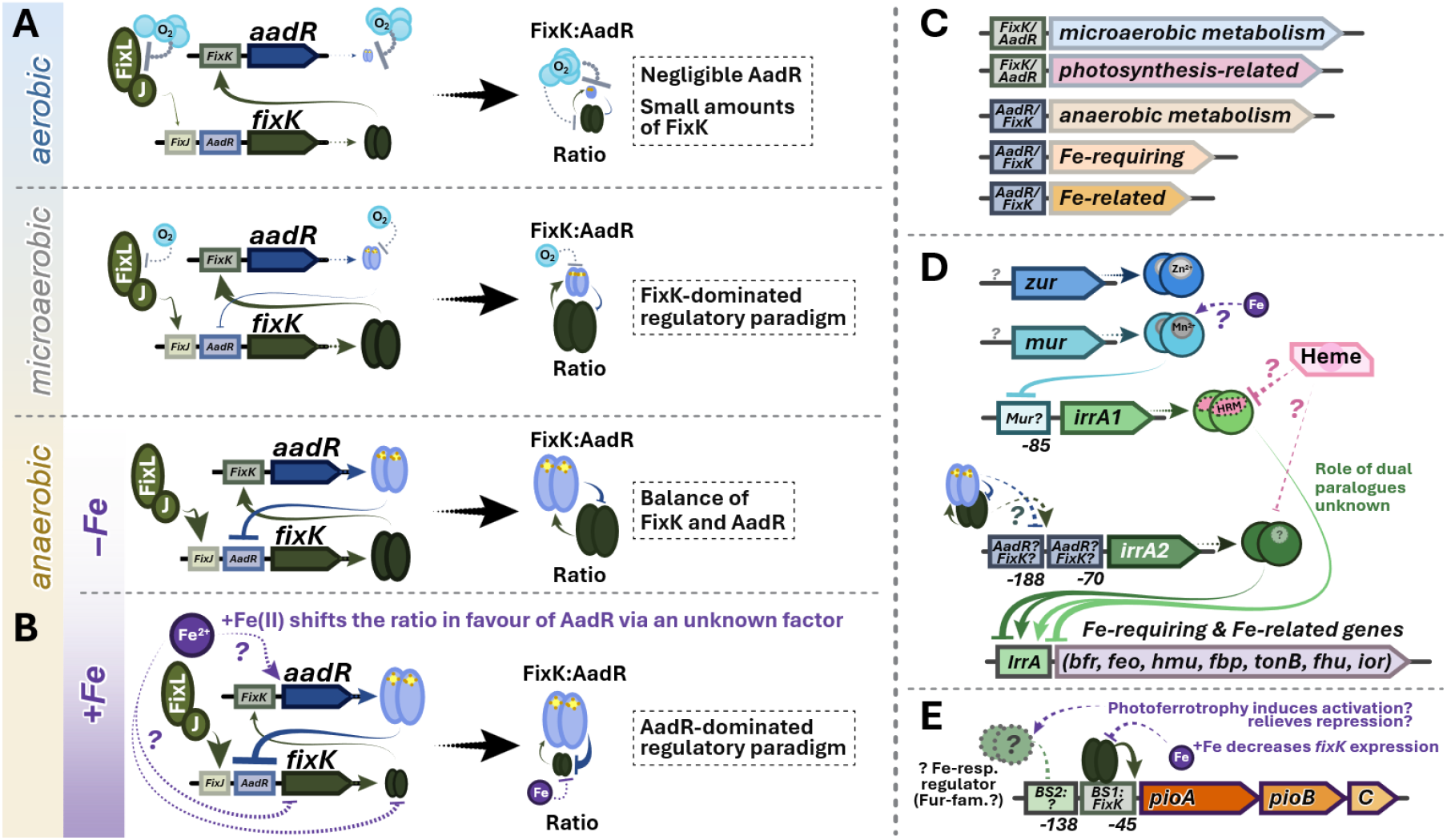
Model of the hypothetical role played by FixK–AadR as integrators of environmental Fe(II) and O_2_ signal response. **(A)** AadR and FixK form a regulatory hierarchy, with the ratio of FixK to AadR increasing and then stabilizing as conditions shift from aerobic to anaerobic.^13^ **(B)** In this work, we extend this framework by discovering that supplementary Fe(II) further shifts the ratio, completing the hierarchical handoff from a FixK-dominated regulatory landscape to one dominated by AadR. **(C)** This leads to the differential regulation of loci including many microaerobic and anaerobic metabolic pathway genes, especially ones that require iron to express or are related to iron metabolism. **(D)** Also under the regulatory scope of this hierarchy are the genes encoding Fur family regulators, the protein products of which then go on to regulate their own regulons. The promoter of *irrA2* contains tandem AadR/FixK/FNR sites, and the *irrA1* promoter has a predicted Mur site.^66^ **(E)** Hypothetical model for *pioABC* operon regulation, given the findings reported in this study. Because +Fe decreases *fixK* expression, reducing activation of *pioABC* via its BS1, the photoferrotrophic growth mode appears to specifically induce expression or activity of a *pioABC* activator or relieve a repressor.

Such links have precedent: in gammaproteobacterium *E. coli*, Fur-mediated iron regulation is shaped by both Fe(II) and oxygen availability, with its anaerobic effects mediated by FNR,^71^ while in gammaproteobacterium *Haemophilus sp*., Fur was shown to repress FNR.^72^ In the divergent *α-*proteobacterium *Cereibacter sphaeroides*, the LysR-family OxyR regulator was demonstrated to activate *fur* in response to oxidative stress, serving as an iron-oxygen regulatory link in that lineage.^73^ A functionally homologous system in *Rhodopseudomonas*, with Fur-family regulated by FixK−AadR and possibly regulating those iron-sensitive regulators in turn, would neatly mirror these findings across a gulf of taxonomic distance. As the proposed conservation of this network architecture is inference based on bioinformatic analysis, experimental validation in related organisms is an important future direction. Our demonstration of FixK−AadR regulation of Fur-family iron-responsive regulators is to our knowledge the first known example proposed in the agriculturally and environmentally relevant clade of Hyphomicrobiales.

We interpret that the novel growth defects in mutant strains were revealed and/or enhanced by the naturomimetic fluorescent lighting, forcing *R. palustris* to operate closer to its metabolic margins and making the regulatory precision of the FixK–AadR feedback loop even more critical (**SI 1.1**). Disrupting this fragile balance carries severe fitness consequences for TIE-1: phototrophy-related genes were repressed in both Δ*fixK* and Δ*aadR*, and the synthetic lethality of the double mutant shows that at least one of FixK or AadR is essential for any anaerobic viability in the wild.^13,34^ Under +Fe conditions, where simultaneous regulation of both iron utilisation and photosystem expression is critical, even the single mutants struggled to grow.

Beyond direct regulation, AadR indirectly regulates many iron-related or iron-requiring loci through mediating expression of Fur-family regulators. In the Δ*aadR* strain, +Fe suppressed *mur* transcription instead of activating it. Mur is the closest homolog to ancestral Fur in *R. palustris*, and it is predicted to directly regulate *irrA1* (**Fig. 6D**).^30,66^ Mur is a Mn-responsive regulator in the closely-related *Bradyrhizobium* that has been shown to become functionally active *in vitro* with Fe(II) replacing Mn(II) as the effector, raising the possibility that *R. palustris* Mur can directly sense iron.^24,31,74^ The photoferrotrophy-specific *irrA1* induction suggests its regulation integrates information on metabolic state beyond iron concentration. Recent work in *R. palustris* strain CGA009 has suggested that the RegSR sensor-regulator system may play a role in regulation of *pioABC* depending on cellular redox state, and it is possibly relevant here as well. In *Bradyrhizobium*, IrrA1 is degraded at the protein level after binding haem,^14^ offering the possibility that the high *irrA1* transcript levels during photoferrotrophy may reflect compensatory transcription in response to rapid protein turnover, rather than greater abundance of IrrA1 protein product. An orthogonal approach such as proteomics may help future studies further refine the list of plausible direct *pioABC* regulator candidates.

Core transcriptomic findings are supported by large effect sizes and *q*-values well below 0.001, and they are further corroborated by consistency across replicates and conditions. Comparison with the independent Guzman *et al*. (2019) and Rey and Harwood (2010) datasets grants supports that the data reflect meaningful differences in AadR-dependent and iron-dependent regulation rather than analytical or methodological bias.^13,65^ AadR’s effects may be mediated wholly or in part by altered dosage of FixK or another subordinate regulator; transcriptomic data alone cannot easily distinguish between direct and indirect relationships. Mechanism notwithstanding, we confirm numerous novel regulatory relationships in a network mediated by AadR and iron. Should AadR’s predicted [4Fe-4S] cluster exist, its oxygen sensitivity would pose a formidable barrier to future *in vitro* work on the protein.^21^ Such work would demand considerable anaerobic protein purification expertise and equipment, but it would help confirm the protein structure and resolve direct binding sites.^16^ Future work should also prioritise examining Fur-family regulator deletion mutants, which could help clarify the precise structure of this intermeshing regulatory network, and it also may help elucidate the mechanism(s) for the effect of Fe(II) on the balance between AadR and FixK.

## Acknowledgements

We thank undergraduate researchers L. Dong and D. Cuthbert for assistance conducting growth experiments, and undergraduate researchers M. Silberman and T. Vu as well as PhD candidate H. Morrison for their support. The following grants to A. Bose supported this work: National Science Foundation (Grant Numbers 2021822, 2124088, 2117198, and 2300081), an NIGMS grant (NIH R01 GM141344), a DEPSCoR grant (FA9550-21-1-0211), and a David & Lucile Packard Foundation Fellowship.

## References

1. Larimer, F. W. et al. Complete genome sequence of the metabolically versatile photosynthetic bacterium Rhodopseudomonas palustris. Nat Biotechnol 22, 55–61 (2004).

2. van Niel, C. B. [1] Techniques for the enrichment, isolation, and maintenance of the photosynthetic bacteria. in Methods in Enzymology vol. 23 3–28 (Academic Press, 1971).

3. Oda, Y. et al. Genotypic and Phenotypic Diversity within Species of Purple Nonsulfur Bacteria Isolated from Aquatic Sediments. Applied and Environmental Microbiology 68, 3467–3477 (2002).

4. Jiao, Y., Kappler, A., Croal, L. & Newman, D. Isolation and Characterization of a Genetically Tractable Photoautotrophic Fe(II)-Oxidizing Bacterium, Rhodopseudomonas palustris Strain TIE-Applied and environmental microbiology 71, 4487–96 (2005).

5. Adessi, A. et al. Draft genome sequence and overview of the purple non sulfur bacterium Rhodopseudomonas palustris 42OL. Stand in Genomic Sci 11, 24 (2016).

6. VanDrisse, C. M. & Escalante-Semerena, J. C. Small-Molecule Acetylation Controls the Degradation of Benzoate and Photosynthesis in Rhodopseudomonas palustris. mBio 9, e01895–18 (2018).

7. Kathol, M. et al. High enzyme promiscuity in lignin degradation mechanisms in Rhodopseudomonas palustris CGA009. Applied and Environmental Microbiology 0, e00573–25 (2025).

8. Hirakawa, H., Hirakawa, Y., Greenberg, E. P. & Harwood, C. S. BadR and BadM Proteins Transcriptionally Regulate Two Operons Needed for Anaerobic Benzoate Degradation by Rhodopseudomonas palustris. Appl Environ Microbiol 81, 4253–4262 (2015).

9. Morrison, H. M. & Bose, A. Purple non-sulfur bacteria for biotechnological applications. Journal of Industrial Microbiology and Biotechnology 52, kuae052 (2025).

10. Poulain, A. J. & Newman, D. K. Rhodobacter capsulatus Catalyzes Light-Dependent Fe(II) Oxidation under Anaerobic Conditions as a Potential Detoxification Mechanism. Applied and Environmental Microbiology 75, 6639–6646 (2009).

11. Nikeleit, V. et al. Phototrophic Fe(II) oxidation by Rhodopseudomonas palustris TIE-1 in organic and Fe(II)-rich conditions. Environmental Microbiology 26, e16608 (2024).

12. Melton, E. D., Schmidt, C., Behrens, S., Schink, B. & Kappler, A. Metabolic Flexibility and Substrate Preference by the Fe(II)-Oxidizing Purple Non-Sulphur Bacterium Rhodopseudomonas palustris Strain TIE-1. Geomicrobiology Journal 31, 835–843 (2014).

13. Rey, F. E. & Harwood, C. S. FixK, a global regulator of microaerobic growth, controls photosynthesis in Rhodopseudomonas palustris. Molecular Microbiology 75, 1007–1020 (2010).

14. Fleischhacker, A. S. & Kiley, P. J. Iron-containing transcription factors and their roles as sensors. Current Opinion in Chemical Biology 15, 335–341 (2011).

15. Tsoy, O. V., Ravcheev, D. A., Čuklina, J. & Gelfand, M. S. Nitrogen Fixation and Molecular Oxygen: Comparative Genomic Reconstruction of Transcription Regulation in Alphaproteobacteria. Front Microbiol 7, 1343 (2016).

16. Myers, K. S. et al. Genome-scale Analysis of Escherichia coli FNR Reveals Complex Features of Transcription Factor Binding. PLoS Genet 9, e1003565 (2013).

17. Nellen-Anthamatten, D. et al. Bradyrhizobium japonicum FixK2, a Crucial Distributor in the FixLJ-Dependent Regulatory Cascade for Control of Genes Inducible by Low Oxygen Levels. Journal of Bacteriology 180, 5251–5255 (1998).

18. Mesa, S. et al. Comprehensive Assessment of the Regulons Controlled by the FixLJ-FixK2-FixK1 Cascade in Bradyrhizobium japonicum. Journal of Bacteriology 190, 6568–6579 (2008).

19. Bonnet, M. et al. The Structure of Bradyrhizobium japonicum Transcription Factor FixK2 Unveils Sites of DNA Binding and Oxidation. J Biol Chem 288, 14238–14246 (2013).

20. Dispensa, M. et al. Anaerobic growth of Rhodopseudomonas palustris on 4-hydroxybenzoate is dependent on AadR, a member of the cyclic AMP receptor protein family of transcriptional regulators. J Bacteriol 174, 5803–5813 (1992).

21. Egland, P. G. & Harwood, C. S. BadR, a New MarR Family Member, Regulates Anaerobic Benzoate Degradation by Rhodopseudomonas palustris in Concert with AadR, an Fnr Family Member. J Bacteriol 181, 2102–2109 (1999).

22. Grant, C. R. et al. Distinct gene clusters drive formation of ferrosome organelles in bacteria. Nature 606, 160–164 (2022).

23. Rodionov, D. A., Gelfand, M. S., Todd, J. D., Curson, A. R. J. & Johnston, A. W. B. Computational Reconstruction of Iron-and Manganese-Responsive Transcriptional Networks in α-Proteobacteria. PLOS Computational Biology 2, e163 (2006).

24. Singh, R., Ranaivoarisoa, T. O., Gupta, D., Bai, W. & Bose, A. Genetic Redundancy in Iron and Manganese Transport in the Metabolically Versatile Bacterium Rhodopseudomonas palustris TIE-Applied and Environmental Microbiology 86, e01057–20 (2020).

25. Rudolph, G., Hennecke, H. & Fischer, H.-M. Beyond the Fur paradigm: iron-controlled gene expression in rhizobia. FEMS Microbiology Reviews 30, 631–648 (2006).

26. O’Brian, M. R. Perception and Homeostatic Control of Iron in the Rhizobia and Related Bacteria. Annu. Rev. Microbiol. 69, 229–245 (2015).

27. Hamza, I., Hassett, R. & O’Brian, M. R. Identification of a Functional fur Gene in Bradyrhizobium japonicum. Journal of Bacteriology 181, 5843–5846 (1999).

28. Todd, J. D., Sawers, G., Rodionov, D. A. & Johnston, A. W. B. The Rhizobium leguminosarum regulator IrrA affects the transcription of a wide range of genes in response to Fe availability. Mol Genet Genomics 275, 564–577 (2006).

29. Sevilla, E., Bes, M. T., Peleato, M. L. & Fillat, M. F. Fur-like proteins: Beyond the ferric uptake regulator (Fur) paralog. Archives of Biochemistry and Biophysics 701, 108770 (2021).

30. Johnston, A. W. B. et al. Living without Fur: the subtlety and complexity of iron-responsive gene regulation in the symbiotic bacterium Rhizobium and other α-proteobacteria. Biometals 20, 501–511 (2007).

31. Fabiano, E. & O’Brian, M. R. Mechanisms and Regulation of Iron Homeostasis in the Rhizobia. in Molecular Aspects of Iron Metabolism in Pathogenic and Symbiotic Plant-Microbe Associations (eds Expert, D. & O’Brian, M. R.) 41–86 (Springer Netherlands, Dordrecht, 2012). doi:10.1007/978-94-007-5267-2_3.

32. Hamza, I., Qi, Z., King, N. D. & O’Brian, M. R. Fur-independent regulation of iron metabolism by Irr in Bradyrhizobium japonicum. Microbiology (Reading) 146 (Pt 3), 669–676 (2000).

33. Ehrenreich, A. & Widdel, F. Anaerobic oxidation of ferrous iron by purple bacteria, a new type of phototrophic metabolism. Appl Environ Microbiol 60, 4517–4526 (1994).

34. Bose, A. & Newman, D. K. Regulation of the phototrophic iron oxidation (pio) genes in Rhodopseudomonas palustris TIE-1 is mediated by the global regulator, FixK. Molecular Microbiology 79, 63–75 (2011).

35. Gupta, D. et al. Photoferrotrophs Produce a PioAB Electron Conduit for Extracellular Electron Uptake. mBio 10, e02668–19 (2019).

36. Jiao, Y. & Newman, D. K. The pio Operon Is Essential for Phototrophic Fe(II) Oxidation in Rhodopseudomonas palustris TIE-1. Journal of Bacteriology 189, 1765–1773 (2007).

37. Haas, N. W. et al. PioABC-Dependent Fe(II) Oxidation during Photoheterotrophic Growth on an Oxidized Carbon Substrate Increases Growth Yield. Applied and Environmental Microbiology 88, e00974–22 (2022).

38. Wan, X.-F. et al. Transcriptomic and proteomic characterization of the Fur modulon in the metal-reducing bacterium Shewanella oneidensis. J Bacteriol 186, 8385–8400 (2004).

39. Kouzuma, A., Kasai, T., Hirose, A. & Watanabe, K. Catabolic and regulatory systems in Shewanella oneidensis MR-1 involved in electricity generation in microbial fuel cells. Front. Microbiol. 6, (2015).

40. Jiao, Y., Kappler, A., Croal, L. R. & Newman, D. K. Isolation and Characterization of a Genetically Tractable Photoautotrophic Fe(II)-Oxidizing Bacterium, Rhodopseudomonas palustris Strain TIE-Appl Environ Microbiol 71, 4487–4496 (2005).

41. Bai, W., Ranaivoarisoa, T. O., Singh, R., Rengasamy, K. & Bose, A. n-Butanol production by Rhodopseudomonas palustris TIE-1. Commun Biol 4, 1–16 (2021).

42. Quandt, J. & Hynes, M. F. Versatile suicide vectors which allow direct selection for gene replacement in gram-negative bacteria. Gene 127, 15–21 (1993).

43. Wetmore, K. M. et al. Rapid Quantification of Mutant Fitness in Diverse Bacteria by Sequencing Randomly Bar-Coded Transposons. mBio 6, 10.1128/mbio.00306-15 (2015).

44. Pechter, K. B., Gallagher, L., Pyles, H., Manoil, C. S. & Harwood, C. S. Essential Genome of the Metabolically Versatile Alphaproteobacterium Rhodopseudomonas palustris. J Bacteriol 198, 867–876 (2016).

45. Khan, S. R., Gaines, J., Roop, R. M. & Farrand, S. K. Broad-Host-Range Expression Vectors with Tightly Regulated Promoters and Their Use To Examine the Influence of TraR and TraM Expression on Ti Plasmid Quorum Sensing. Appl Environ Microbiol 74, 5053–5062 (2008).

46. Dehio, C. & Meyer, M. Maintenance of broad-host-range incompatibility group P and group Q plasmids and transposition of Tn5 in Bartonella henselae following conjugal plasmid transfer from Escherichia coli. Journal of Bacteriology 179, 538–540 (1997).

47. Widdel, F. & Bak, F. Gram-Negative Mesophilic Sulfate-Reducing Bacteria. The Prokaryotes 10.1007/978-1-4757-2191-1_21 (1992) doi:10.1007/978-1-4757-2191-1_21.

48. Zwietering, M. H., Jongenburger, I., Rombouts, F. M. & van ‘t Riet, K. Modeling of the Bacterial Growth Curve. Applied and Environmental Microbiology 56, 1875–1881 (1990).

49. Stookey, L. L. Ferrozine---a new spectrophotometric reagent for iron. Anal. Chem. 42, 779–781 (1970).

50. Love, M. I., Huber, W. & Anders, S. Moderated estimation of fold change and dispersion for RNA-seq data with DESeq2. Genome Biol 15, 550 (2014).

51. Arkin, A. P. et al. KBase: The United States Department of Energy Systems Biology Knowledgebase. Nat Biotechnol 36, 566–569 (2018).

52. Schurch, N. J. et al. How many biological replicates are needed in an RNA-seq experiment and which differential expression tool should you use? RNA 22, 839–851 (2016).

53. Coenye, T. Do results obtained with RNA-sequencing require independent verification? Biofilm 3, 100043 (2021).

54. Muzellec, B., Teleńczuk, M., Cabeli, V. & Andreux, M. PyDESeq2: a python package for bulk RNA-seq differential expression analysis. Bioinformatics 39, btad547 (2023).

55. Pedregosa, F. et al. Scikit-learn: Machine learning in Python. Journal of machine learning research 12, 2825–2830 (2011).

56. Virtanen, P. et al. SciPy 1.0: fundamental algorithms for scientific computing in Python. Nat Methods 17, 261–272 (2020).

57. Garber, A. I. et al. FeGenie: A Comprehensive Tool for the Identification of Iron Genes and Iron Gene Neighborhoods in Genome and Metagenome Assemblies. Front Microbiol 11, 37 (2020).

58. Potter, S. C. et al. HMMER web server: 2018 update. Nucleic Acids Res 46, W200–W204 (2018).

59. Valasatava, Y., Rosato, A., Banci, L. & Andreini, C. MetalPredator: a web server to predict iron–sulfur cluster binding proteomes. Bioinformatics 32, 2850–2852 (2016).

60. Kuo, F.-S.Chien, Y.-H. & Chen, C.-J. Effects of light sources on growth and carotenoid content of photosynthetic bacteria Rhodopseudomonas palustris. Bioresource Technology 113, 315–318 (2012).

61. Ritchie, R. J. The Use of Solar Radiation by the Photosynthetic Bacterium, Rhodopseudomonas palustris: Model Simulation of Conditions Found in a Shallow Pond or a Flatbed Reactor. Photochemistry and Photobiology 89, 1143–1162 (2013).

62. Diáková, K., Holcová, V., Šíma, J. & Dušek, J. The Distribution of Iron Oxidation States in a Constructed Wetland as an Indicator of Its Redox Properties. Chemistry & Biodiversity 3, 1288–1300 (2006).

63. Hädrich, A. et al. Microbial Fe(II) oxidation by Sideroxydans lithotrophicus ES-1 in the presence of Schlöppnerbrunnen fen-derived humic acids. FEMS Microbiol Ecol 95, (2019).

64. Hädrich, A., Heuer, V. B., Herrmann, M.Hinrichs, K.-U. & Küsel, K. Origin and fate of acetate in an acidic fen. FEMS Microbiology Ecology 81, 339–354 (2012).

65. Guzman, M. S. et al. Phototrophic extracellular electron uptake is linked to carbon dioxide fixation in the bacterium Rhodopseudomonas palustris. Nat Commun 10, 1355 (2019).

66. Novichkov, P. S. et al. RegPrecise 3.0 – A resource for genome-scale exploration of transcriptional regulation in bacteria. BMC Genomics 14, 745 (2013).

67. Imlay, J. A. Iron-sulphur clusters and the problem with oxygen. Molecular Microbiology 59, 1073–1082 (2006).

68. Arai, H., Roh, J. H. & Kaplan, S. Transcriptome Dynamics during the Transition from Anaerobic Photosynthesis to Aerobic Respiration in Rhodobacter sphaeroides 2.4.1. Journal of Bacteriology 190, 286–299 (2008).

69. Bailey, T. L., Johnson, J., Grant, C. E. & Noble, W. S. The MEME Suite. Nucleic Acids Res 43, W39–W49 (2015).

70. Grant, C. E., Bailey, T. L. & Noble, W. S. FIMO: scanning for occurrences of a given motif. Bioinformatics 27, 1017–1018 (2011).

71. Beauchene, N. A. et al. O2 availability impacts iron homeostasis in Escherichia coli. Proc Natl Acad Sci U S A 114, 12261–12266 (2017).

72. Gil-Campillo, C. et al. Epigenetic control of the ferric uptake regulator (Fur) and fumarate nitrate reductase (FNR) master regulatory proteins contributes to Haemophilus influenzae survival during lung infection. mBio 16, e01355–25 (2025).

73. Remes, B., Berghoff, B. A., Förstner, K. U. & Klug, G. Role of oxygen and the OxyR protein in the response to iron limitation in Rhodobacter sphaeroides. BMC Genomics 15, 794 (2014).

74. Amato, G. E. et al. A previously uncharacterized Fur-family metalloregulator integrates iron- and manganese-sensing to control virulence gene regulatory networks in Brucella. Sci Rep 15, 44910 (2025).

75. Hördt, A. et al. Analysis of 1,000+ Type-Strain Genomes Substantially Improves Taxonomic Classification of Alphaproteobacteria. Front Microbiol 11, 468 (2020).

76. Parks, D. H. et al. GTDB: an ongoing census of bacterial and archaeal diversity through a phylogenetically consistent, rank normalized and complete genome-based taxonomy. Nucleic Acids Research 50, D785–D794 (2022).

77. Price, M. N., Dehal, P. S. & Arkin, A. P. FastTree 2–approximately maximum-likelihood trees for large alignments. PloS one 5, e9490 (2010).

