## Supplemental Materials for "Naturomimetic conditions expose an unusual oxygen–iron regulatory network in *Rhodopseudomonas* and related *α-proteobacteria*"


### Author Contributions

B.G., T.R., P.P., D.G., J.K., and A.B. designed research; B.G., T.R., P.P., J.L., A.R., D.G., and J.K. performed research; B.G., P.P., J.L., and A.R. analysed the data presented here; B.G. and A.B. wrote the paper. All authors reviewed and approved the final manuscript.


### Mailing Address

Arpita Bose
Department of Biology; Washington University in St. Louis
1 Brookings Drive,
St. Louis MO 63130,
United States of America



### Preprints and Other Versions

A version of this manuscript was deposited to bioRxiv as a pre-print on 29 June 2026: https://doi.org/10.64898/2026.06.27.734994, with the text made available under a CC-BY-NC-ND 4.0 International license.

Copyright is retained by the author(s) and funder(s).


### Keywords

FixK–AadR; Fur-family; Irr; negative feedback loop; *Rhodopseudomonas palustris*; Rhizobiales; Hyphomicrobiales; Iron-responsive regulation

**Supporting Information Text**

**Supplementary Section SI 1: The naturomimetic growth system was designed to better reflect conditions experienced by environmental microorganisms than conventional laboratory growth.**

SI 1.1. Meta-analysis of light used for *R. palustris* growth informed our naturomimetic system:

A broader open question regarding the regulation of metabolic responses in *Rhodopseudomonas palustris* is which environmental signals the bacterium responds to, and how these environmental growth conditions affect the kinetics of its growth. Because of its metabolic versatility, *R. palustris* can survive in a staggering diversity of environments, but it needs regulation to temper this versatility to ensure it is using the most thermodynamically optimal growth mode. Conditions experienced by a microbe in nature are far more complex than those commonly used in laboratory culture, where minimising variability between experiments is key to ensure informative statistics.

As both the spectrum and intensity of light lead to meaningful regulation and phenotypic differences in *R. palustris* under phototrophic growth,[1,2] a decision as to which light source to employ here was necessary. A wide variety of photosynthetic pigments (bacteriochlorophylls and carotenoids) with different absorption wavelengths may be expressed by *R. palustris* (**Fig. S3**).[2] *R. palustris* responds to low light levels by increasing expression of light-harvesting complexes, but the different spectral bands output by different types of light source induce expression of different pigments.[2] A great diversity of light sources have been employed in *R. palustris* studies, with little standardisation or consensus concerning reported bulb type or total irradiance.[1–7] In a study of solar irradiance available to suspended *Rhodopseudomonas* cells in an outdoor pond/photobioreactor (performed by Ritchie (2013)), short-wavelength blue-violet (~350-450 nm) and long near-infrared (762-925 nm) light are quickly depleted at depths below the surface of the water.[8] In a model of a pond with a dense *Rhodopseudomonas* suspension, the attenuated light that reaches cells deeper in a turbid wetland water column consists of a main peak at 674 nm (red light), with a smaller secondary peak observable at ~550 nm (greenish-yellow).[8] However, in a real wetland rather than a pure bacterial suspension, additional spectral filtering occurs. CDOM (chromophoric/coloured dissolved organic matter, made up of humic and fulvic acids from the decomposing vegetable matter common to wetlands) absorbs shorter visible wavelengths, yellow-shifting the available light (**Fig. S3A,B**).[9–11] The chlorophyll of planktonic green photosynthesisers, like algae, absorbs in the ~435 nm (blue) and ~680 nm (red) wavelengths, together tending to leave a green-yellow transmission window.[9–11] This makes the incandescent bulb (commonly used in laboratories for its broad spectrum and high intensity across the infrared band) a less accurate analogue for the conditions experienced by *Rhodopseudomonas* cells in natural field conditions (**Fig. S3C,D; Fig. S9**).

Because our question is most particularly concerned with the life of a microbe in nature, we instead chose to perform all our phototrophic growth experiments under fluorescent lighting. Fluorescent lights match natural wetland conditions in low intensity (consistent with attenuated aquatic light), negligible near-infrared, and a yellow-shifted spectrum with a major peak at ~570 nm (greenish yellow) and minor peak at ~445 nm (blue) (**Fig. S3C,D**). Kuo *et al.* (2012) found that while fluorescent light yielded the second-greatest optical density and cell count (behind incandescent and blue LED), cells grown under it yielded the smallest amount of carotenoids for any light source they tested.[2] While not identical to the available irradiance modelled in the bacterial pond,[8] it is consistent with the light available to cells living within a wetland (especially a waterlogged soil wetland such as a peat bog or fen) or outdoor photobioreactor with modest amounts of CDOM and phytoplankton,[11] precisely the sorts of environments from which we frequently isolate *Rhodopseudomonas palustris*.

SI 1.2. Oxygen concentrations and the oxidation/bioavailability of iron are intrinsically linked:

Gradients of iron and oxygen are inseparably linked in natural environments, and their chemical interplay has shaped microbial evolution for billions of years.[12,13] Regulation of iron homeostasis is essential for microbes, but the concentration and oxidation state of iron vary greatly in natural and constructed wetlands. Iron is the fourth most abundant element in the Earth's crust, and it plays numerous critical roles in biological systems (e.g., as an enzyme cofactor).[14] Importantly, when not bound up in a haem group, environmental iron is found in either the Fe(II) (ferrous) or Fe(III) (ferric) state, with Fe(II) the more soluble and bioavailable of the two by a wide margin.[15] Iron, especially Fe(III), must be chelated by organic molecules to be truly accessible by most forms of life.[16] These chelators, including the siderophores produced by microorganisms, bind and solubilise iron ions allowing their uptake, and have been found to be ubiquitous in soil microbiomes.[17] Fe(II) is swiftly oxidised by oxygen, when present, and therefore is only very abundant in anaerobic microenvironments, such as water-saturated wetland sediments.[18,19] These environments frequently display substantially greater Fe(II) concentrations than aerobic ones. Fens, for example, are groundwater-fed wetlands which may regularly display millimolar-scale Fe(II) concentrations.[19,20] To emulate such environments, for our "iron-supplemented" conditions, we added 5 millimolar ferrous chloride ($FeCl_2$) with nitrilotriacetic acid (NTA) as a chelator.

The ability to break down NTA via anaerobic metabolism is unlikely,[21] and past experiments in TIE-1 showed minimal difference in growth or transcriptional response to Fe(II) with or without NTA.[3] In addition to allowing the persistence of Fe(II), the anaerobic conditions in these wetlands also greatly slow the degradation of recalcitrant organic molecules, especially heavily aromatic ones such as lignin and its degradation intermediates, monolignols.[5,22] The resident microbiota have adapted to the anaerobic conditions, high levels of bioavailable iron, and aromatic carbon substrates by encoding an arsenal of metabolic pathways to take advantage of these environments. The anaerobic breakdown of aromatic carbon by TIE-1 and other strains takes advantage of the abundance of iron by incorporating [Fe-S] clusters as key cofactors of anaerobic enzymes.[12,23]

### SI 1.3. +Fe mixotrophy permits the study of iron oxidation by reluctant photoferrotrophs:

The ability to thrive in diverse microenvironments depends on a microbe's ability to effectively regulate its competing metabolic processes to leverage its limited stocks of cellular energy and reducing power.[24] When multiple possible metabolic pathways are simultaneously available to an organism, the microbe is said to be under mixotrophic conditions. Mixotrophy is thought to be the default metabolic condition presented by natural environments to microorganisms— especially ones so metabolically versatile as *R. palustris*,[6] but photoferrotrophic mixotrophy (growth with an organic carbon source and supplemental Fe(II)) has not commonly been studied in *R. palustris* until relatively recently.[6] An important (and recently growing) body of work has been performed to examine the growth of *R. palustris* and related microorganisms mixotrophically, on either multiple carbon sources, or a carbon source and a separate electron source such as hydrogen or Fe(II).[6,25–29]

Ehrenreich and Widdel (1994) were the first to show that a strain of a photoferrotrophic bacterium (*Rhodobacter*) first consumed a provided organic carbon source (acetate) before shifting its metabolism to begin Fe(II) oxidation.[30] Later work by Melton *et al.* (2014) showed that TIE-1 exhibits the same 'preference', consuming the acetate in its growth medium before engaging in photoferrotrophy,[29] and work by Kopf and Newman (2012) showed that a strain of *Rhodobacter* incapable of strict photoferrotrophy was able to grow and oxidise iron mixotrophically.[31] More recently, Nikeleit *et al.* (2024) showed that when grown with 2 mM Fe(II) and 0.6 mM of an organic carbon source, *R. palustris* TIE-1 oxidised the Fe(II) during growth on one set of carbon sources (e.g., lactate) but oxidised the Fe(II) only *after* consuming other carbon sources (e.g., acetate).[27] Most directly relevant to the present work, Haas *et al.* (2022) showed that *R. palustris* CGA010 grown in media with Fe(II) and either acetate or malate, cells growing on the more oxidised carbon source malate oxidised significantly more iron than those grown with the more reduced carbon source acetate, which did not oxidise a significant amount of iron relative to the abiotic control, a difference that they attribute to the redox state sensing two-component regulator RegSR.[6] Ultimately, that photoferrotrophic mixotrophy may permit the study of strains (such as CGA010) that are unable to effectively grow autotrophically via photoferrotrophy is a benefit of the method, one which we leverage in the present study to examine the effect of supplementary Fe(II) on the transcriptomes and Fe(II) oxidation kinetics of our regulatory mutants.

### SI 1.4. Analysis of Growth curve data through logistic model fitting

For this study, we chose to employ a logistic growth model based on the modified Gompertz formula initially described by Zwietering *et al.* (1990) for the modelling of bacterial growth curves.[32] Such a model is frequently chosen for work of this type for its robustness and ability to provide accurate estimations of key growth parameters.[22,33]

$$\mathrm{OD_{660}(t)} = \frac{\mathrm{Y_{max}} - \mathrm{Y_0}}{1 + \exp\left(\frac{4\mu}{\mathrm{Y_{max}} - \mathrm{Y_0}}\,(\lambda - \mathrm{t}) + 2\right)} + \mathrm{Y_0}$$

Where $t$ = the time elapsed in hours, $Y_{max}$ = the maximum $OD_{660}$, $Y_0$ is the starting $OD_{660}$, μ = the maximum growth rate in exponential growth phase, and λ = the lag time before true exponential growth begins (defined mathematically here as the value of the x-axis intercept of a tangent line drawn through the inflection point[32,34]). As $\mu$ is an absolute rate (rate of change in OD) and not a specific rate (rate of change in OD relative to OD), it needs to be converted into a form that allows calculation of doubling time:

$$t_{\text{doubling}} = \frac{\ln(2)}{k_{\max}} = \ln(2) \cdot \frac{Ymax - Y0}{2\mu}$$

Analyses of the raw data were performed in *R* (version 4.5.1) via RStudio.

**Supplementary Section SI 2: BadM, the repressor of the anaerobic benzoate degradation operon, shows a large magnitude (but small scope) of effect based on wild-type vs. Δ*badM* experiments.**

<u>SI 2.1. BadM is the regulator of an iron-requiring gene cluster for anaerobic benzoate degradation:</u>

Among α-proteobacteria of the order Rhizobiales, three Rrf2-family regulators are well-established: RirA, the iron-responsive regulator in the taxonomic family Rhizobiaceae described previously; NsrR, the nitrite-responsive regulator whose only established presence in Rhizobiales is in genus *Xanthobacter*; and BadM, the acetamidobenzoate-responsive repressor of the *BadDEFG(AB)* operon.[5,35] Of these three Rrf2-family regulators, *Rhodopseudomonas palustris* has only the third. BadM (Rpal_0730) is well-studied as the co-regulator of the *bad* operon in CGA010, alongside AadR (Fig. S10).[36–39] BadM has a well-established role in literature as serving this narrow primary function as an anaerobic aromatic metabolic regulator. The BadDEFG holoenzyme catalyses a critical step in the anaerobic degradation of aromatic organic molecules (Fig. S10E).[5] The holoenzyme contains three predicted [4Fe-4S] clusters (Fig. S10F), and BadB (which works alongside it) is a ferredoxin,[5] making the Bad enzymes relatively iron-requiring for the cell to express. As BadM lacks the conserved cysteine residues necessary to form its own operative sensing [4Fe-4S] cluster, however, it has not been explored in the context of any potential overlap of function with RirA or IscR.[39,40]

Despite the thorough study of regulation of these genes and the action of their protein products, the acquisition of the bad operon in *Rhodopseudomonas* has not yet (to our knowledge) been examined in depth. In some preliminary analyses of gene neighbourhoods surrounding high-similarity badM homologues, it seems likely that *Rhodopseudomonas palustris* received BadM from *Rhodomicrobium* in a relatively recent horizontal gene transfer event (Fig. S11).[41] Even other species of *Rhodopseudomonas*, such as *R. rhenobacensis* and *R. pseudoplaustris*, as well as all species of neighboring genus *Bradyrhizobium*, appear to completely lack the locus. *Rhodomicrobium* (of the same order but different families) does possess a *badM* gene with ~45% identity to that of *R. palustris*, and the *badDEFG* operon adjacent to it with 65-70% identity throughout the four enzyme-subunit-encoding genes. All other variants of the locus appeared to be of only greater homological divergence.

While a small region of the BadD promoter was identified as DNAse-protected by BadM, the specific binding motif within that region has not yet been identified.[38] *Rhodopseudomonas* is not too distantly related to organisms relying heavily upon RirA and NsrR, being within the same family as Xanthobacter and same order as Rhizobium, so it is somewhat plausible that it may host Rrf2-binding DNA motifs in promoter regions that the gain or loss of these other factors in its taxonomic relatives may have left a vacuum open for off-target binding by BadM. Thus, because BadM is a relatively recent (re-) addition to the lineage of an Rrf2-family transcriptional regulator, we thought it worthy of testing whether there might be some vestigial Rrf2-sites lying dormant in the *Rhodopseudomonas palustris* genome in the promoters of ancestral targets like IscR or ones recently brought under regulation of RirA in its Rhizobiaceae cousins.

In some preliminary pilot work on this subject, Gupta (2020) showed that a Δ*badM* mutant strain showed a marked defect in iron oxidation and growth rates under photoferrotrophic conditions (unp. dissertation - 2020), raising the possibility that BadM might be serving some broader iron-related regulatory role (71). The fact that it works via concerted action with AadR (which does contain a [4Fe-4S] cluster) on the single canonical promoter regulated by it only made this more relevant to the current work, and as such, in parallel with the Δ*aadR* work we constructed and tested a Δ*badM* single deletion strain as well as a double mutant in the Δ*aadR* background (Δ*aadR*Δ*badM*) (Table ST3). The question we hoped to answer was whether the growth and iron oxidation defects observed by Gupta were due to potential off-target regulatory differences beyond the *bad* operon, or whether they were instead due to the metabolic burden of constitutively expressing staggering quantities of mRNA and protein that was ultimately metabolically frivolous, under a growth condition already at the fraying edge of what not even all *Rhodopseudomonas* strains can accomplish.

SI 2.2. Deletion of *badM* constitutively derepresses *badDEFG*, even during growth on benzoate:

Consistent with its established role as a transcriptional repressor, deletion of *badM* led to significant upregulation of *badDEFG* transcription relative to wild-type TIE-1 (**Fig**. **S12**). On FWBenzoate, where the effector molecule acetamidobenzoate (produced from benzoate by BadL) is expected to partially release BadM's repression in wild-type,[5] we nonetheless observed that *badDEFG* transcripts were significantly higher in Δ*badM* than in wild-type, with an average fold change of ~3.7 across the operon (individual genes ranging from ~2.4-fold for *badD* to ~5.3-fold for *badG*; **Fig**. **S10, S12**). This indicates that even during active growth on benzoate, BadM-mediated repression is not fully released in wild-type — the repressor still provides a "brake" on *badDEFG* expression that conservatively limits the total enzyme pool even under conditions ideal for maximum expression of the genes. Under conditions where benzoate is absent (FWAc, FWAc+Fe, YPS), the fold change between Δ*badM* and wild-type was dramatically larger, exceeding 400-fold on FWAc and approaching 600-fold on FWAc+Fe, reflecting the near-complete repression of *badDEFG* by BadM in the absence of its inducing ligand.

This substantial derepression did not significantly affect the growth rate of Δ*badM* on FWBenzoate relative to wild-type, but the mutant did grow to a significantly higher maximum biomass (as measured by $OD_{660}$; $p < 0.001$, **Fig. S10D**). This is consistent with the idea that more complete commitment to expression of the benzoyl-CoA reductase complex allows the Δ*badM* strain to more thoroughly metabolise the available benzoate, converting more substrate to biomass (possibly at the expense of other processes, such as secondary metabolism). This observation may have implications for biotechnological applications requiring efficient anaerobic degradation of aromatic compounds, such as the efficient conversion of lignocellulosic biomass to value-added products.[5,22,42]

SI 2.3. The displacement of the Δ*badM* transcriptomes in the PCA supports the interpretation of PC2 as an "iron requirement" axis, not just an iron-related expression axis:

Beyond confirming BadM's canonical role in CGA009 holds true in TIE-1, the Δ*badM* strain provides a critical test of our interpretation of the PC2 axis in Results (Main Text, **Fig. 4A**). The BadDEFG holoenzyme contains three predicted [4Fe-4S] clusters (inferred via structural homology with Geobacter metallireducens BamBCDE; **Fig. S12F**), and its co-substrate BadB is a [4Fe-4S]-containing ferredoxin,[38] making this enzyme complex one of the most iron-requiring in the TIE-1 genome. Constitutive expression of *badDEFG* in Δ*badM* therefore imposes a substantial iron demand on the cell, but only the *badDEFG* operon itself (and not the dozens of other genes in the AadR regulon) is affected. If PC2 not only reflects an "iron-related" axis but indeed reflects the balance of iron investment relative to iron supply (i.e., an "iron-spendy vs. iron-thrifty" transcriptome axis) then constitutive BadDEFG expression should shift the Δ*badM* transcriptome along PC2 primarily when iron is limiting, but not when it is abundant.

We observed that this was indeed the case (**Fig. S13**). In the PCA displacement analysis, Δ*badM* showed its largest transcriptomic displacement on FWAc (28 units), where iron is at basal levels and the constitutive iron demand of wantonly producing BadDEFG represents a burden on the cell's iron budget (**Fig. S13C; Fig. S6D**). On FWAc+Fe and FWB, the displacement was substantially smaller, and on YPS (aerobic) it was negligible. This shows that having enough iron to spare, having the produced BadDEFG be metabolically useful instead of a burden, and/or removal of the constitutive anaerobic activation of the locus by AadR, respectively, may rescue the defective mutant phenotype, resulting in performance near-identical to that of wild-type. This pattern is the opposite of what was observed for Δ*aadR*, whose largest displacement was on FWAc+Fe (42 units) (Main Text, **Fig. 4C**). As losing AadR disrupts the cell's management of iron-responsive gene expression, the consequences are greatest when iron is plentiful and there is more to manage. In contrast, Δ*badM*'s displacement is greatest when iron is scarce and the constant mass-production of unnecessary iron-requiring enzymes becomes a measurable burden.

Together, the Δ*aadR* and Δ*badM* displacement patterns support the interpretation that PC2 captures the cell's investment in iron-requiring enzymes relative to iron availability. AadR coordinates iron resource investment broadly, while BadM controls a single locus of narrow scope (*badDEFG*) within it. Removing the global regulator AadR causes widespread discoordination that manifests most strongly under iron-supplemented conditions, while removing a single brake (BadM) on one iron-requiring enzyme causes a localised cost that manifests most strongly when the limiting resource is most scarce.

SI 2.4. Conditional effects of *badM* deletion beyond the *bad* operon:

Although BadM's primary regulatory target is the *badDEFG* operon, we noted a small set of genes conditionally affected by *badM* deletion, specifically under basal-iron conditions. Among these, the denitrification-related genes *nirK2* and *norCBQDE* were derepressed in Δ*badM* under FWAc (basal iron) but showed no significant change under FWAc+Fe. As NorCB (nitric oxide reductase) and the associated *cbb*$_3$ oxidase subunits are haem-iron enzymes, their conditional derepression in an iron-limited background is consistent with a secondary consequence of altered iron partitioning in Δ*badM*: when iron is scarce and the constitutively expressed BadDEFG complex is consuming [4Fe-4S] clusters, reduced iron availability for haem biosynthesis may indirectly relieve repression of these genes. When iron is abundant (FWAc+Fe), the additional iron demand from BadDEFG is easily met, and no downstream effects on haem-iron enzymes are observed.

While that is by far the most parsimonious explanation, a small possibility remains open that BadM could be directly regulating the locus, considering that NsrR is an ancestral nitrite-responsive Rrf2-family transcription factor, which directly regulates *nirK* expression in the *β*-proteobacterium *Nitrosomonas*.[35,43–45] The gene was lost in the lineage of order Rhizobiales in all but *Xanthobacter,*[35] which does mean that a relatively recent ancestor of *Rhodopseudomonas* likely did have a copy of it, lost long before the *badM* gene was acquired. Whether BadM directly regulates the *nirK2/nor* region or whether this effect is entirely indirect remains to be determined, but the observation reinforces the broader theme that iron allocation has pleiotropic transcriptomic consequences even when the primary regulatory perturbation is narrow in scope.

**Supplementary Section SI 3: Meta-analysis of the AadR-FixK negative feedback loop and the four Fur-family regulators present in *R. palustris* TIE-1.**

SI 3.1. Oxygen and Light sense-response transcriptional regulation is controlled by the AadR-FixK negative feedback loop in *Rhodopseudomonas*:

In *Bradyrhizobium japonicum* (*Bj*), a close phylogenetic relative of *Rhodopseudomonas palustris* (*Rp*), the aerobic-anaerobic shift is controlled by the expression of CRP-Fnr family regulator genes *Bj fixK*$_1$ and *Bj fixK*$_2$, which are activated by the FixL-J two-component regulatory system. Sensing a drop in extracellular oxygen, FixL-J activates expression of *Bj fixK*$_2$*,* the product of which in turn activates *Bj fixK*$_1$ expression.[1,46] In *Rhodopseudomonas*, the transcriptional regulator *Rp* FixK is orthologous to *Bj* FixK$_2$, with 72% amino acid identity, a comparable gene neighbourhood, and a role regulating transcription during decreasing oxygen levels from aerobic to microaerobic conditions after activation by FixLJ.[1,46–48] The precise number of operons and genes regulated by *R. palustris* FixK has been challenging to determine due to the similarity of its binding site to that of other Fnr-family regulators, especially the close paralogue AadR (the *Rp* orthologue of *Bj* FixK$_1$),[1,35,49] but it is well-established that FixK is a global metabolic regulator with at least 55 targets with altered expression in a CGA010 *fixK* mutant having a palindromic FixK box (TTGA*T*-N$_4$-*A*TCAA) in their regulatory region.[1,50]

AadR shares a relatively high 59% positional amino acid sequence identity with *Bj* FixK$_1$ (and is 92% identical within the active helix-turn-helix motif),[36] and it was first characterised as regulating the benzoate and 1-hydroxybenzoate in *Rhodopseudomonas palustris* CGA009 (and was initially named for its characterised scope as an anaerobic aromatic degradation Regulator).[36,37] Transcription of *aadR* is activated by FixK, and in turn AadR represses *fixK* expression, as well as activating the metabolic gene clusters *badDEFG* for anaerobic benzoate degradation and *hbaBCD* (indirectly, via *hbaR* activation) for anaerobic 4-hydroxybenzoate degradation.[1,36]

AadR's role was later characterised well beyond aromatic degradation in an analysis by Rey and Harwood (2010) to include a myriad of loci: it repressed more than 20 genes involved in ferric iron transport and metabolism, activated a putative iron response regulator (*rpa2339*/*Rpal_2583,* Irr homologue 1 of the two encoded), and activated six genes of unknown function (*rpa2333-2338*/*Rpal_2577-2582*)[1] — since identified as the *fez* operon for ferrosome assembly, a class of iron storage organelles.[51] AadR also repressed ~80 genes including *fixK* and several FixK-activated targets such as the *cbb*$_3$-oxidase,[1] suggesting opposing regulatory activity between AadR and FixK on shared targets. However, these observations were from a single microaerobic growth condition on 10 mM succinate + 3 mM p-coumarate (to induce activation of aromatic degradation genes)— iron was not varied as an experimental parameter, and no fully anoxic condition was tested. The list of genes in *Rhodopseudomonas palustris* under FixK regulation includes *pioABC* for utilisation of Fe(II),[3] and at least five other major transcriptional regulators, among them *aadR*— though, surprisingly, *nifA* does not appear to be under its control as it is in *Rhizobiaceae.*[1] Despite *pioABC* having two canonical FixK boxes in its promoter region, only one of these (FixK-BSI) was shown to bind purified FixK in vitro by Bose and Newman (2011).[3] The similarity (or possibly identicality) of the FixK box motif to the site bound by its own paralogue AadR has made disambiguating the regulons of the two regulators prohibitively challenging.[1,49] Due to this similarity, they were unable to fully deconvolute whether their co- and anti-regulation of shared sites happens via direct or indirect means (i.e., repression of *fixK* leading to indirect repression of FixK's targets).[1]

AadR protein activity is almost certainly suppressed by the presence of oxygen, which it is hypothesised to sense via its conserved cysteine residues (**Fig. S16A**) which form a [4Fe-4S] centre in other Fnr-family homologues.[37] *E. coli* FNR is the archetype of the family, which senses oxygen via an iron-sulfur cluster. The spacing between the key N-terminal cysteine residues (3 amino acids, then 2 AAs, then 5 AAs) and the presence of one distant in primary structure (at position 140 (**Fig. S16B**)) but positioned closely in secondary structure allows it to hold the cluster. Other FNR-family regulators, such as DNR (which senses NO), differ in their cysteine-rich domain architecture. *R. palustris* and *Bradyrhizobium* FixK proteins are divergent in sequence from the others through this region, completely lacking 3 to all 4 of the key residues and therefore lack a cluster and are unable to sense oxygen. Other regulators, such as Rrf2-family RirA (not aligned here due to the far greater sequence dissimilarity) have similar cysteine-rich domains that coordinate a [4Fe-4S] that senses not just oxygen but iron.[40,52]

AadR is more similar to FixK than to other FNR-family regulators, owing to the recent paralogous duplication of *fixK* from *aadR.*[46,50] While FixK has lost the cysteines, AadR retains them, though in an odd spacing compared with other FNRs— spacing between cysteines of 4 AAs (instead of 3), 3 AAs (instead of 2), and 7 AAs (instead of 5) (**Fig. S16A**). RirA also has greater spacing between its cysteines (e.g., …C-[5 AAs]-C-[7 AAs]-C… ).[53] Thus, it is perhaps somewhat plausible that AadR's putative cluster may also serve an iron-sensing role. Because of this putative [4Fe-4S], it would be a formidable challenge to purify the AadR protein for binding site analyses or other in vitro work. What is clearly known is that with AadR and FixK counterbalancing one another, the expression of only the sets of genes relevant to current environmental conditions is fluently maintained over the shift from aerobic to anaerobic growth.

### SI 3.2. Despite lacking Fur, TIE-1 encodes regulators in the family with shared and divergent qualities

The ancestor of orders Rhizobiales, Rhodobacterales, and Rhodospirillales gained the gene for the Fur-family protein Irr (iron-responsive regulator), and the ancestral Fur functionally diverged to become Mur, primarily regulating manganese uptake instead of iron.[54,55] In order Rhizobiales (renamed recently to Hyphomicrobiales), IscR was then lost, leading to somewhat of a gap in effective iron sense-response. As iron is so critical for the metabolisms Rhizobiales are known for (nitrogen fixation, anoxygenic photosynthesis, etc.), evolution filled the gap differently in each familial lineage. In *Rhizobiaceae*, the Rrf2-family protein RirA (Rhizobiales iron-responsive regulator A) is the conserved mechanism for the sense-response of iron and oxygen,[56] which it performs via an integral iron-sulfur cluster.[40,52] Descendants of family *Nitrobacteraceae*, which includes *Rhodopseudomonas* and *Bradyrhizobium*, are instead considered to regulate iron response via the Fur family.[56–59]

Several Rrf2-family regulators contain these Fe-S clusters, which may sense oxygen, iron levels, and/or cellular redox status,[43] including the *Rhizobium/Sinorhizobium* RirA.[59,60] RirA (Rhizobiales iron-responsive regulator) forms a homodimer and binds IRO Box sequences to repress expression of iron-requiring genes during iron-limited conditions, while under iron-replete conditions this repression is lifted.[40,52] Pellicer Martinez *et al.* (2019) revealed the mechanism of this sensing: the reversible and oxygen-independent dissociation of $Fe^{2+}$ from the $[4Fe\text{-}4S]^{2+}$ cluster converts it to a $[3Fe\text{-}4S]^{0}$ cluster, stopping the RirA protein from binding IRO Boxes when intracellular iron levels are sufficient to drive this cluster degradation reaction.[52] In *Bj* and *R. palustris*, however, no characterised RirA homologue is present.[60] IrrA (Iron responsive regulator) is thought to directly activate many of the missing RirA's targets (iron transport, TCA cycle, and haem utilisation (via *hmuP*)) while continuing to repress its own conserved targets haem synthesis, iron export, and storage.[57,59,60]

*R. palustris* is known to encode two copies of the Fur-family iron regulator *irr*/*irrA*, numbered here according to the schema proposed by Rodionov *et al.* (2006): *irrA2* (*Rpal_2583*), and *irrA1* (*Rpal_0428*).[54] While multiple other closely related species (*Bradyrhizobium japonicum, Rhizobium leguminosarum, etc.*) also possess paralogous duplications of *irrA* (which Todd *et al.* (2006) term *irrA* and *irrB*), the pair of these genes in *Rhodopseudomonas* appear to both cluster more closely with the *B. japonicum irrA* than either does to *B. japonicum irrB*, hence why we term them *irrA2*and *irrA1* here. IrrA1 retains the residues necessary to coordinate a haem-recognition motif unique to *Xanthobacteraceae*,[54,57] allowing it to regulate haem-related metabolism in response to cellular haem iron levels, while *irrA2* is missing that motif. *irrA2* is adjacent to the *fez* operon (**Fig. S1**), which encodes the machinery for formation of ferrosomes — a relatively recently discovered class of iron-storage organelles expressed under iron-limited conditions.[51] *irrA1* is in a separate genomic location, and it is not physically adjacent to anything predicted to be particularly iron-relevant. As *irrA1* is the one with the ancestral haem-sensing motif preserved in its sequence, *irrA2* may serve a different role, possibly even regulating the *fez* operon it shares a promoter region with.

**SI Figures:**

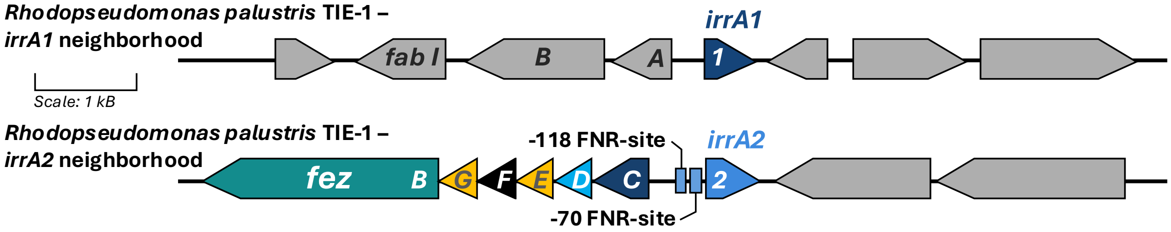


**Figure S1. Gene neighbourhoods for the two *irrA* paralogs found in the genome of *R. palustris* TIE-1.** The gene encoding IrrA2 is upstream (though in the opposite sense) of the *fez* ferrosome operon.[51] It has two predicted FNR-family regulator binding motifs in its promoter region.[35] The gene encoding IrrA1 (which retains the haem-sensing residues) is at an entirely separate locus, ~2000 genes away.

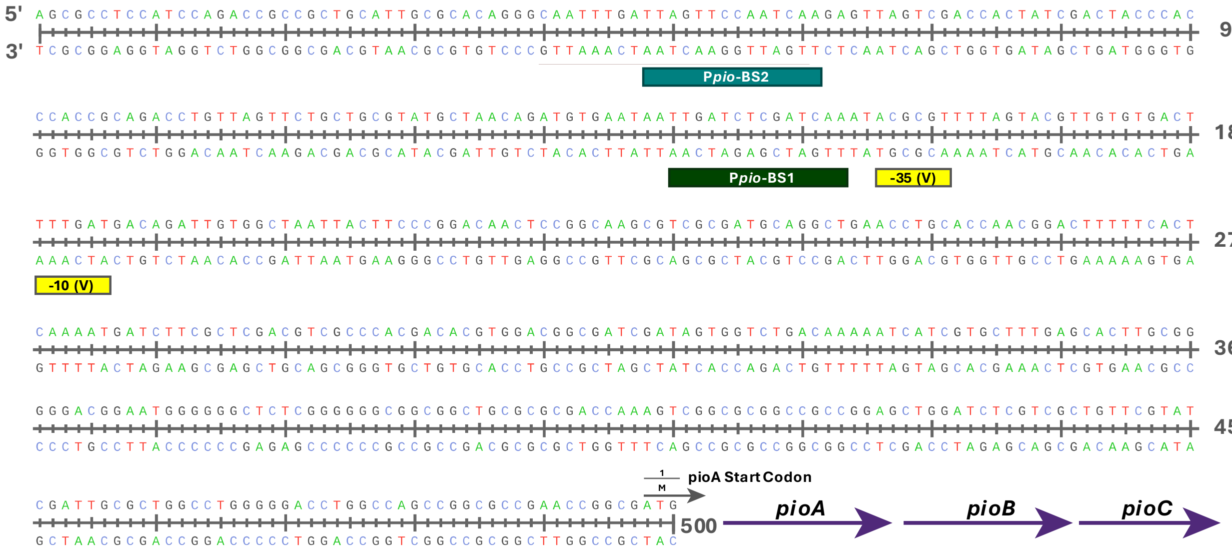


**Figure S2.** ***pioABC* promoter (P*pio*) region map.**

The 500 bp upstream of the TIE-1 *pioABC* operon (including the start codon of *pioA*) are represented above. The two regulator binding sites validated by Bose and Newman (2011) are indicated: P*pio*-BS1, which is bound by FixK *in vitro*, and P*pio*-BS2, which is not.[3] Both have a sequence motif predictive of a CRP/FNR-family regulator as the cognate factor, though the motif is more perfect in BS1 than BS2. The -35 and -10 sites (both validated, (V)) are also shown.

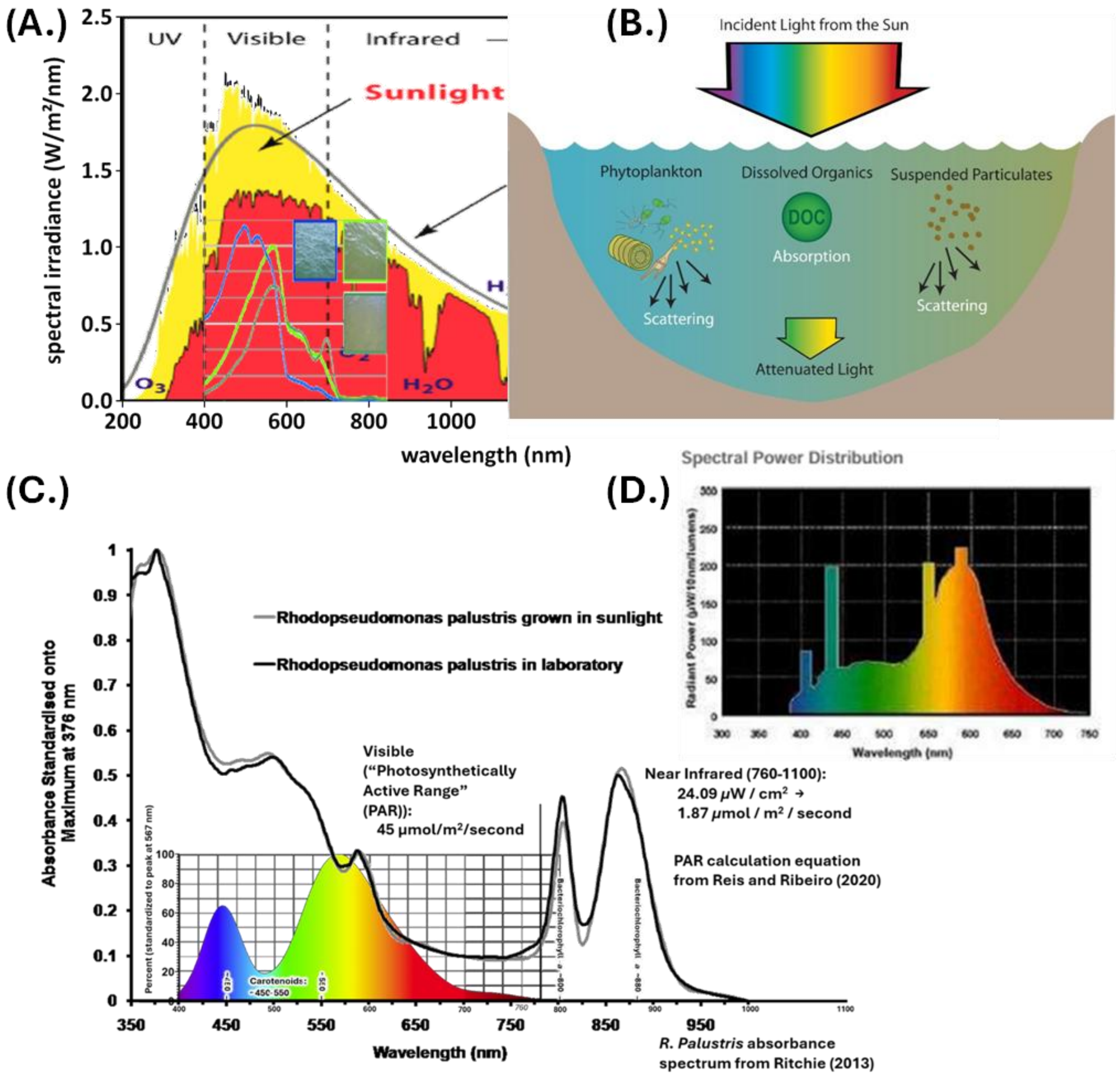


**Figure S3. Naturomimetic growth conditions were designed to match the attenuated light present in wetlands.**

Above: Visual of attenuation of sunlight into wetlands. **(A)** Sunlight spectrum (yellow is sunlight in space, red is sunlight at sea level) adapted from.[61] Spectra of light in waters of different turbidity (blue = low turbidity, light green = medium turbidity, light green =high turbidity) adapted from[62] (x-axis scale adjusted, y-axis not scale adjusted). **(B)** Attenuation of light in the water column of turbid wetlands leads to a yellow-shift in spectrum. Adapted from.[63] **(C)** Wavelengths emitted by the GE fluorescent light tubes in our 30°C phototrophy growth chamber setup. *R. palustris* bacteriochlorophyll a peaks per[64] and absorbance spectrum per[8] show only at the minor peak (~580 nm) is there any good overlap between the spectrum produced by our lighting and the spectrum usable by *R. palustris*. PAR calculated for each range (visible and IR) based on two distinct light meters, using the equations in.[65,66] **(D)** Spectral power distribution graph (inset, top right) is from the General Electric (GE) Lighting Specification Sheet for the #23075 – F60T12/CWHO bulbs ("T12 fluorescent tube lights") used in here.

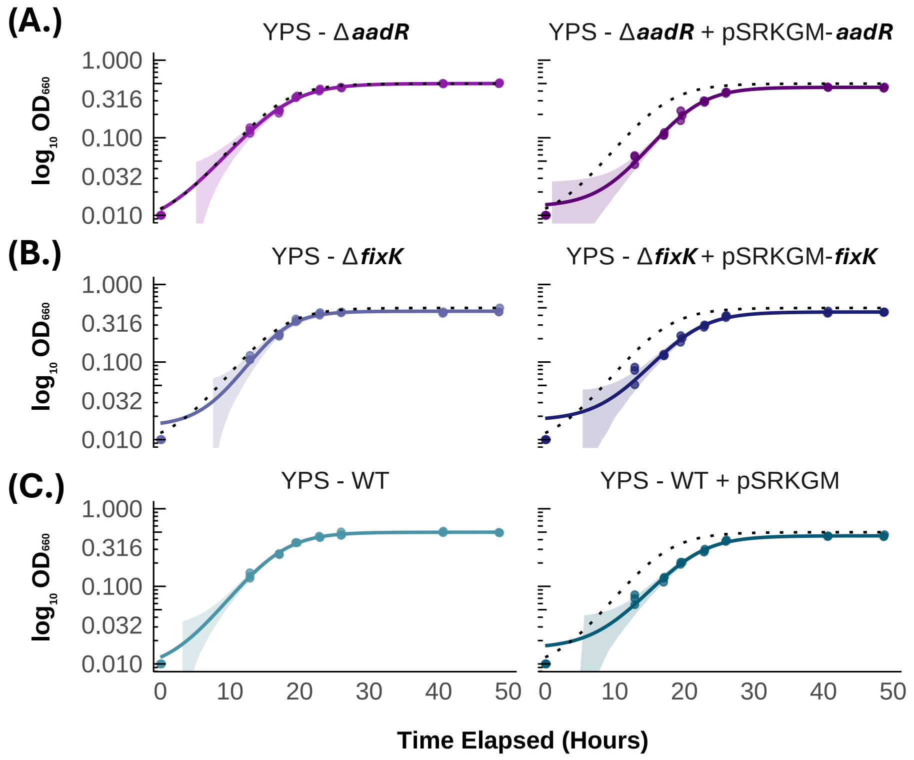


**Figure S4. Growth of mutant strains and their complements on YPS (aerobic, dark).** Complements were grown in identical media, but amended with 400 µg/mL gentamicin and 10 mg/mL IPTG. **(A.)** Growth of Δ*aadR* and the *aadR* complement strain (Δ*aadR*::pSRKGM:*Plac-aadR*). **(B.)** Growth of Δ*fixK* and the *fixK* complement strain (Δ*fixK*::pSRKGM:*Plac-fixK*). **(C.)** Growth of wild-type and wild-type with empty vector. Black dotted lines represent growth of wild-type for comparison. No strain showed a significant difference in lag time or doubling time. The ΔfixK strains showed a defect in maximum $OD_{660}$ achieved, but this phenotype has been previously reported and is expected due to the defect in photosystem expression of that strain.[3] For a table of calculated curve-fit values, please see **Supplementary Dataset S08a**.

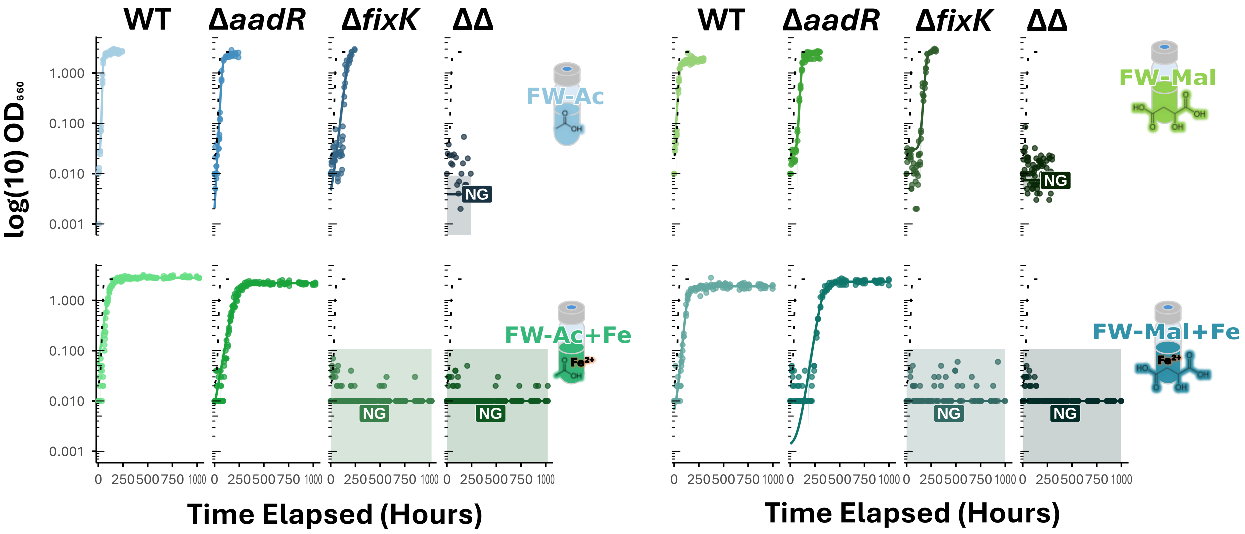


**Figure S5. Δ*fixK* did not grow on FWAc+Fe or FWMal+Fe, even though it grew on each carbon source under –Fe conditions.**

NG = "No Growth". The double mutant (Δ*aadR*Δ*fixK*) did not grow on either FWAc or FWMal, regardless of iron concentration. For a table of calculated curve-fit values, please see **Supplementary Dataset S08b**.

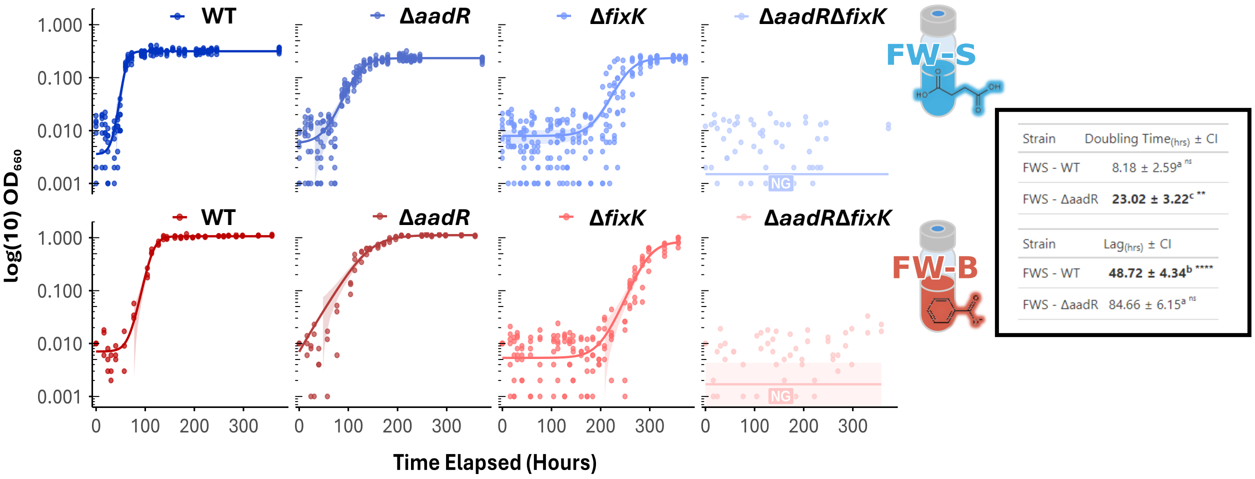


| Strain | Doubling Time(hrs) ± CI |
|---|---|
| FWS - WT | 8.18 ± 2.59[a] ns |
| FWS - ΔaadR | **23.02 ± 3.22[c] **** |

| Strain | Lag(hrs) ± CI |
|---|---|
| FWS - WT | **48.72 ± 4.34[b] ****** |
| FWS - ΔaadR | 84.66 ± 6.15[a] ns |

**Figure S6. Growth of wild-type and mutant strains on Freshwater + Benzoate.**

TIE-1 Δ*aadR* grew significantly more slowly than wild-type on FWB (Benzoate), as expected,[36] but also showed a significant growth defect on FWS (Succinate), unexpectedly. NG = "No Growth". The double mutant (Δ*aadR*Δ*fixK*) did not grow on either FWS or FWB, regardless of iron concentration. Inset table shows doubling times and lag time in hours of WT and Δ*aadR*. *(inset)* Significance was calculated using two-way ANOVA with Tukey's correction; stars indicate *p*-values as stated in **Table ST7**. For a table of calculated curve-fit values, please see **Supplementary Dataset S08c**.

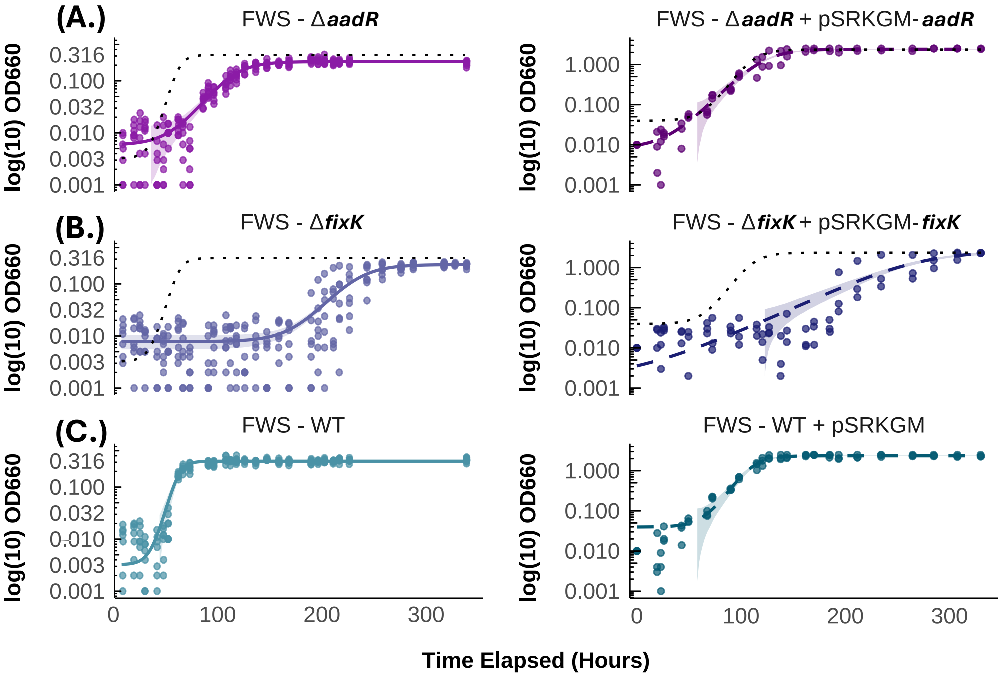


**Figure S7. Growth of strains and their complements on FWSucc. photoheterotrophy.**

**(A.)** Growth of Δ*aadR* and the *aadR* complement strain. **(B.)** Growth of Δ*fixK* and the *fixK* complement strain. **(C.)** Growth of wild-type and wild-type with empty vector. Black dotted lines represent growth of wild-type (and wild-type plus empty vector) for comparison. See **Supplementary Dataset S08d**.

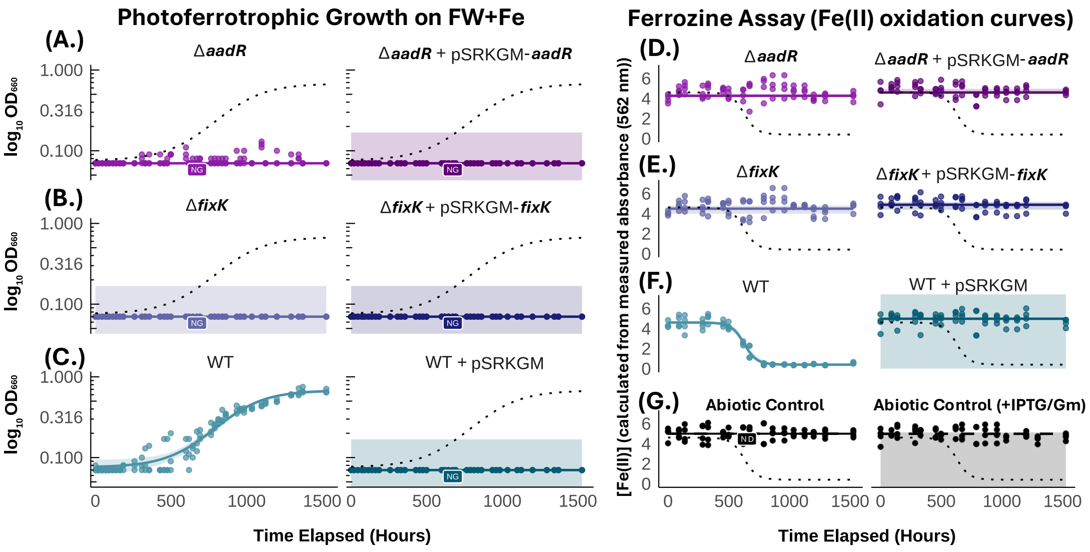


**Figure S8. Growth of strains and their complements on FW+Fe photoferrotrophy.**

**(A.)** Growth of Δ*aadR* and the *aadR* complement strain. **(B.)** Growth of Δ*fixK* and the *fixK* complement strain. **(C.)** Growth of wild-type and wild-type with empty vector. Black dotted lines represent growth of wild-type for comparison. **(D.)** Fe(II) oxidation by Δ*aadR* and the *aadR* complement strain. **(E.)** Fe(II) oxidation by Δ*fixK* and the *fixK* complement strain. **(F.)** Fe(II) oxidation by wild-type and wild-type with empty vector. **(G.)** Fe(II) oxidation in the abiotic control and the complement strains abiotic control (same medium but amended with IPTG and gentamicin). See **Supplementary Dataset S08e-g**.

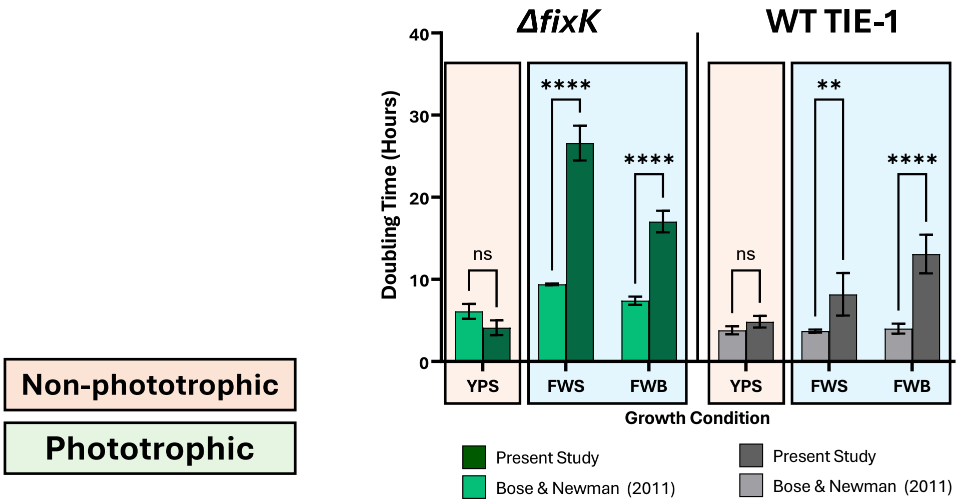


| Šídák's multiple comparisons test | Mean Diff. | 95.00% CI of diff. | Below threshold? | Summary | Adjusted P Value |
|---|---|---|---|---|---|
| YPS:B&N *ΔfixK* vs. YPS:Us *ΔfixK* | 1.990 | -1.145 to 5.125 | No | ns | 0.3990 |
| FWS:B&N *ΔfixK* vs. FWS:Us *ΔfixK* | -17.20 | -20.33 to -14.07 | Yes | **** | <0.0001 |
| FWB:B&N *ΔfixK* vs. FWB:Us *ΔfixK* | -9.640 | -12.77 to -6.505 | Yes | **** | <0.0001 |
| YPS:B&N WT TIE-1 vs. YPS:Us WT TIE-1 | -1.040 | -4.175 to 2.095 | No | ns | 0.9254 |
| FWS:B&N WT TIE-1 vs. FWS:Us WT TIE-1 | -4.480 | -7.615 to -1.345 | Yes | ** | 0.0025 |
| FWB:B&N WT TIE-1 vs. FWB:Us WT TIE-1 | -9.080 | -12.21 to -5.945 | Yes | **** | <0.0001 |

**Figure S9. Comparison of the effect on growth rate of wild-type and Δ*fixK* TIE-1 under conventional high incandescent light and naturomimetic fluorescent light.**

Data for high light (lighter shaded bars) are from the data released by Bose and Newman (2011).[3] Data for naturomimetic light in the present study (darker shaded bars) are from growth of the same strain genotypes in the same growth medium and glassware types. No significant difference was observed in growth under chemoheterotrophy in the dark (orange background), i.e., when light is not a changed variable between the two studies. A significant defect in doubling time was observed in not just Δ*fixK* but wild-type as well, under multiple different carbon sources (FWS = Freshwater medium + succinate; FWB = Freshwater medium + benzoate). Significance of difference was assessed using a one-way paired ANOVA of means, Šidák-corrected for multiple comparisons; stars indicate *p*-values per the table.

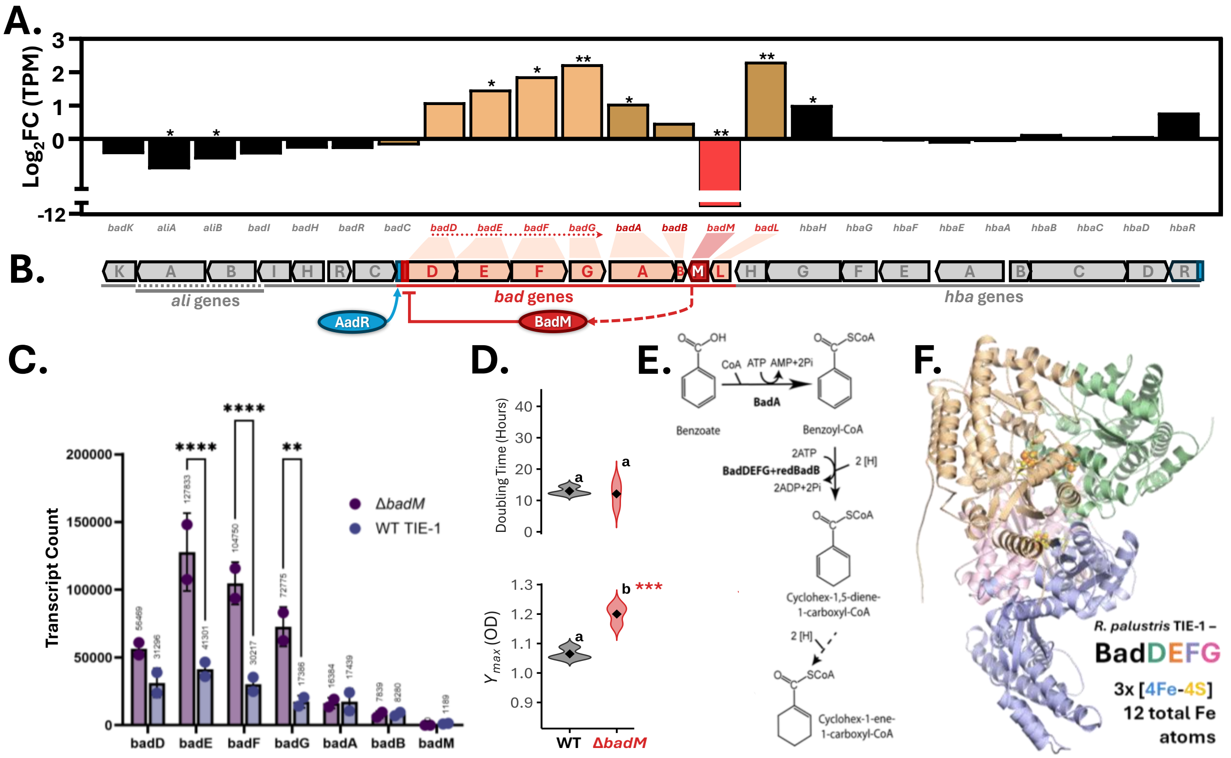


**Figure S10. Δ*badM* has greater transcriptional expression of BadDEFG, the critical enzyme for anaerobic reductive dearomatisation, than WT TIE-1 on FWB.**

**(A.)** $Log_2$ fold change (FC) in transcripts-per-million (TPM)-normalised expression in the Δ*badM* strain vs WT TIE-1. In Δ*badM*, *badDEFG* are expressed an average of 3.3-fold greater than in WT TIE-1 on FW media + 1 mM benzoate. Stars indicate *q*-values per **Table ST7** (*n*=2 biological replicates, DESeq2). **(B.)** Diagram of the *hba-bad-ali* gene cluster for anaerobic aromatic acid degradation. The *badDEFG* genes form an operon under control of two known transcriptional regulators. Repression of the *badD* promoter by BadM is released by its binding of acetamidobenzoates (produced from benzoate in a reaction catalysed by BadL), allowing AadR-activated transcription to occur unimpeded. **(C.)** When *badM* is deleted, *badDEFG* expression is significantly increased over WT, even during growth on benzoate (where it would be expected that repression by BadM would be fully released). Stars represent significance via pairwise 2-way ANOVA, Tukey-adjusted for multiple comparisons; stars indicate *p*-values per **Table ST7**. **(D.)** Δ*badM* grew at the same rate as WT TIE-1, but it grew to a significantly higher biomass (as measured via the proxy of optical density). **(E.)** Abbreviated figure showing the key *badDEFG*-*badA*-badB-catalysed steps in the anaerobic dearomatisation reaction. Figure adapted from VanDrisse & Escalante-Semerena (2018).[5] **(F.)** BadDEFG holoenzyme structure with the three [4Fe-4S] clusters shown (inferred via structural homology with *Geobacter metallireducens* BamBCDE (PDB 4Z3X)). Structural prediction made using AlphaFold3 (Google DeepMind, ptm=0.82, iptm=0.78, including low-confidence terminal IDRs), and visualised in Open Source PyMol.

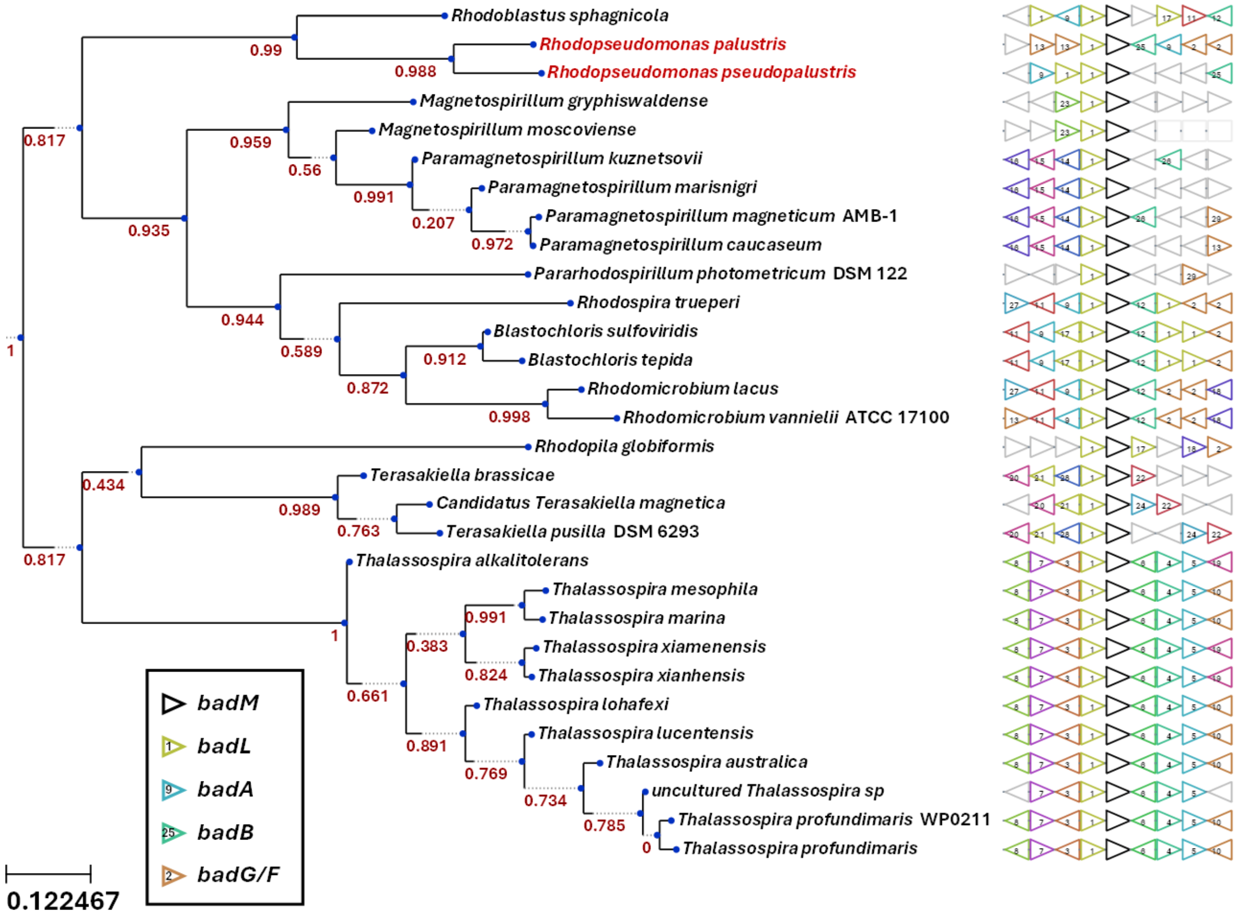


**Figure S11.** ***badM* and *badDEFG…* gene neighbourhood synteny plot shows that *Rhodopseudomonas palustris* either retains a frequently-lost ancestral cluster, or it got its benzoate degradation abilities from inter-family horizontal gene transfer.**

***Left:*** species tree showing taxonomic relationships between genera/species with best *badM* orthologs. *Rhodopseudomonas* stands alone in its family— *Bradyrhizobium*, *Nitrobacter*, and even some other *Rhodopseudomonas* species (e.g., *R. rhenobacensis*) do not possess this locus. The closest relative to the *R. palustris* cluster is in *Rhodoblastus*, in family *Roseiarcaceae* (of the same order). Aside from *Blastochloris*, the remaining clusters of note are all in either order *Rhodobacterales* or *Rhodospirillales*.
***Right:*** Gene synteny diagram showing clusters, centred on *badM* (black). *badL* (1), *badB* (25), *badA* (9), *badG/F* (2) are also shown.

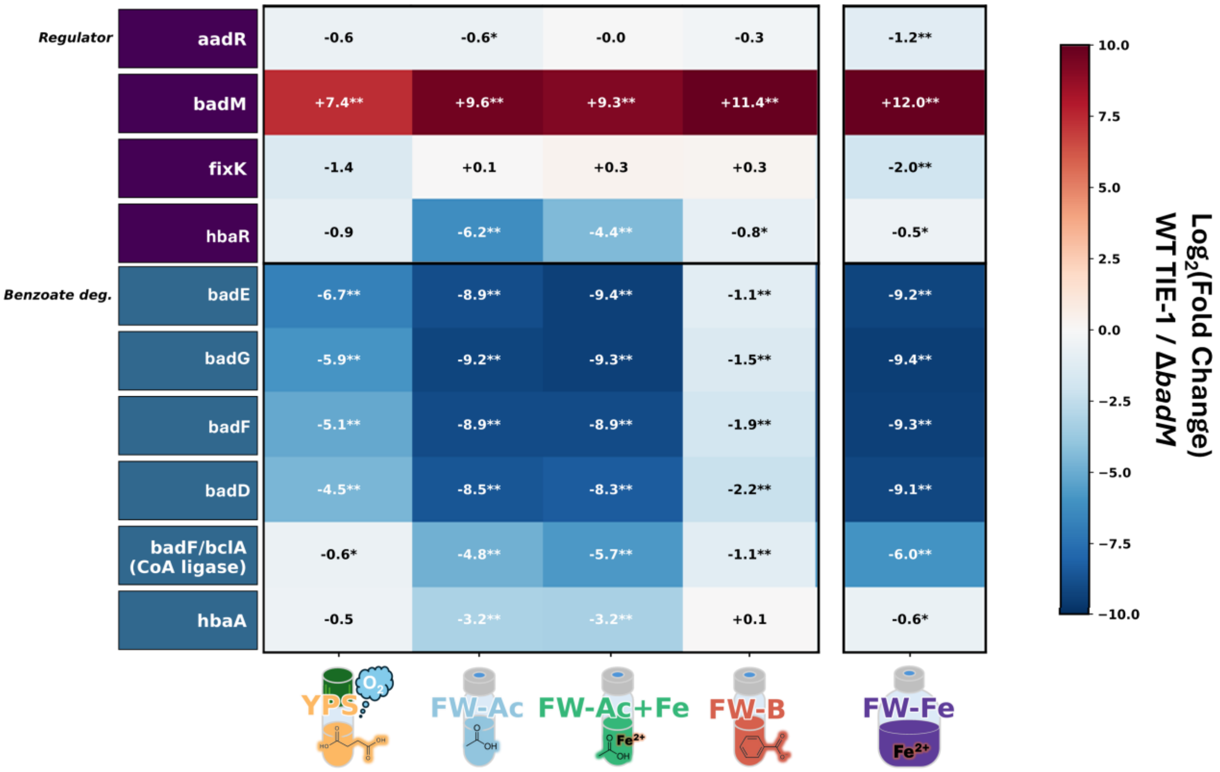


**Figure S12. Deletion of *badM* constitutively derepresses the *bad* genes.**

Heatmap showing $Log_2FC$ in transcript of relevant genes. Stars indicate *p*-values per **Table ST7**.

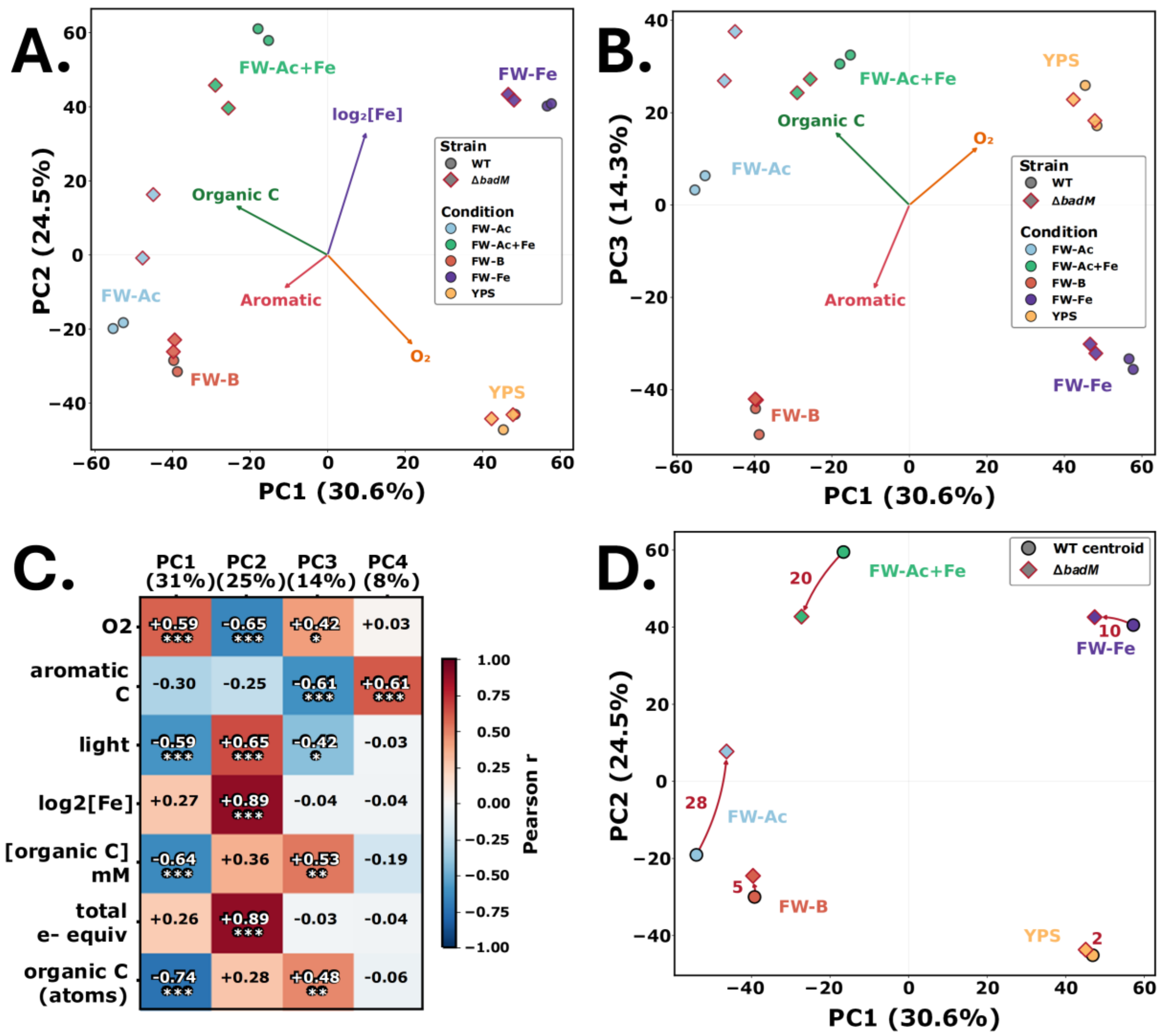


**Figure S13. Principal component analysis (PCA) reveals drivers behind the transcriptome displacement of Δ*badM*.**

**(A.)** PCA biplot of PC1 vs PC2. Metadata vectors and biological replicate points for VST-adjusted whole transcriptomes are shown. PC1 explains 30.6% of the variation, while PC2 explains 24.5%. **(B.)** As A, but in this panel comparing PC3 vs PC1. PC3 explains 14.3% of the variation between samples. **(C.)** Correlation matrix, mapping metadata indices to principal components. PC1 correlates positively with oxygen, and negatively with light and amount of organic carbon provided. PC2 correlates very strongly (r=+0.89, $p$<0.001) with iron concentration, positively with light, and negatively with oxygen. **(D.)** PC2 vs PC1 displacement plot of how the Δ*badM* knockout mutation shifted the transcriptome relative to that of wild-type on each growth condition (centroids of replicate point averages are shown). **Throughout:** Tested factor metadata include growth under aerobic/anaerobic conditions, presence/absence of an aromatic carbon source, light/dark, iron concentration, organic carbon provided, and total reducing (electron) equivalents provided (organic C and/or $Fe^{2+}$). PCA was performed to correlate whole transcriptome differential expression ($n$=2) in wild-type and Δ*badM* TIE-1 for growth conditions (see Methods). Metadata arrows show correlation of metadata indices with principal components.

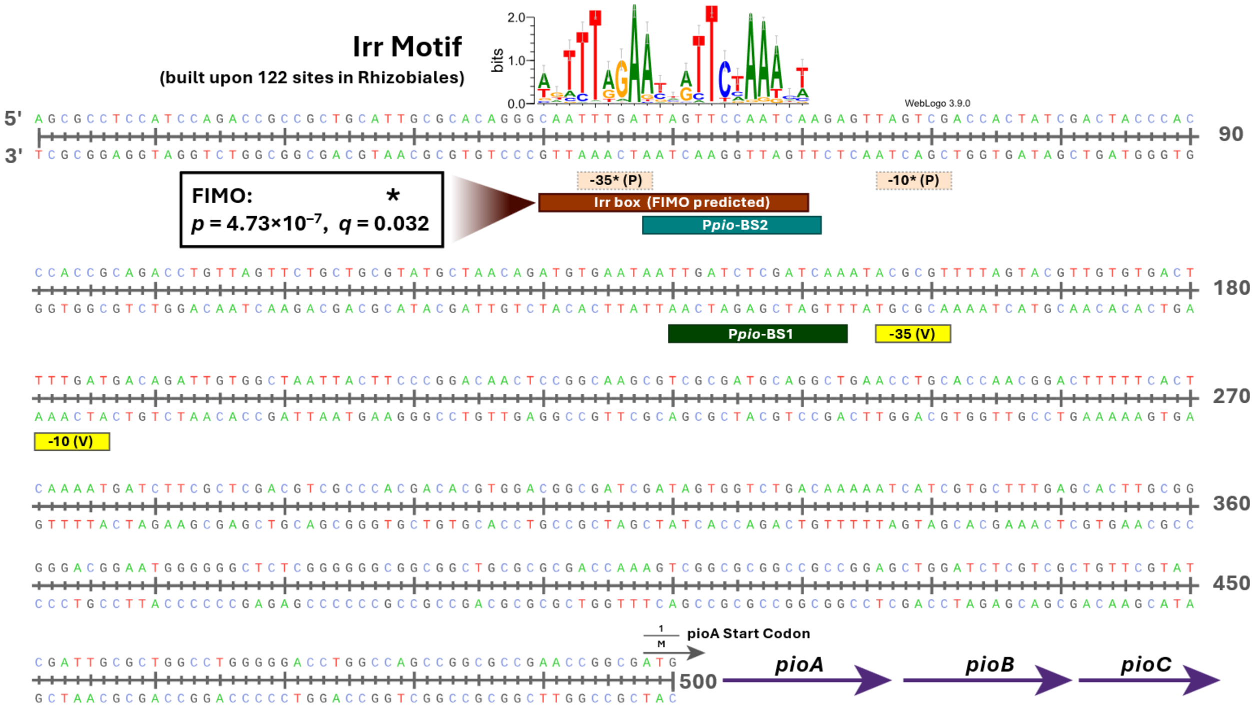


**Figure S14. Promoter region of *pioABC* (out to -497 nt), with the FIMO-predicted Irr binding site overlapping P*pio*-BS2 shown.**

Using FIMO (Find Motif Occurrence, MEME-suite,[67,68]) with the consensus IRR-binding motif built upon 122 Irr-regulated sites across order Rhizobiales (RegPrecise,[35]), we identified a significant prediction for an Irr site overlapping P*pio*-BS2 ($p = 4.73 \times 10^{-7}$ , $q = 0.032$). Also shown are the verified (V) -35 and -10 sites, as well as the second pair of sites predicted (P) using SAPPHIRE.

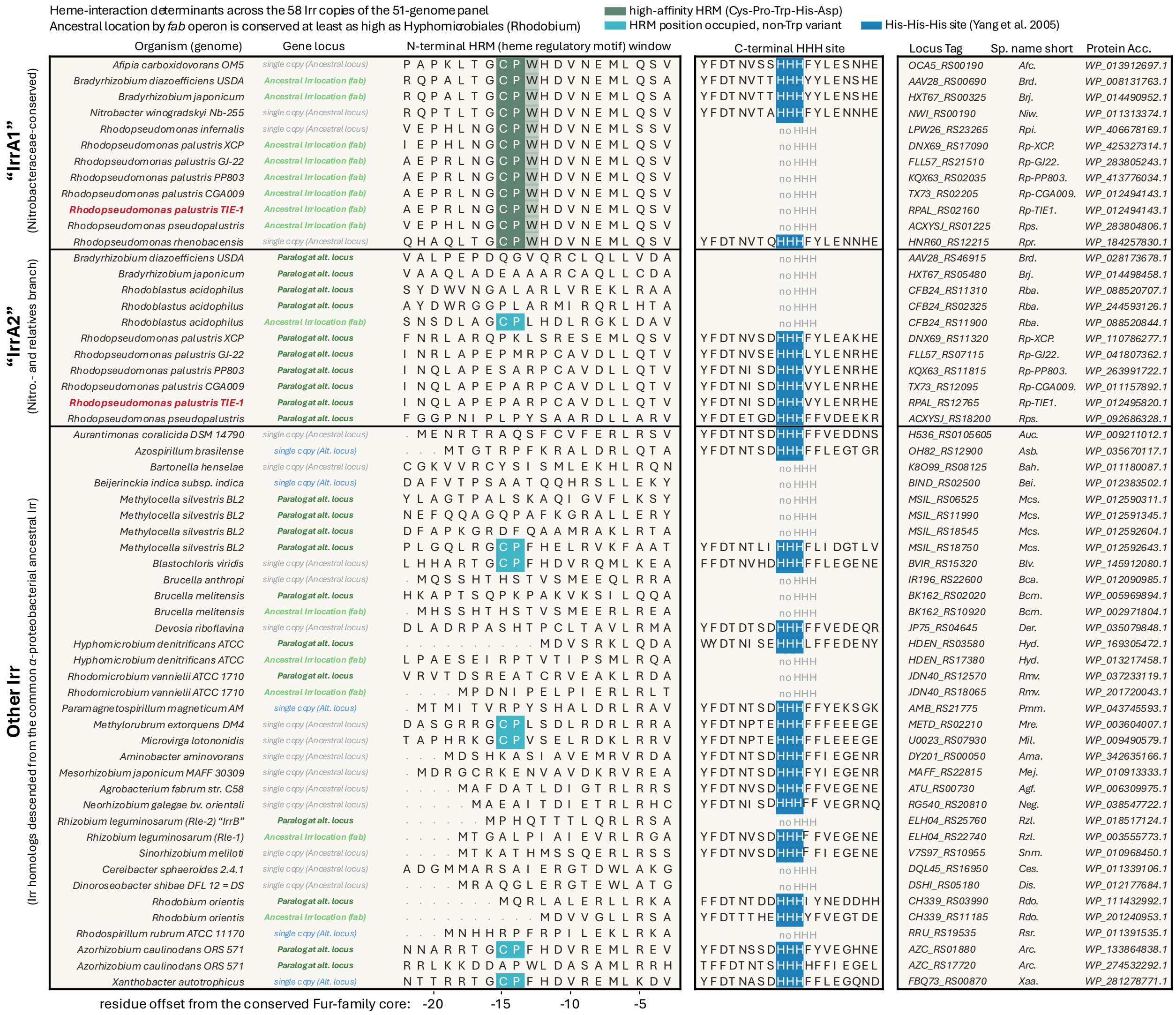


**Figure S15. The high-affinity haem-regulatory motif of IrrA marks the ancestral *fab*-locus copy across the *Nitrobacteraceae* and is absent from the derived paralogue**

Sequence view of the two published Irr haem-interaction determinants across all 58 Irr-family copies in the 50-genome panel: the N-terminal haem-regulatory motif (HRM; Cys-Pro, high-affinity when immediately followed by Trp) and the C-terminal histidine triad (HHH; determinants per Qi & O'Brian (2002), Yang *et al.* (2005), and Todd *et al.* (2006)).[69–71] Rows are grouped as ancestral *fab*-locus copies, derived paralogues at alternative loci, and single-copy-bearing genomes (at either the ancestral or alternate locus); each row gives species, locus assignment, protein accession and the short abbreviation as labelled in the **Fig. 5** tree. Positions were read from raw sequence anchored on the conserved Fur-core landmark (trimming off the variable N-terminus); on this anchor every Cys-Pro in the set falls at a single fixed offset. Green shading marks the high-affinity HRM (Cys-Pro-Trp), teal the lower-affinity Cys-Pro, and blue the histidine triad. High-affinity HRM calls are monophyletic on the Irr gene tree and contain exactly the motif-positive copies within *Nitrobacteraceae*. Two copies (of 58) fall below the landmark threshold and are scored "not assessable" rather than absent. *n* = 58 Irr copies from 50 genomes; 2 not assessable. TIE-1 rows are red: ***irrA1*** (Rpal_0428/RPAL_RS02160), *fab* locus, HRM$^{+}$; ***irrA2*** (Rpal_2583/RPAL_RS12765), alternative locus, HRM$^{-}$. See **Dataset S09** for the underlying data presented in tabular format.

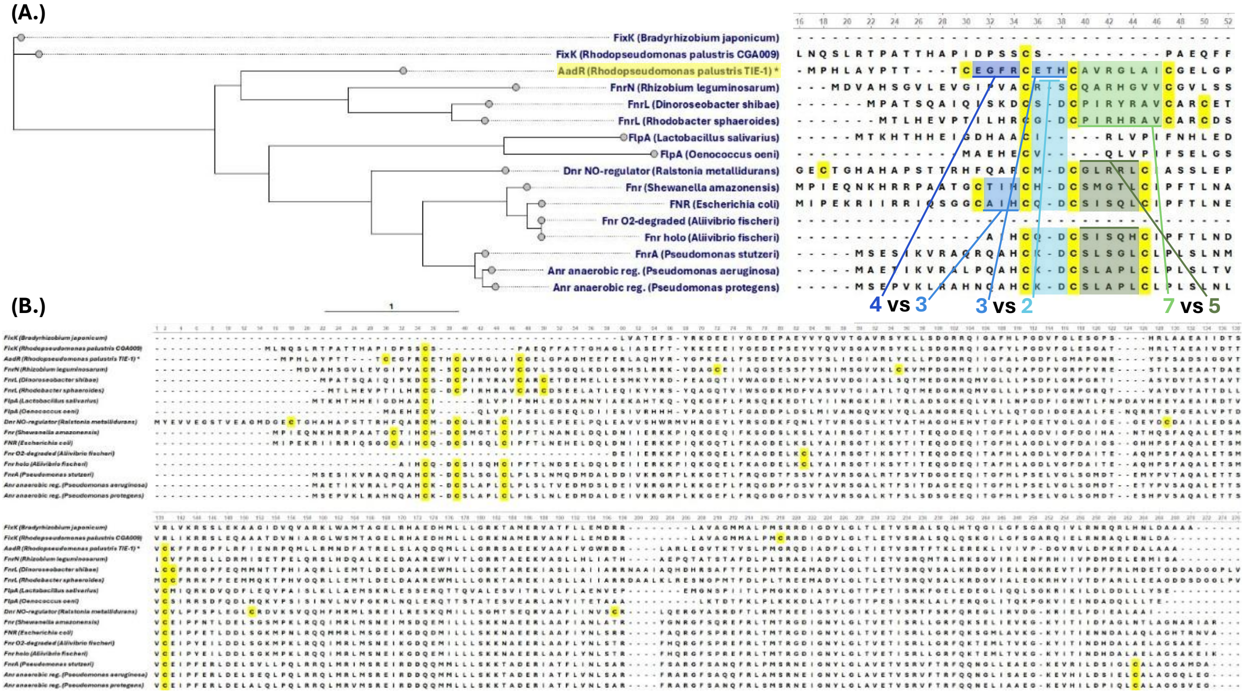


**Figure S16: Conserved cysteine residues in FNR-family regulators coordinate the binding of a [4Fe-4S] cluster (or other metal cofactor) to enable sensing oxygen, iron, NO, etc.**

**(A.)** Gene tree of selected FNR-family regulators across proteobacteria, and alignment of the N-terminal cysteine-rich domain (4 C) that helps coordinate a [4Fe-4S] cluster in regulators that possess it. Differences in AA spacer length between these aligned sequences are called out and colour coded. **(B.)** Full alignment of selected FNR-family regulators. Note the additional conserved cysteine at position 140 (as labelled here), which— while distant in primary structure— is close in secondary structure, completing the [4Fe-4S]-binding sensor domain of the regulator. A previous study (Green *et al.* 1993,[72]) identified that the second, third, fourth Cs in the domain (numbered 35, 38, and 44 in the above alignment here) are essential to FNR function in *E.c.* FNR, as is the distant C (numbered 140 here), while the first C (numbered 30 here) is dispensable.

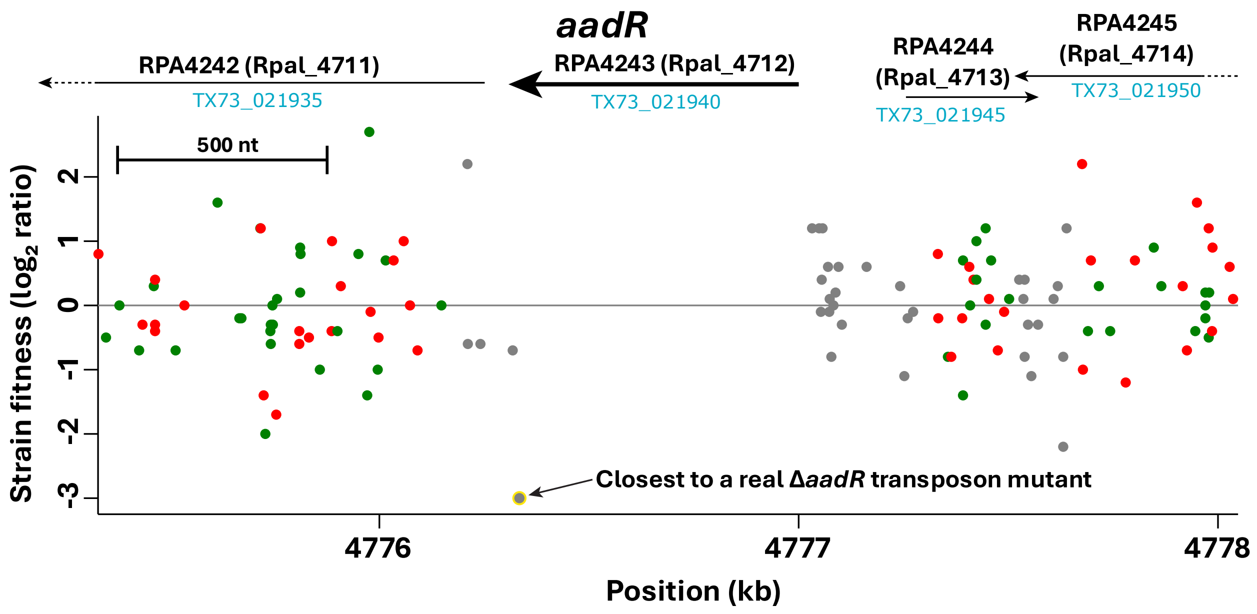


**Figure S17. Growth capability of strains produced through a transposon mutagenesis and fitness assay relative to the parental wild-type *R. palustris* CGA009.[73,74] No successful Δ*aadR* insertion mutants made it into their dataset.**

Green and red points show positive and negative strand insertions, respectively. Grey points are insertions near the edge of a gene (first or last 10%) or between genes. A positive strain fitness indicates superior growth of the mutant strain relative to wild-type, while a negative fitness indicates inferior mutant fitness (for full details, please see reference[73]). In the same dataset, *fixK* and *fixJ* show similar patterns, highlighting the importance of a functional AadR to successful growth (and evidencing the difficulty of producing a clean Δ*aadR* deletion strain, even aerobically and with kanamycin selection.[74] Data from: https://fit.genomics.lbl.gov/cgi-bin/strainTable.cgi?orgId=RPal_CGA009&locusId=TX73_021940&expName=set4S42

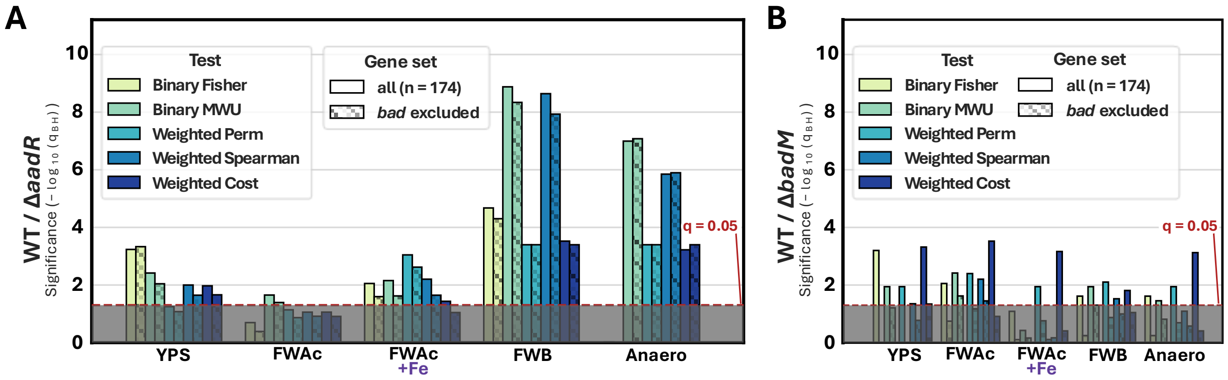


**Figure S18. Robustness of the AadR-dependent iron-requiring gene enrichment to statistical test and to exclusion of the bad operon.**

Enrichment of the iron-requiring gene set (n = 174 genes) among genes differentially expressed between Δ*aadR* and wild-type TIE-1 **(A)** or Δ*badM* and wild-type TIE-1 **(B)** were evaluated with five statistical tests: Binary Fisher's exact test, Binary Mann–Whitney U (MWU), and PERMANOVA, weighted Spearman and weighted iron-cost tests (the same tests as in **Figure 3E**). Each test was run on the both the full iron-requiring gene set (solid bars) and on the set with the *badDEFG* and *badAB* ("*bad* operon") genes excluded (hatched bars). Bars are grouped by growth condition (YPS, FWAcetate, FWAc+Fe, FWB, and all anaerobic), as in Figure 3. Significance is plotted as $-\log_{10}$ of the Benjamini–Hochberg-adjusted *q*-value; the dashed line marks $q = 0.05$ and the grey band below it denotes non-significance. Enrichment that reached significance for the full gene set was largely lost when the bad operon genes were excluded, indicating that these iron-requiring bad genes contribute most of the enrichment signal. Underlying data are in **SI Dataset S03**.

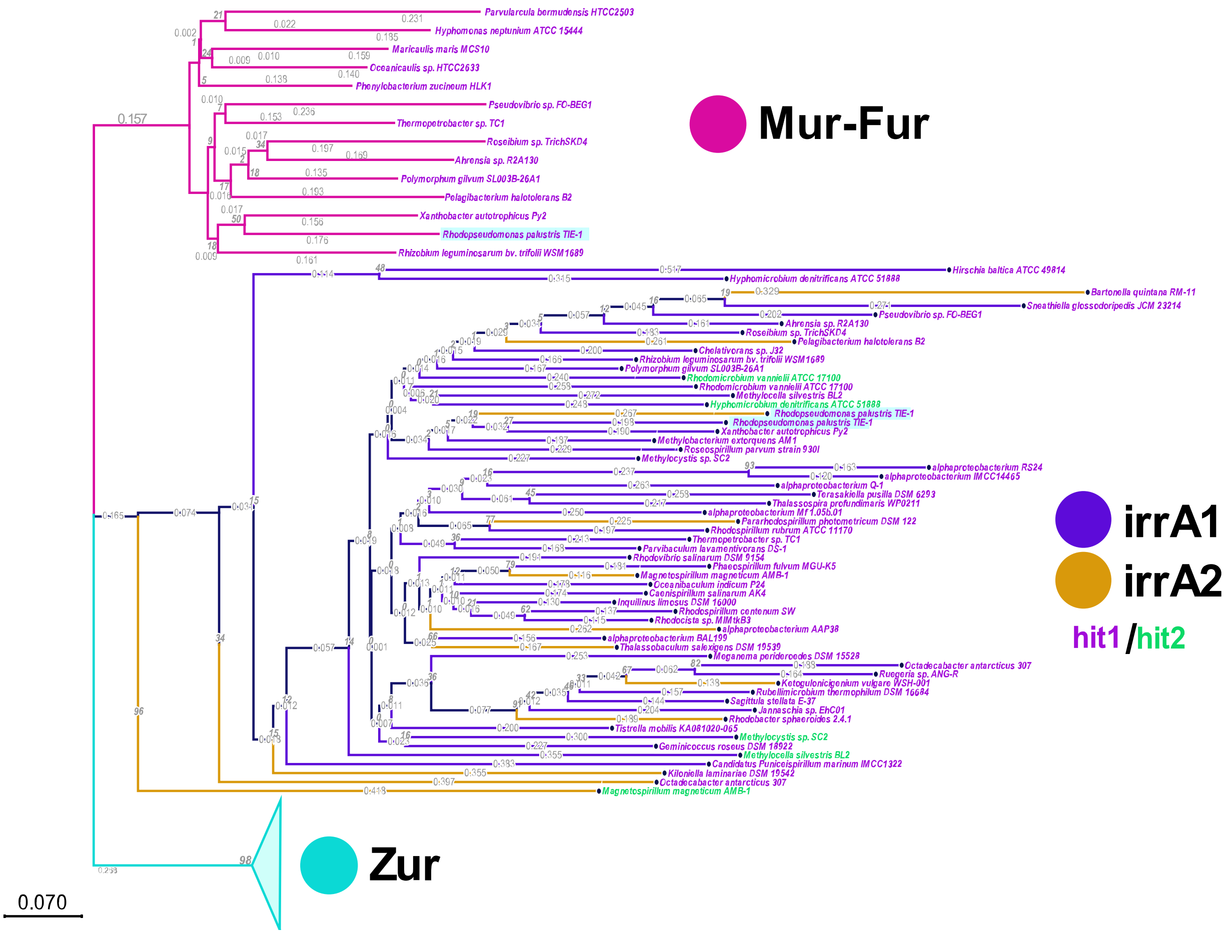


**Figure S19. Phylogeny of Fur-family regulators in diverse α-proteobacteria**

Fur-family regulators were identified across diverse alphaproteobacterial genomes using tBLASTn,[47] coloured here by their best match to each TIE-1 "probe" sequence (Mur, Zur, IrrA2, or IrrA1). Blue branches (Zur) and pink branches (Mur) are partly/completely collapsed, and clustered independently from the Irr branches. Gold (best match to IrrA2) and purple (best match to IrrA1) branches were interspersed with one another, indicating that sequences closer to each paralog were dispersed across alphaproteobacterial lineages. Names in green are genomes where multiple good BLAST hits on the same chromosome (indicating probable paralogs) were recovered. Overall, this supports the hypothesis that Irr paralogs in diverse alphaproteobacteria are the result of multiple independent gene duplication events.[71] Unrooted Maximum-Likelihood tree built from 100 bootstraps. The list of diverse alphaproteobacterial genome accessions used to build the tree was from Hördt *et al.* (2020).[47]

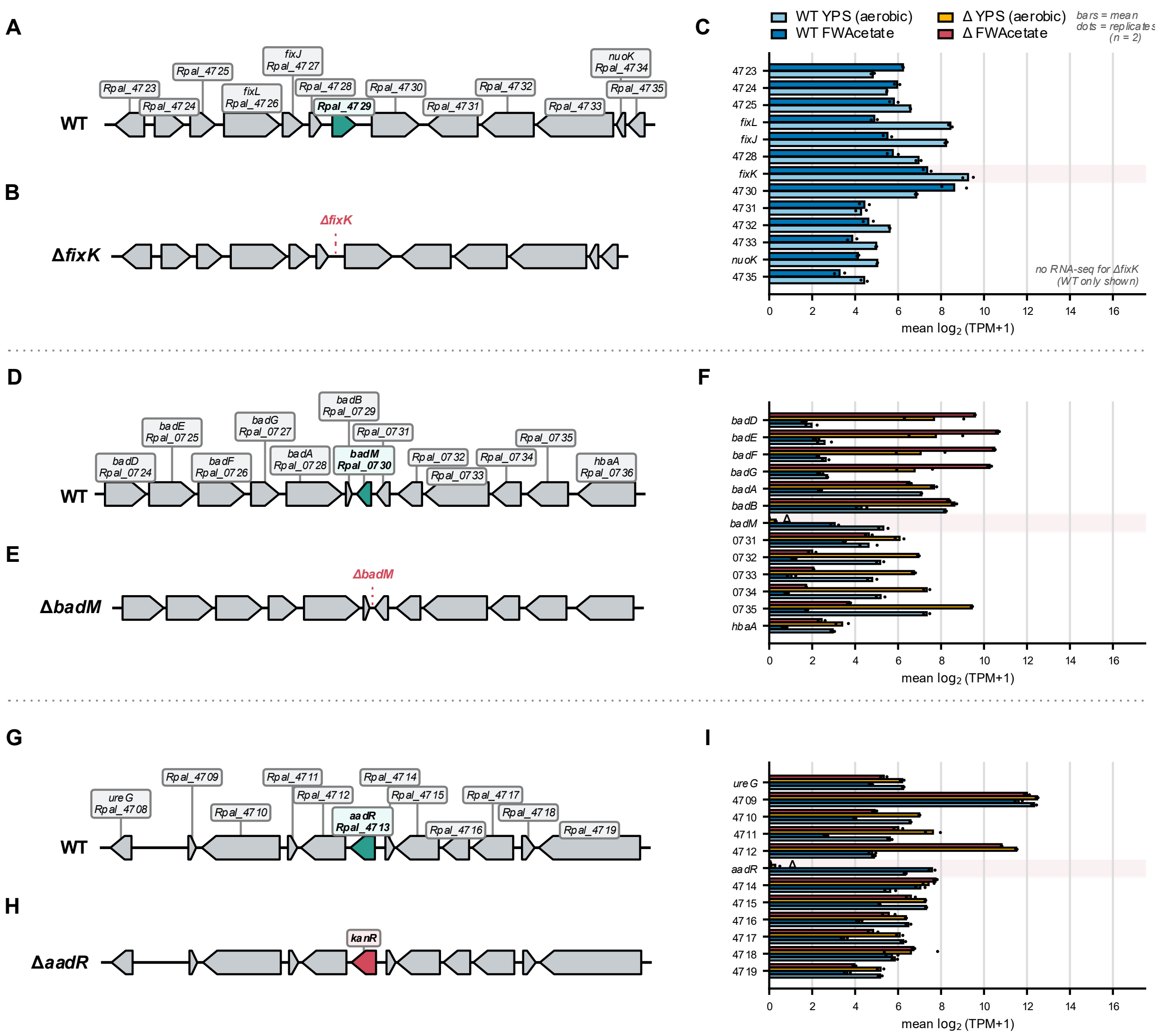


**Figure S20. Gene locus neighbourhoods of mutated regulators with relative expression of surrounding genes in transcripts per million (TPM).**

For each regulator, genes upstream and downstream are shown in wild-type *Rhodopseudomonas palustris* TIE-1 (**A**, **D**, **G**) and in the corresponding regulator deletion mutant strain (**B**, **E**, **H**), drawn to genomic scale from the TIE-1 reference annotation (GCF_000020445.1). Gene arrows indicate coding strand, and each is labelled with its gene symbol (where assigned) and locus tag. Deleted genes are highlighted in teal in the wild-type panels. Markerless deletions of *fixK* (Rpal_4729) and *badM* (Rpal_0730), and in-place replacement of *aadR* (Rpal_4713) with a kanamycin-resistance cassette (*kanR*, red). (**C**, **F**, **I**) Expression of each neighbouring gene as $\log_2(\text{TPM} + 1)$, in wild-type and the relevant deletion strain on aerobic chemoheterotrophy (YPS) and anaerobic photoheterotrophy (FWAcetate). Bars display mean expression and dots show individual biological replicate values ($n = 2$ per condition). RNA-seq was not performed for Δ*fixK*, so panel **C** shows wild-type values only. TPM values were computed from the raw-count matrix (**SI Dataset S07**) using reference CDS lengths. The deleted regulators read at or near zero in their own deletion strains (*aadR* and *badM*), confirming the deletions transcriptionally.

**SI Datasets and Software:**

**Dataset S01 (separate file).** Unified *Rhodopseudomonas palustris* TIE-1 – CGA009 feature table, mapping locus tags between the two strains (pairwise reciprocal best-hit BLAST) and between the GenBank and RefSeq versions of the genomes.

**Dataset S02 (separate file).** Unified Counts-Per-Million (CPM) transcriptomics dataset, as used in the analysis for **Figure 2A**.

**Dataset S03 (separate file).** Fe-requiring and Fe-related genes analysis spreadsheet. Contains rationales, lists, and calculations used to produce **Figure** 3.

**Dataset S04 (separate file).** Unified table of all growth (and Fe(II) oxidation) curves under each growth condition, used in the plotting of growth data in this study (**Figs. 1A-B,D-E, 4C-D, S4-S9, S10D**).

**Dataset S05 (separate file).** Exported data on *aadR* from the FitnessBrowser transposon mutagenesis and mutant relative fitness dataset in *R. palustris* CGA009.

**Dataset S06 (separate file).** *fixK* / *aadR* / Fur-family regulator iron response differential expression analysis spreadsheet, used for some data analysis/processing in **Figure 2** and **Figure 3.**

**Dataset S07 (separate file).** Complete set of Transcripts-Per-Million (TPM) matrices output by StringTie, as exported from KBase (see **Methods**). Provided here for reproducibility of our results.

**Dataset S08 (separate file).** Modified Gompertz logistic model fit parameters and calculated values (mu, lag, doubling time, maximum $OD_{660}$) from model fits on growth data in R (see **SI Text section 1.4**).

**Dataset S09 (separate file).** List of single-copy orthologues, full genome names and accession numbers on NCBI, and other metadata used to create **Figure 5**.

**Software S1 (separate file).** “SI_Software_S1__tie-1_*aadR*_paper_analysis_code-main.zip”. Contains python scripts used for analysis/visualisation of data, included for reproducibility of our results. This is a .zip archived frozen backup of the repository hosted on GitHub.

**SI Tables:**

**Table ST1.** Genes predicted to be regulated by FixK/AadR in *R. palustris* TIE-1. The TIE-1 genome was scanned for putative AadR/FixK binding site motifs using FIMO (MEME-suite).[67,68] Rows highlighted in yellow are genes whose locus tags are expressly reported in Bose and Newman (2011) as ones thought to be under regulation of FixK and whose transcripts have significantly greater expression during anaerobic vs. aerobic growth.[3]

| Score | E-value | Sequence | Gene | Name | Offset | Gene Description |
|---|---|---|---|---|---|---|
| 16.28 | 3.62E-10 | agtTTGATCtgGATCAAacg | Rpal_1206 | Rpal_1206 | -151 | hypothetical protein |
| | | | Rpal_1207 | COG1695 | -173 | transcriptional regulator, PadR-like family |
| 15.44 | 4.03E-09 | taaTTGATCtaGATCAAgcg | Rpal_4382 | Rpal_4382 | -80 | hypothetical protein |
| 15.12 | 8.19E-09 | gatTTGATttgGATCAAatg | Rpal_4994 | COG-OmpW | -98 | OmpW family protein |
| 14.76 | 1.68E-08 | gaaTTGATCaaGgTCAAacc | Rpal_4720 | COG3620 | -161 | CBS domain containing membrane protein |
| | | | Rpal_4721 | COG2187 | -149 | Uncharacterized protein conserved in bacteria |
| 14.2 | 4.53E-08 | aatTTGATCcgcATCAAatg | Rpal_0724 | COG-HgdB | -120 | benzoyl-CoA reductase, subunit C |
| **14.09** | **5.42E-08** | **tatTTGATCgaGATCAAtta** | **Rpal_0817** | **Rpal_0817** | **-232** | **cytochrome C family protein *(pioA)*** |
| 13.87 | 7.62E-08 | cgaTTGATCcgcATCAAacc | Rpal_1868 | COG-OsmY | -114 | transport-associated |
| 13.85 | 7.85E-08 | cggTTGATCcaGcTCAAagc | Rpal_3316 | COG-AcrR | -66 | transcriptional regulator, TetR family |
| 13.44 | 1.42E-07 | cgcTTGAgCtgGATCAAccg | Rpal_1079 | Rpal_1079 | -112 | putative exported protein of unknown function |
| | | | Rpal_1080 | Rpal_1080 | -237 | putative exported protein of unknown function |
| 12.99 | 2.61E-07 | gttTTGATCcaGAgCAAaga | Rpal_0261 | Rpal_0261 | -191 | cobalamin B12-binding |
| 12.99 | 2.61E-07 | gctTTGAgCcaGAaCAAacc | Rpal_1431 | COG-CaiD | -197 | enoyl-CoA hydratase |
| | | | Rpal_1432 | COG-TolQ | -174 | tonB-system energizer ExbB |
| 12.87 | 3.06E-07 | cgcTTGAcCttGcTCAAgcc | Rpal_4722 | COG-PhaC | -238 | Poly-beta-hydroxybutyrate polymerase domain protein |
| 12.79 | 3.38E-07 | ggcTTGATgaaGATCAAtgc | Rpal_0922 | COG-BioF | -169 | 5-aminolevulinate synthase |
| 12.76 | 3.48E-07 | gatTTGAgCtaGAaCAAtgc | Rpal_1713 | Rpal_1713 | -73 | hypothetical protein |
| 12.68 | 3.85E-07 | cgtTTGATCgaaATCAAaat | Rpal_0020 | COG-CcoN | -79 | cytochrome c oxidase, cbb3-type, subunit I |
| 12.57 | 4.38E-07 | tcaTTGATCtaccTCAAata | | | | Not near gene start |
| **12.41** | **5.36E-07** | **cggTTGATCaaGATCAAcag** | **Rpal_1280** | **COG-Crp** | **-78** | **transcriptional regulator, Crp/Fnr family *(Rpal_1280)*** |
| 12.28 | 6.16E-07 | cctTTGATtcaaATCAAgtt | Rpal_2130 | COG4235 | -157 | cytochrome c-type biogenesis protein CcmI |
| 12.26 | 6.35E-07 | ggcTTGATgcgGcTCAAgcg | | | | Not near gene start |
| 12.22 | 6.67E-07 | tgtTTGATgccGATCAAgct | | | | Not near gene start |
| 12.1 | 7.60E-07 | cgtTTGATtttGcTCAAtcg | Rpal_4121 | Rpal_4121 | -120 | putative bacterioferritin |
| 12.01 | 8.40E-07 | ttgTTGAcgcaGATCAAacg | Rpal_1864 | COG1032 | -71 | magnesium-protoporphyrin IX monomethyl ester anaerobic oxidative cyclase |
| 11.98 | 8.65E-07 | tccTTGATgcacATCAAacg | Rpal_2583 | COG-Fur | -188 | ferric uptake regulator, Fur family |
| 11.86 | 9.86E-07 | gctTTGATgcgcgTCAAatg | Rpal_5202 | COG-PorA | -245 | pyruvate flavodoxin/ferredoxin oxidoreductase domain protein |
| 11.78 | 1.08E-06 | ggaTTGAgacaGATCAAtcc | Rpal_4728 | COG-CheY | -92 | response regulator receiver protein |
| 11.69 | 1.19E-06 | catTTGAcgcacgTCAAata | Rpal_1198 | Rpal_1198 | -96 | protocatechuate 4,5-dioxygenase subunit alpha |
| 11.69 | 1.18E-06 | cgaTTGATgcaaATCAAtta | Rpal_4717 | COG-Ttg2B | -100 | protein of unknown function DUF140 |
| | | | Rpal_4718 | Rpal_4718 | -70 | hypothetical protein |
| 11.67 | 1.20E-06 | cggTTGATgcaGcTCAAtcg | | | | Not near gene start |
| 11.62 | 1.28E-06 | agaTTGAcgtgGATCAAcga | Rpal_1860 | COG-GloA | -79 | Glyoxalase/bleomycin resistance protein/dioxygenase |
| 11.57 | 1.34E-06 | agcTTGATCcatcTCAAtcc | | | | Not near gene start |
| 11.51 | 1.43E-06 | agcTTGAcacaGATCAAggt | Rpal_4713 | COG-Crp | -127 | transcriptional regulator, Crp/Fnr family |

**Table ST1 - *continued*.**

| | | | | | | |
|---|---|---|---|---|---|---|
| | | | Rpal_4714 | COG-CccA | -87 | conserved hypothetical protein; putative cytochrome c |
| 11.46 | 1.49E-06 | tggTTGAcgcgGATCAActc | Rpal_4743 | COG-NuoA | -130 | NADH-ubiquinone/plastoquinone oxidoreductase chain 3 |
| 11.41 | 1.58E-06 | cgcTTGAgCcatgTCAAacg | Rpal_1903 | COG-PaaJ | -111 | acetyl-CoA acetyltransferase |
| | | | Rpal_1904 | COG-AcrR | -40 | transcriptional regulator, TetR family |
| 11.4 | 1.59E-06 | caaTTGAgCtgtATCAAaac | Rpal_1861 | Rpal_1861 | -200 | hypothetical protein |
| 11.35 | 1.67E-06 | gcaaTGAgCtgGATCAAacc | Rpal_1479 | Rpal_1479 | -164 | hypothetical protein |
| 11.35 | 1.67E-06 | cttTTGAcCttGcTCAAtat | Rpal_1691 | COG4949 | -24 | Uncharacterized membrane-anchored protein conserved in bacteria |
| | | | Rpal_1692 | COG-HemC | -246 | porphobilinogen deaminase |
| 11.32 | 1.73E-06 | caaTTGcgCttGATCAAacg | Rpal_4083 | tnaA | -90 | tryptophanase |
| | | | Rpal_4084 | COG-ErfK | -200 | ErfK/YbiS/YcfS/YnhG family protein |
| 11.14 | 2.07E-06 | ctcTTGAcCtgagTCAAgga | Rpal_2012 | Rpal_2012 | -50 | hypothetical protein |
| 11.11 | 2.11E-06 | ggtTTGATttgGcaCAAaac | Rpal_5117 | COG-FeoA | -140 | FeoA family protein |
| | | | Rpal_5118 | Rpal_5118 | -241 | hypothetical protein |
| 11.08 | 2.17E-06 | gtgTTGAgCagGATCAAgga | Rpal_4806 | COG-FusA | -130 | elongation factor G |
| 11.04 | 2.26E-06 | aggTTGAggaaGATCAAttc | | | | Not near gene start |
| 11.04 | 2.26E-06 | gctTTGATgccGgTCAAccg | Rpal_3851 | Rpal_3851 | -133 | hypothetical protein |
| 10.95 | 2.47E-06 | gtcTTGATCttGATCAtgcc | Rpal_2490 | Rpal_2490 | -68 | hypothetical protein |
| | | | Rpal_2491 | COG-FtsJ | -149 | ribosomal RNA methyltransferase RrmJ/FtsJ |
| 10.92 | 2.55E-06 | gagTTGATCtgcggCAAatg | Rpal_0700 | Rpal_0700 | +87 | hypothetical protein |
| 10.91 | 2.55E-06 | gccTTGtTgcaGATCAAtca | Rpal_0174 | COG-AtpH | -212 | F0F1 ATP synthase subunit delta |
| 10.89 | 2.63E-06 | gcaTTGtgCcgGATCAAttg | Rpal_1810 | Rpal_1810 | -90 | hypothetical protein |
| | | | Rpal_1811 | Rpal_1811 | -130 | Rpal_1811 |
| 10.83 | 2.76E-06 | acaTTGAcgcgcATCAAatg | Rpal_4625 | COG-DeoA | -126 | thymidine phosphorylase |
| 10.82 | 2.79E-06 | aaaTTGATCcgGcgCAAtgc | Rpal_0995 | Rpal_0995 | -72 | hypothetical protein |
| 10.65 | 3.27E-06 | agcTTGATgtaacTCAAtac | Rpal_3850 | Rpal_3850 | -76 | hypothetical protein |
| 10.65 | 3.27E-06 | actTTGAcCggcgTCAAaga | Rpal_4709 | Rpal_4709 | -205 | hypothetical protein |
| 10.64 | 3.27E-06 | gacaTGAgCttGATCAAaca | Rpal_0283 | COG-LolA | +90 | outer membrane lipoprotein carrier protein LolA |
| 10.64 | 3.31E-06 | acaTTGATCagaATCAAcga | Rpal_1868 | COG-OsmY | -81 | transport-associated |
| 10.63 | 3.31E-06 | ccaTTGcgCtgGATCAAgga | Rpal_1865 | Rpal_1865 | -73 | hypothetical protein |
| | | | Rpal_1866 | Rpal_1866 | -168 | metallophosphoesterase |
| 10.55 | 3.58E-06 | cctTTGtTCctaATCAAgcc | Rpal_4996 | COG-Acs | -93 | AMP-dependent synthetase and ligase |
| 10.53 | 3.65E-06 | tgtTTGATCtgccgCAAgcg | Rpal_3439 | Rpal_3439 | -124 | hypothetical protein |
| 10.51 | 3.69E-06 | acaTTGATCtgGcgCAAccg | Rpal_4955 | Rpal_4955 | -134 | protein of unknown function DUF395 YeeE/YedE |
| 10.42 | 4.04E-06 | tctTTGATCagGcgCAAaga | Rpal_1001 | COG-UbiD | -67 | UbiD family decarboxylase |
| | | | Rpal_1002 | COG3154 | -68 | Putative lipid carrier protein |
| 10.41 | 4.08E-06 | gatTTGActgcGgTCAAttc | Rpal_2583 | COG-Fur | -70 | ferric uptake regulator, Fur family |

589

**Table ST2.** Growth characteristics of *ΔfixK* and wild-type TIE-1 on YP MOPS with 1 mM succinate (YPS), freshwater with succinate (FWS), acetate (FWA), and benzoate (FWB). Data are compared with numbers reported by Bose and Newman (2011),.[3] In their hands both the *ΔfixK* and wild-type TIE-1 grew much faster, almost certainly due to the difference in growth chamber lighting conditions, but the ratio of defect in doubling time between *ΔfixK* and WT is similar.

| **Strain:** | **ΔfixK** | | **WT** | | Ratio of Δ*fixK* / WT | |
|---|---|---|---|---|---|---|
| Metric | Doubling Time (hours) | Lag Time (hours) | Doubling Time (hours) | Lag Time (hours) | Doubling Time (hours) | Lag Time (hours) |
| — This Study | | | | | | |
| YPS (aerobic) | 4.11 ± 0.9 | 11.27 ± 3.13 | 4.84 ± 0.7 | 9.66 ± 1.64 | 0.9 | 1.1 |
| FWS (1 mM) | 26.60 ± 2.12 | 215.45 ± 3.37 | 8.18 ± 2.59 | 48.72 ± 4.3 | 2.0* | 5.2* |
| FWB (1 mM) | 17.04 ± 1.32 | 259.3 ± 2.1 | 13.08 ± 2.36 | 96.09 ± 4.0 | 2.1 | 2.7 |
| — Bose & Newman (2011)[3] | | | | | | |
| YPS (aerobic) | 6.1 ± 0.9 | 16.3 ± 1.3 | 3.8 ± 0.5 | 15.6 ± 2.2 | 1.6 | 1.0 |
| FWS (1 mM) | 9.4 ± 0.1 | 52.9 ± 0.8 | 3.7 ± 0.2 | 20.9 ± 1.4 | 2.5 | 2.5 |
| FWB (1 mM) | 7.4 ± 0.5 | 49.0 ± 3.6 | 4.0 ± 0.6 | 20.1 ± 3.1 | 1.9 | 2.4 |

**Table ST3.** Δ*aadR* and wild-type TIE-1 rates of Fe(II) oxidation and minimum amount of Fe(II) remaining in the growth medium for mixotrophic growth.

| Strain | Condition | Slope | $[Fe_{(II)}]_{minimum}$ |
|---|---|---|---|
| WT | FWAc+Fe | -6.4 ± 0.4 | 2.84 ± 0.01 |
| Δ*aadR* | FWAc+Fe | -5.7 ± 0.3 | 4.47 ± <0.01 |
| WT | FWMal+Fe | -3.4 ± 0.2 | 1.26 ± 0.04 |
| Δ*aadR* | FWMal+Fe | -1.6 ± 0.1 | 3.98 ± 0.03 |

**Table ST4.** Table of strains and plasmids used in this study.

| *Strain/Plasmid* | *Label* | *Genotype/Phenotype* | *Reference* |
|---|---|---|---|
| ***R. palustris strains*** | | | |
| *Wild-type TIE-1* | AB415 | Wild-type *Rhodopseudomonas palustris* TIE-1 | [75] |
| *ΔaadR* | AB209 | Δ*aadR* mutant of TIE-1 (*kanR* marker) | This study |
| *ΔaadRΔfixK* | AB207 | Δ*aadR*Δ*fixK* double mutant of TIE-1 | This study |
| *ΔaadRΔbadM* | AB208 | Δ*aadR*Δ*badM* double mutant of TIE-1 | This study |
| *ΔfixKΔbadM* | AB185 | Δ*fixK*Δ*badM* double mutant of TIE-1 | This study |
| *ΔfixK* | AB10 | Δ*fixK* single markerless deletion mutant of TIE-1 | [3] |
| *ΔbadM* | AB184 | Δ*badM* single markerless deletion mutant of TIE-1 | This study |
| *ΔaadRΔfixKΔbadM* | AB206 | Triple mutant of all three regulators (ΔΔΔ) | This study |
| *Ppio-ΔBS1* | AB12 | TIE-1 with *pioABC* promoter FNR-site1 (FixK Box) deleted | This study |
| *Ppio-ΔBS2* | AB18 | TIE-1 with *pioABC* promoter FNR-site2 deleted | This study |
| *ΔaadR complement* | AB215 | Δ*aadR* carrying pSRKGM:P*lac-aadR* | This study |
| *ΔbadM complement* | AB216 | Δ*badM* carrying pSRKGM:P*lac-badM* | This study |
| *ΔfixK complement* | AB21 | Δ*fixK* carrying pSRKGM:P*lac-fixK* | [3] |
| *Empty vector comp.* | AB114 | TIE-1 carrying pSRKGM plasmid | This study |
| ***E. coli strains*** | | | |
| *NEB-10* | NEB-10 | Plasmid holding strain, derivative of DH10B | New England Biolabs, Cat#c3019 |
| *WM6026* | ABB9 | DAP auxotroph mating strain, derivative of B2155 | [76] |
| ***Plasmids*** | | | |
| | pSRKGM | Expression vector used in complement strains | [77] |
| | pJQ200KS | Suicide plasmid used in strain construction | [78] |
| | pAB363 | *fixK* plasmid (in pSRKGM) | [3] |
| | pAB873 | *aadR* complementation plasmid (in pSRKGM) | This study |
| | pAB874 | *badM* complementation plasmid (in pSRKGM) | This study |

**Table ST5.** Primer sequences used for constructs made and PCR checks performed in this study.

| Primer Name | Purpose | Sequence |
|---|---|---|
| ***Cloning primers*** | | |
| Rp730_Up_FP | *ΔbadM* (upstr.) | agagGAGCTCAGGAGGCGATCTACCAGATG |
| Rp730_Up_RP | *ΔbadM* (upstr.) | agagTCTAGACAATCTTCATGTTGGTCTCGACCAC |
| Rp730_Dn_FP | *ΔbadM* (down) | agagTCTAGACCCTCAACCGAACTTTCAGTC |
| Rp730_Dn_RP | *ΔbadM* (down) | cagaGGATCCGTCATTTCGGCATCGACATC |
| *aadR*_HiFi_insert_FW | *aadR* compl. | GATAACAATGCCAGGAGAGTTCAGTCTGG |
| *aadR*_HiFi_insert_RV | *aadR* compl. | TTGCTAGCGCGTTTCAGGCCGCG |
| *aadR*_HiFi_vector_FW | *aadR* compl. | GAAACGCGCTAGCAATTCGAAAGCAAATTCGAC |
| *aadR*_HiFi_vector_RV | *aadR* compl. | CTCTCCTGGCATTGTTATCCGCTCACAATTCCAC |
| KanR_HiFi_insert_FW | *ΔaadR* cloning | ggagagttcagtctggcGACCGCATGGAGTGAT-GGCTGCATTGACATAAGCCTGTTCGGTTCG |
| KanR_HiFi_insert_RV | *ΔaadR* cloning | attccgggcggatgcgcGGCCGAAAGTGAAGCGTTCTGC-ATCAGAAGAACTCGTCAAGAAGGCGATAGAAGGCG |
| BS1-BS2_1Kb-Up_Fw_SacI* | P*pio-ΔBS1/2* cloning* | atgagatGAGCTCGAGATTCACGTCACCTCGATCTTC |
| BS1_up_BS1_Kb_Rv_SacI | P*pio-ΔBS1*cloning | tagctcACTAGTTTATTCACATCTGTTAGCATACGCAGCAGA |
| BS2_up_BS2_Kb_Rv_SacI | P*pio-ΔBS2* cloning | tagtctACTAGTTCAAATTGCCCTGTGCGCAATGCAG |
| ***Check primers*** | | |
| *aadR*_out_check_FW | *ΔaadR* check primer | CTGCCTTGACGAATTGCCGAC |
| *aadR*_out_check_RV | *ΔaadR* check primer | CGCTGAGCACGATGAAGAAGACT |
| *aadR*_in_check_ FW | *ΔaadR* check primer | ATCCGACGACGACTTGCGAG |
| *aadR*_in_check_ RV | *ΔaadR* check primer | ATCCGACGACGACTTGCGAG |
| 730_out_check_FW | *ΔbadM* check primer | TTTTCACTCAGGCGGGGGT |
| 730_out_check_RV | *ΔbadM* check primer | GTGTGTCCGAACCAGGCGA |
| fixK_in_check_FW | *ΔfixK* check primer | CAAGAAGGAAGAGGAAATCTACGGCG |
| fixK_out_check_FW | *ΔfixK* check primer | TCCTGTTAAGACATCGGTTTTCCCCA |
| fixK_in_check_RV | *ΔfixK* check primer | AGCAGCATGTGGTCTTCGGC |
| fixK_out_check_RV | *ΔfixK* check primer | TTTCGGGGTGGAGCTTCTGGG |
| kanR_in_check_RV | *ΔaadR* check primer | TGCTCTTCGTCCAGATCATCCTGAT |
| kanR_in_check_FW | *ΔaadR* check primer | GCACAACAGACAATCGGCTGC |
| GmR_check_RV | pSRKGM vector check | GATGAATGTCTTACTACGGAGCAAG |
| BS1-BS2_Fw_Check* | P*pio-ΔBS1/2* check* | ACAATCAGTCGACCCGACCAATCA |
| BS1_Rev_Check | P*pio-ΔBS1* check | TTGATCGAGATCAATTATTCAC |
| BS2_Rev_Check | P*pio-ΔBS2* check | AGCCACAATCTGTCATCA |

* *Because* P*pio*-BS1 *and* P*pio*-BS2 *are so close together, the same forward primer was used for cloning/checking both Δ*BS *mutants, with specificity granted through use of different reverse primers. Sequencing was used to confirm these mutants (as was used for gene mutants, see* ***Methods, Main Text)***

**Table ST6.** Locus tags of key genes mentioned in the text of this paper. For a comprehensive map of names and locus tags of every gene in the TIE-1 genome, please see **SI Dataset S01.**

| **Locus Tag** | **Name Used** | **Protein** | **Relevance to Present Study** |
|---|---|---|---|
| *Rpal_0428* | *irrA1* | Fur-family regulator | Iron responsive regulator; *irrA2* paralog[71] |
| *Rpal_0454* | *mur* | Fur-family regulator | $Mn^{2+}$ uptake regulator; possible Fur act.[55,57,59] |
| *Rpal_0518* | *zur* | Fur-family regulator | $Zn^{2+}$ uptake regulator |
| *Rpal_0730* | *badM* | Rrf2-family regulator | Regulator of the *bad* benzoate deg. operon (37–40) |
| *Rpal_0815* | *pioC* | High potential Fe-S | Involved in Fe(II) oxidation[3,79,80] |
| *Rpal_0816* | *pioB* | Outer membrane porin | Involved in Fe(II) oxidation[3,79,80] |
| *Rpal_0817* | *pioA* | Decahaem cytochrome | Involved in Fe(II) oxidation[3,79,80] |
| *Rpal_1280* | *Rpal_1280* | FNR-family regulator | Regulated by FixK[3] |
| *Rpal_2583* | *irrA2* | Fur-family regulator | Iron responsive regulator; *irrA1* paralog[71] |
| *Rpal_4713* | *aadR* | FNR-family regulator | Primary topic of the work; FNR-family regulator |
| *Rpal_4729* | *fixK* | FNR-family regulator | In negative feedback loop with AadR; FNR-family regulator |

**Table ST7.** Default interpretation of significance indicator stars. Stars in figures and tables indicate *p* or *q* bands of the following ranges unless otherwise specified in the caption.

| **Symbol** | ***p*-value interpretation** |
|---|---|
| ***For one-way and two-way ANOVA of means*** | |
| *ns* | $p>0.05$ |
| * | $p<0.05$ |
| ** | $p<0.01$ |
| *** | $p<0.001$ |
| **** | $p<0.0001$ |
| ***For fold-change / DESeq2 comparisons*** | |
| | $q>0.05$ |
| * | $0.05<q<0.0001$ |
| ** | $q<0.0001$ |

**Table ST8.** Iron-requiring genes in the TIE-1 proteome were ones with HMMer hits against the following PFAMs,[81] plus ones identified as coordinating Fe by MetalPredator.[82] For more information, including a list of all 174 genes we determined to be iron-requiring, please see **Dataset S03**.

| PFAM ID | PFAM NAME | # Uniq. | Locus Tags |
|---|---|---|---|
| **PF04055** | Radical SAM superfamily ([4Fe-4S]) | 29 | Rpal_0252, Rpal_0330, Rpal_0401, Rpal_0452, Rpal_0746, Rpal_1225, Rpal_1382, Rpal_1862, Rpal_1864, Rpal_2003, Rpal_2161, Rpal_2191, Rpal_2227, Rpal_2257, Rpal_2413, Rpal_2598, Rpal_2640, Rpal_2772, Rpal_2794, Rpal_2857, Rpal_2885, Rpal_2906, Rpal_2950, Rpal_4068, Rpal_4256, Rpal_4269, Rpal_4523, Rpal_4999, Rpal_5111 |
| **PF00034** | Cytochrome c | 14 | Rpal_0017, Rpal_0810, Rpal_0849, Rpal_1103, Rpal_1578, Rpal_1641, Rpal_1724, Rpal_2214, Rpal_2428, Rpal_4215, Rpal_4495, Rpal_4714, Rpal_4956, Rpal_4962 |
| **PF00111** | 2Fe-2S iron-sulfur cluster binding (ferredoxin) | 12 | Rpal_0737, Rpal_1085, Rpal_1770, Rpal_2078, Rpal_2139, Rpal_3292, Rpal_4141, Rpal_4285, Rpal_4326, Rpal_4477, Rpal_4738, Rpal_5147 |
| **PF00037** | 4Fe-4S dicluster domain (ferredoxin) | 11 | Rpal_0137, Rpal_0490, Rpal_0729, Rpal_0805, Rpal_1323, Rpal_1418, Rpal_3290, Rpal_4736, Rpal_5093, Rpal_5110, Rpal_5112 |
| **PF12838** | 4Fe-4S dicluster domain (Fer4_7) | 11 | Rpal_0016, Rpal_0490, Rpal_0805, Rpal_1205, Rpal_1323, Rpal_3290, Rpal_4736, Rpal_5093, Rpal_5110, Rpal_5112, Rpal_5202 |
| **PF13237** | 4Fe-4S dicluster domain (Fer4_10) | 7 | Rpal_0215, Rpal_0490, Rpal_1323, Rpal_3290, Rpal_4736, Rpal_5093, Rpal_5112 |
| **PF00175** | Oxidoreductase NAD-binding domain | 5 | Rpal_1085, Rpal_1766, Rpal_4141, Rpal_4232, Rpal_4285 |
| **PF00394** | Multicopper oxidase (Cu-oxidase) | 5 | Rpal_1040, Rpal_1091, Rpal_3312, Rpal_3727, Rpal_4623 |
| **PF00355** | Rieske [2Fe-2S] domain | 4 | Rpal_1175, Rpal_1208, Rpal_1384, Rpal_4139 |
| **PF01568** | Molybdopterin oxidoreductase Fe4S4 domain | 4 | Rpal_0384, Rpal_0805, Rpal_1532, Rpal_4710 |
| **PF13510** | 2Fe-2S iron-sulfur cluster binding (Fer2_4) | 4 | Rpal_0137, Rpal_0805, Rpal_3292, Rpal_4738 |
| **PF00115** | Cytochrome c/quinol oxidase subunit I (haem-copper; haem a/a3) | 3 | Rpal_0020, Rpal_0900, Rpal_1642 |
| **PF01512** | Respiratory-chain NADH dehydrogenase 51 kDa subunit (Fe-S) | 3 | Rpal_0804, Rpal_3293, Rpal_4739 |
| **PF01077** | Nitrite/sulfite reductase ferredoxin-like half domain ([4Fe-4S]+sirohaem) | 2 | Rpal_4231, Rpal_4694 |
| **PF01794** | Ferric reductase-like transmembrane component | 2 | Rpal_1216, Rpal_3123 |
| **PF04324** | BFD-like [2Fe-2S] binding domain | 2 | Rpal_4121, Rpal_5090 |
| **PF13186** | SPASM/twitch domain ([4Fe-4S], radical-SAM-associated) | 2 | Rpal_2161, Rpal_4523 |
| **MetalPredator-identified genes** | | 66 | Rpal_0199, Rpal_0240, Rpal_0346, Rpal_0457, Rpal_0474, Rpal_0502, Rpal_0520, Rpal_0602, Rpal_0739, Rpal_0745, Rpal_0803, Rpal_0959, Rpal_0960, Rpal_1152, Rpal_1154, Rpal_1162, Rpal_1169, Rpal_1229, Rpal_1247, Rpal_1417, Rpal_1608, Rpal_1620, Rpal_1622, Rpal_1623, Rpal_1688, Rpal_1710, Rpal_1711, Rpal_1712, Rpal_1740, Rpal_1789, Rpal_1796, Rpal_1933, Rpal_2138, Rpal_2350, Rpal_2591, Rpal_2592, Rpal_2597, Rpal_2607, Rpal_2608, Rpal_2742, Rpal_2744, Rpal_2745, Rpal_2748, Rpal_2886, Rpal_2887, Rpal_2888, Rpal_2907, Rpal_2908, Rpal_2909, Rpal_3196, Rpal_3295, Rpal_3299, Rpal_4121, Rpal_4255, Rpal_4397, Rpal_4740, Rpal_4742, Rpal_4751, Rpal_4784, Rpal_4978, Rpal_5083, Rpal_5089, Rpal_5091, Rpal_5097, Rpal_5098, Rpal_5109 |
| **Literature-identified genes** | | 14 | Rpal_0817, Rpal_0816, Rpal_0815, Rpal_5101, Rpal_5100, Rpal_5099, Rpal_1733, Rpal_1730, Rpal_1731, Rpal_0724, Rpal_0725, Rpal_0726, Rpal_0727, Rpal_0729 |

**Table ST9.** Manual checks/additions of genes to iron-related and/or iron-requiring sets (curated from the literature). Superscripts in the "notes" column refer to the paper citation for literature backing of additions.

| Locus Tag | RefSeq Tag | Type | # Fe atoms | Gene | Source |
|---|---|---|---|---|---|
| Rpal_0817 | RPAL_RS04115 | decahaem_c-type | 10 | *pioA* | [79] |
| Rpal_0816 | RPAL_RS04110 | outer_membrane_porin | 0 | *pioB* | [79] |
| Rpal_0815 | RPAL_RS04105 | HiPIP_[4Fe-4S] | 4 | *pioC* | [83] |
| Rpal_5101 | RPAL_RS25275 | [4Fe-4S]_dimer-bridging | 4 | *nifH* | [84] |
| Rpal_5100 | RPAL_RS25270 | FeMo-cofactor_+_P-cluster | 11 | *nifD* | [84] |
| Rpal_5099 | RPAL_RS25265 | P-cluster_[8Fe-7S] | 4 | *nifK* | [84] |
| Rpal_1733 | RPAL_RS08600 | [4Fe-4S] | 4 | *bchL* | [85] |
| Rpal_1730 | RPAL_RS08585 | [4Fe-4S] | 4 | *bchN* | [85] |
| Rpal_1731 | RPAL_RS08590 | [4Fe-4S] | 4 | *bchB* | [85] |
| Rpal_0724 | RPAL_RS03650 | ATPase_no_FeS | 0 | *badD* | [37,86] |
| Rpal_0725 | RPAL_RS03655 | [4Fe-4S]_bridging | 4 | *badE* | [37,86] |
| Rpal_0726 | RPAL_RS03660 | [4Fe-4S]_catalytic | 4 | *badF* | [37,86] |
| Rpal_0727 | RPAL_RS03665 | [4Fe-4S]_catalytic | 4 | *badG* | [37,86] |
| Rpal_0729 | RPAL_RS03675 | 2x[4Fe-4S]_ferredoxin | 8 | *badB* | [37,86] |

**Table ST10.** Iron-related genes set from the TIE-1 proteome. For the full lists of the 435 genes we determined to be iron-related, please see **Dataset S03**.

| Category | Number of Genes (includes overlap) | Genes (for the full set, see Dataset S03) |
|---|---|---|
| **FeGenie: iron_aquisition-heme_oxygenase** | 3 | Rpal_1727, Rpal_2417, Rpal_3698 |
| **FeGenie: iron_aquisition-heme_transport** | 60 | Rpal_0017, Rpal_0019, Rpal_0204, Rpal_0401, Rpal_0477, Rpal_0664, ... |
| **FeGenie: iron_aquisition-iron_transport** | 103 | Rpal_0050, Rpal_0057, Rpal_0102, Rpal_0103, Rpal_0134, Rpal_0191, ... |
| **FeGenie: iron_aquisition-siderophore_synthesis** | 140 | Rpal_0028, Rpal_0050, Rpal_0057, Rpal_0102, Rpal_0103, Rpal_0132, ... |
| **FeGenie: iron_aquisition-siderophore_transport** | 156 | Rpal_0050, Rpal_0057, Rpal_0090, Rpal_0102, Rpal_0103, Rpal_0132, ... |
| **FeGenie: iron_aquisition-siderophore_transport_potential** | 15 | Rpal_1308, Rpal_1309, Rpal_1432, Rpal_1433, Rpal_1434, Rpal_2220, ... |
| **FeGenie: iron_gene_regulation** | 17 | Rpal_0428, Rpal_0454, Rpal_0518, Rpal_1207, Rpal_1268, Rpal_1537, ... |
| **FeGenie: iron_oxidation** | 3 | Rpal_0810, Rpal_0816, Rpal_0817 |
| **FeGenie: iron_reduction** | 6 | Rpal_0525, Rpal_0817, Rpal_1232, Rpal_1294, Rpal_1707, Rpal_2356 |
| **FeGenie: iron_storage** | 4 | Rpal_1468, Rpal_1521, Rpal_3094, Rpal_4120 |
| **FeGenie: magnetosome_formation** | 5 | Rpal_1230, Rpal_2140, Rpal_3637, Rpal_4002, Rpal_5052 |
| **FeGenie: possible_iron_oxidation_and_possible_iron_red.** | 62 | Rpal_0026, Rpal_0030, Rpal_0176, Rpal_0185, Rpal_0215, Rpal_0270, ... |

**Table ST11.** α-proteobacterial genomes used to build the Figure 5 tree figure (type strain genomes from the Hördt et al. (2020) list,[47] plus select Rhodopseudomonas), with accession number and taxonomic assignment by GTDB and NCBI, listed in tip order counterclockwise from root.

| tip_order | short_name | organism | genome_accession | GTDB_order | NCBI_order | GTDB_family | NCBI_family | lineage_group |
|---|---|---|---|---|---|---|---|---|
| 1 | **Mpf.** | Mariprofundus ferrooxydans PV-1 | GCF_000153765.1 | Mariprofundales | Mariprofundales | Mariprofundaceae | Mariprofundaceae | outgroups |
| 2 | **Eco.** | Escherichia coli str. K-12 substr. MG1655 | GCF_000005845.2 | Enterobacterales | Enterobacterales | Enterobacteriaceae | Enterobacteriaceae | outgroups |
| 3 | **Rvg.** | Rubrivivax gelatinosus DSM 1709 | GCF_004340905.1 | Burkholderiales | Burkholderiales | Burkholderiaceae | Sphaerotilaceae | outgroups |
| 4 | **Sdl.** | Sideroxydans lithotrophicus ES-1 | GCF_000025705.1 | Burkholderiales | Nitrosomonadales | Gallionellaceae | Gallionellaceae | outgroups |
| 5 | **Gac.** | Gallionella capsiferriformans ES-2 | GCF_000145255.1 | Burkholderiales | Nitrosomonadales | Gallionellaceae | Gallionellaceae | outgroups |
| 6 | **Mam.** | Magnetococcus marinus MC-1 | GCF_000014865.1 | Magnetococcales | Magnetococcales | Magnetococcaceae | Magnetococcaceae | other alpha-orders |
| 7 | **Acc.** | Acidiphilium cryptum JF-5 | GCF_000016725.1 | Rhodospirillales | Acetobacterales | Acetobacteraceae | Acidocellaceae | other alpha-orders |
| 8 | **Rlg.** | Rhodopila globiformis DSM 161 | GCF_002937115.1 | Rhodospirillales | Acetobacterales | Acetobacteraceae | Acetobacteraceae | other alpha-orders |
| 9 | **Asb.** | Azospirillum brasilense Sp 7 | GCF_007827425.1 | Rhodospirillales | Rhodospirillales | Azospirillaceae | Azospirillaceae | other alpha-orders |
| 10 | **Pmm.** | Paramagnetospirillum magneticum AMB-1 | GCF_000009985.1 | Rhodospirillales | Rhodospirillales | Magnetospirillaceae | Magnetospirillaceae | other alpha-orders |
| 11 | **Rsr.** | Rhodospirillum rubrum ATCC 11170 | GCF_000013085.1 | Rhodospirillales | Rhodospirillales | Rhodospirillaceae | Rhodospirillaceae | other alpha-orders |
| 12 | **Noa.** | Novosphingobium aromaticivorans DSM 12444 | GCF_000013325.1 | Sphingomonadales | Sphingomonadales | Sphingomonadaceae | Sphingomonadaceae | other alpha-orders |
| 13 | **Cav.** | Caulobacter vibrioides NA1000 | GCF_000022005.1 | Caulobacterales | Caulobacterales | Caulobacteraceae | Caulobacteraceae | other alpha-orders |
| 14 | **Dis.** | Dinoroseobacter shibae DFL 12 = DSM 16493 | GCF_000018145.1 | Rhodobacterales | Rhodobacterales | Rhodobacteraceae | Roseobacteraceae | other alpha-orders |
| 15 | **Ces.** | Cereibacter sphaeroides 2.4.1 | GCF_003324715.1 | Rhodobacterales | Rhodobacterales | Rhodobacteraceae | Paracoccaceae | other alpha-orders |
| 16 | **Pad.** | Paracoccus denitrificans ATCC 19367 | GCF_004063735.1 | Rhodobacterales | Rhodobacterales | Rhodobacteraceae | Paracoccaceae | other alpha-orders |
| 17 | **Hyd.** | Hyphomicrobium denitrificans ATCC 51888 | GCF_000143145.1 | Hyphomicrobiales | Hyphomicrobiales | Hyphomicrobiaceae | Hyphomicrobiaceae | Hyphomicrobiales outgroup |
| 18 | **Rmv.** | Rhodomicrobium vannielii ATCC 17100 | GCF_016461745.1 | Hyphomicrobiales | Hyphomicrobiales | Rhodomicrobiaceae | Hyphomicrobiaceae | Hyphomicrobiales outgroup |
| 19 | **Der.** | Devosia riboflavina IFO13584 | GCF_000743575.1 | Hyphomicrobiales | Hyphomicrobiales | Devosiaceae | Devosiaceae | "& relatives" (Rhizobiaceae Branch) |
| 20 | **Rdo.** | Rhodobium orientis DSM 11290 | GCF_003258835.1 | Hyphomicrobiales | Hyphomicrobiales | Rhodobiaceae | Rhodobiaceae | "& relatives" (Rhizobiaceae Branch) |
| 21 | **Auc.** | Aurantimonas coralicida DSM 14790 | GCF_000421645.1 | Hyphomicrobiales | Hyphomicrobiales | Rhizobiaceae | Aurantimonadaceae | "& relatives" (Rhizobiaceae Branch) |
| 22 | **Snm.** | Sinorhizobium meliloti MABNR56 | GCF_037023865.1 | Hyphomicrobiales | Hyphomicrobiales | Rhizobiaceae | Rhizobiaceae | Rhizobiaceae (RirA+) |
| 23 | **Rzl.** | Rhizobium leguminosarum SM52 | GCF_004306555.1 | Hyphomicrobiales | Hyphomicrobiales | Rhizobiaceae | Rhizobiaceae | Rhizobiaceae (RirA+) |
| 24 | **Agf.** | Agrobacterium fabrum str. C58 | GCF_000092025.1 | Hyphomicrobiales | Hyphomicrobiales | Rhizobiaceae | Rhizobiaceae | Rhizobiaceae (RirA+) |
| 25 | **Neg.** | Neorhizobium galegae bv. orientalis str. HAMBI 540 | GCF_000731315.1 | Hyphomicrobiales | Hyphomicrobiales | Rhizobiaceae | Rhizobiaceae | Rhizobiaceae (RirA+) |
| 26 | **Mej.** | Mesorhizobium japonicum MAFF 303099 | GCF_000009625.1 | Hyphomicrobiales | Hyphomicrobiales | Rhizobiaceae | Phyllobacteriaceae | Rhizobiaceae (RirA+) |
| 27 | **Ama.** | Aminobacter aminovorans NCTC10684 | GCF_900445235.1 | Hyphomicrobiales | Hyphomicrobiales | Rhizobiaceae | Phyllobacteriaceae | Rhizobiaceae (RirA+) |
| 28 | **Bah.** | Bartonella henselae FDAARGOS_1462 | GCF_019930925.1 | Hyphomicrobiales | Hyphomicrobiales | Rhizobiaceae | Bartonellaceae | "& relatives" (Rhizobiaceae Branch) |
| 29 | **Bcm.** | Brucella melitensis BwIM_TUR_17 | GCF_002191655.1 | Hyphomicrobiales | Hyphomicrobiales | Rhizobiaceae | Brucellaceae | "& relatives" (Rhizobiaceae Branch) |
| 30 | **Bca.** | Brucella anthropi PBO | GCF_015326295.1 | Hyphomicrobiales | Hyphomicrobiales | Rhizobiaceae | Brucellaceae | "& relatives" (Rhizobiaceae Branch) |
| 31 | **Mre.** | Methylorubrum extorquens DM4 | GCF_000083545.1 | Hyphomicrobiales | Hyphomicrobiales | Beijerinckiaceae | Methylobacteriaceae | "Other " (Nitrobacteraceae Branch) |
| 32 | **Mil.** | Microvirga lotononidis HAMBI_3237 | GCF_034627025.1 | Hyphomicrobiales | Hyphomicrobiales | Beijerinckiaceae | Methylobacteriaceae | "Other" (Nitrobacteraceae Branch) |
| 33 | **Rba.** | Rhodoblastus acidophilus DSM 137 | GCF_900187365.1 | Hyphomicrobiales | Hyphomicrobiales | Beijerinckiaceae | Rhodoblastaceae | "Other " (Nitrobacteraceae Branch) |
| 34 | **Bei.** | Beijerinckia indica subsp. indica ATCC 9039 | GCF_000019845.1 | Hyphomicrobiales | Hyphomicrobiales | Beijerinckiaceae | Beijerinckiaceae | "Other " (Nitrobacteraceae Branch) |
| 35 | **Mcs.** | Methylocella silvestris BL2 | GCF_000021745.1 | Hyphomicrobiales | Hyphomicrobiales | Beijerinckiaceae | Beijerinckiaceae | "Other " (Nitrobacteraceae Branch) |
| 36 | **Blv.** | Blastochloris viridis ATCC 19567 | GCF_001402875.1 | Hyphomicrobiales | Hyphomicrobiales | Xanthobacteraceae | Blastochloridaceae | "Other " (Nitrobacteraceae Branch) |
| 37 | **Arc.** | Azorhizobium caulinodans ORS 571 | GCF_000010525.1 | Hyphomicrobiales | Hyphomicrobiales | Xanthobacteraceae | Xanthobacteraceae | "Other " (Nitrobacteraceae Branch) |
| 38 | **Xaa.** | Xanthobacter autotrophicus DSM 432 | GCF_005871085.1 | Hyphomicrobiales | Hyphomicrobiales | Xanthobacteraceae | Xanthobacteraceae | "Other " (Nitrobacteraceae Branch) |
| 39 | **Afc.** | Afipia carboxidovorans OM5 | GCF_000218565.1 | Hyphomicrobiales | Hyphomicrobiales | Xanthobacteraceae | Nitrobacteraceae | Nitrobacteraceae |
| 40 | **Niw.** | Nitrobacter winogradskyi Nb-255 | GCF_000012725.1 | Hyphomicrobiales | Hyphomicrobiales | Xanthobacteraceae | Nitrobacteraceae | Nitrobacteraceae |
| 41 | **Brd.** | Bradyrhizobium diazoefficiens USDA 110 | GCF_001642675.1 | Hyphomicrobiales | Hyphomicrobiales | Xanthobacteraceae | Nitrobacteraceae | Nitrobacteraceae |
| 42 | **Brj.** | Bradyrhizobium japonicum 5038 | GCF_013752735.1 | Hyphomicrobiales | Hyphomicrobiales | Xanthobacteraceae | Nitrobacteraceae | Nitrobacteraceae |
| 43 | **Rpr.** | **Rhodopseudomonas rhenobacensis DSM 12706** | GCF_014203125.1 | Hyphomicrobiales | Hyphomicrobiales | Xanthobacteraceae | Nitrobacteraceae | Nitrobacteraceae |
| 44 | **Rpi.** | **Rhodopseudomonas infernalis HC1** | GCF_022271405.1 | Hyphomicrobiales | Hyphomicrobiales | Xanthobacteraceae | Nitrobacteraceae | Nitrobacteraceae |
| 45 | **Rpp.** | **Rhodopseudomonas pseudopalustris DSM 123** | GCF_058222895.1 | Hyphomicrobiales | Hyphomicrobiales | Xanthobacteraceae | Nitrobacteraceae | Nitrobacteraceae |
| 46 | **Rp-GJ22** | **Rhodopseudomonas palustris GJ-22** | GCF_007005445.1 | Hyphomicrobiales | Hyphomicrobiales | Xanthobacteraceae | Nitrobacteraceae | Nitrobacteraceae |
| 47 | **Rp-XCP** | **Rhodopseudomonas palustris XCP** | GCF_003226555.1 | Hyphomicrobiales | Hyphomicrobiales | Xanthobacteraceae | Nitrobacteraceae | Nitrobacteraceae |
| 48 | **Rp-PP803** | **Rhodopseudomonas palustris PP803** | GCF_025811355.1 | Hyphomicrobiales | Hyphomicrobiales | Xanthobacteraceae | Nitrobacteraceae | Nitrobacteraceae |
| 49 | **Rp-TIE1** | **Rhodopseudomonas palustris TIE-1** | GCF_000020445.1 | Hyphomicrobiales | Hyphomicrobiales | Xanthobacteraceae | Nitrobacteraceae | Nitrobacteraceae |
| 50 | **Rp-CGA009** | **Rhodopseudomonas palustris CGA009** | GCF_000195775.2 | Hyphomicrobiales | Hyphomicrobiales | Xanthobacteraceae | Nitrobacteraceae | Nitrobacteraceae |

**SI References:**


1. Rey, F. E. & Harwood, C. S. FixK, a global regulator of microaerobic growth, controls photosynthesis in *Rhodopseudomonas palustris*. *Molecular Microbiology* **75**, 1007–1020 (2010).
2. Kuo, F.-S., Chien, Y.-H. & Chen, C.-J. Effects of light sources on growth and carotenoid content of photosynthetic bacteria *Rhodopseudomonas palustris*. *Bioresource Technology* **113**, 315–318 (2012).
3. Bose, A. & Newman, D. K. Regulation of the phototrophic iron oxidation (*pio*) genes in *Rhodopseudomonas palustris* TIE-1 is mediated by the global regulator, FixK. *Molecular Microbiology* **79**, 63–75 (2011).
4. Muzziotti, D., Adessi, A., Faraloni, C., Torzillo, G. & De Philippis, R. Acclimation strategy of *Rhodopseudomonas palustris* to high light irradiance. *Microbiological Research* **197**, 49–55 (2017).
5. VanDrisse, C. M. & Escalante-Semerena, J. C. Small-Molecule Acetylation Controls the Degradation of Benzoate and Photosynthesis in *Rhodopseudomonas palustris*. *mBio* **9**, e01895-18 (2018).
6. Haas, N. W. *et al.* PioABC-Dependent Fe(II) Oxidation during Photoheterotrophic Growth on an Oxidized Carbon Substrate Increases Growth Yield. *Applied and Environmental Microbiology* **88**, e00974-22 (2022).
7. Yin, L. *et al.* BIODEGRADATION OF CYPERMETHRIN BY *RHODOPSEUDOMONAS PALUSTRIS* GJ-22 ISOLATED FROM ACTIVATED SLUDGE. *Fresenius Environmental Bulletin* **21**, (2012).
8. Ritchie, R. J. The Use of Solar Radiation by the Photosynthetic Bacterium, *Rhodopseudomonas palustris*: Model Simulation of Conditions Found in a Shallow Pond or a Flatbed Reactor. *Photochemistry and Photobiology* **89**, 1143–1162 (2013).
9. Clark, C. D., De Bruyn, W. J., Brahm, B. & Aiona, P. Optical properties of chromophoric dissolved organic matter (CDOM) and dissolved organic carbon (DOC) levels in constructed water treatment wetland systems in southern California, USA. *Chemosphere* **247**, 125906 (2020).
10. Maurice, N., Pochet, C., Adouani, N. & Pons, M.-N. Role of Seasons in the Fate of Dissolved Organic Carbon and Nutrients in a Large-Scale Surface Flow Constructed Wetland. *Water* **14**, (2022).
11. Vähätalo, A. V., Wetzel, R. G. & Paerl, H. W. Light absorption by phytoplankton and chromophoric dissolved organic matter in the drainage basin and estuary of the Neuse River, North Carolina (U.S.A.). *Freshwater Biology* **50**, 477–493 (2005).
12. Fontecilla-Camps, J. C., Amara, P., Cavazza, C., Nicolet, Y. & Volbeda, A. Structure–function relationships of anaerobic gas-processing metalloenzymes. *Nature* **460**, 814–822 (2009).
13. Garcia, P. S. *et al.* An early origin of iron–sulfur cluster biosynthesis machineries before Earth oxygenation. *Nat Ecol Evol* **6**, 1564–1572 (2022).
14. Frey, P. A. & Reed, G. H. The Ubiquity of Iron. *ACS Chem. Biol.* **7**, 1477–1481 (2012).
15. Ems, T., St Lucia, K. & Huecker, M. R. Biochemistry, Iron Absorption. in *StatPearls* (StatPearls Publishing, Treasure Island (FL), 2026).
16. Bridgman, E. & Haas, K. Iron acquisition in bacteria: Siderophores. *Chemistry LibreTexts* https://chem.libretexts.org/Courses/Saint_Marys_College_Notre_Dame_IN/CHEM_342%3A_Bio-inorganic_Chemistry/Readings/Metals_in_Biological_Systems_(Saint_Mary's_College)/Iron_acquisition_in_bacteria%3A_Siderophores (2016).
17. Kramer, J., Özkaya, Ö. & Kümmerli, R. Bacterial siderophores in community and host interactions. *Nat Rev Microbiol* **18**, 152–163 (2020).
18. Diáková, K., Holcová, V., Šíma, J. & Dušek, J. The Distribution of Iron Oxidation States in a Constructed Wetland as an Indicator of Its Redox Properties. *Chemistry & Biodiversity* **3**, 1288–1300 (2006).
19. Hädrich, A. *et al.* Microbial Fe(II) oxidation by *Sideroxydans lithotrophicus* ES-1 in the presence of Schlöppnerbrunnen fen-derived humic acids. *FEMS Microbiol Ecol* **95**, (2019).

20. Hädrich, A., Heuer, V. B., Herrmann, M., Hinrichs, K.-U. & Küsel, K. Origin and fate of acetate in an acidic fen. *FEMS Microbiology Ecology* **81**, 339–354 (2012).

21. Meyer, J.-M. & Hohnadel, D. Use of nitrilotriacetic acid (NTA) by *Pseudomonas* species through iron metabolism. *Appl Microbiol Biotechnol* **37**, 114–118 (1992).

22. Kathol, M. *et al.* High enzyme promiscuity in lignin degradation mechanisms in *Rhodopseudomonas palustris* CGA009. *Applied and Environmental Microbiology* **0**, e00573-25 (2025).

23. Lill, R. Function and biogenesis of iron–sulphur proteins. *Nature* **460**, 831–838 (2009).

24. Larimer, F. W. *et al.* Complete genome sequence of the metabolically versatile photosynthetic bacterium *Rhodopseudomonas palustris*. *Nat Biotechnol* **22**, 55–61 (2004).

25. Haas, N. W. *et al.* Iron oxidation is regulated by the two-component system, RegSR, and plays a role in photolithoheterotrophic growth in *Rhodopseudomonas palustris*. 2021.08.19.456965 Preprint at https://doi.org/10.1101/2021.08.19.456965 (2021).

26. Young, E. B., Reed, L. & Berges, J. A. Growth parameters and responses of green algae across a gradient of phototrophic, mixotrophic and heterotrophic conditions. *PeerJ* **10**, e13776 (2022).

27. Nikeleit, V. *et al.* Phototrophic Fe(II) oxidation by *Rhodopseudomonas palustris* TIE-1 in organic and Fe(II)-rich conditions. *Environmental Microbiology* **26**, e16608 (2024).

28. Croal, L. R., Jiao, Y., Kappler, A. & Newman, D. K. Phototrophic Fe(II) oxidation in an atmosphere of H2: implications for Archean banded iron formations. *Geobiology* **7**, 21–24 (2009).

29. Melton, E. D., Schmidt, C., Behrens, S., Schink, B. & Kappler, A. Metabolic Flexibility and Substrate Preference by the Fe(II)-Oxidizing Purple Non-Sulphur Bacterium *Rhodopseudomonas palustris* Strain TIE-1. *Geomicrobiology Journal* **31**, 835–843 (2014).

30. Ehrenreich, A. & Widdel, F. Anaerobic oxidation of ferrous iron by purple bacteria, a new type of phototrophic metabolism. *Appl Environ Microbiol* **60**, 4517–4526 (1994).

31. Kopf, S. H. & Newman, D. K. Photomixotrophic growth of *Rhodobacter capsulatus* SB1003 on ferrous iron. *Geobiology* **10**, 216–222 (2012).

32. Zwietering, M. H., Jongenburger, I., Rombouts, F. M. & van 't Riet, K. Modeling of the Bacterial Growth Curve. *Applied and Environmental Microbiology* **56**, 1875–1881 (1990).

33. Alsiyabi, A. *et al.* Synergistic experimental and computational approach identifies novel strategies for polyhydroxybutyrate overproduction. *Metabolic Engineering* **68**, 1–13 (2021).

34. Perni, S., Andrew, P. W. & Shama, G. Estimating the maximum growth rate from microbial growth curves: definition is everything. *Food Microbiology* **22**, 491–495 (2005).

35. Novichkov, P. S. *et al.* RegPrecise 3.0 – A resource for genome-scale exploration of transcriptional regulation in bacteria. *BMC Genomics* **14**, 745 (2013).

36. Dispensa, M. *et al.* Anaerobic growth of *Rhodopseudomonas palustris* on 4-hydroxybenzoate is dependent on AadR, a member of the cyclic AMP receptor protein family of transcriptional regulators. *J Bacteriol* **174**, 5803–5813 (1992).

37. Egland, P. G. & Harwood, C. S. BadR, a New MarR Family Member, Regulates Anaerobic Benzoate Degradation by *Rhodopseudomonas palustris* in Concert with AadR, an Fnr Family Member. *J Bacteriol* **181**, 2102–2109 (1999).

38. Hirakawa, H., Hirakawa, Y., Greenberg, E. P. & Harwood, C. S. BadR and BadM Proteins Transcriptionally Regulate Two Operons Needed for Anaerobic Benzoate Degradation by *Rhodopseudomonas palustris*. *Appl Environ Microbiol* **81**, 4253–4262 (2015).

39. Peres, C. M. & Harwood, C. S. BadM Is a Transcriptional Repressor and One of Three Regulators That Control Benzoyl Coenzyme A Reductase Gene Expression in *Rhodopseudomonas palustris*. *Journal of Bacteriology* **188**, 8662–8665 (2006).

40. Martinez, M. T. P. *et al.* Sensing iron availability via the fragile [4Fe–4S] cluster of the bacterial transcriptional repressor RirA. *Chemical Science* **8**, 8451–8463 (2017).

41. chevrettelab. chevrettelab/gator-gc. (2025).

42. Kumar, A. & Chandra, R. Ligninolytic enzymes and its mechanisms for degradation of lignocellulosic waste in environment. *Heliyon* **6**, e03170 (2020).

43. Rohac, R. *et al.* Structural determinants of DNA recognition by the NO sensor NsrR and related Rrf2-type [FeS]-transcription factors. *Commun Biol* **5**, 769 (2022).

44. Beaumont, H. J. E., Lens, S. I., Reijnders, W. N. M., Westerhoff, H. V. & van Spanning, R. J. M. Expression of nitrite reductase in *Nitrosomonas europaea* involves NsrR, a novel nitrite-sensitive transcription repressor. *Mol Microbiol* **54**, 148–158 (2004).

45. Rodionov, D. A., Dubchak, I. L., Arkin, A. P., Alm, E. J. & Gelfand, M. S. Dissimilatory metabolism of nitrogen oxides in bacteria: comparative reconstruction of transcriptional networks. *PLoS Comput Biol* **1**, e55 (2005).

46. Nellen-Anthamatten, D. *et al. Bradyrhizobium japonicum* FixK2, a Crucial Distributor in the FixLJ-Dependent Regulatory Cascade for Control of Genes Inducible by Low Oxygen Levels. *Journal of Bacteriology* **180**, 5251–5255 (1998).

47. Hördt, A. *et al.* Analysis of 1,000+ Type-Strain Genomes Substantially Improves Taxonomic Classification of Alphaproteobacteria. *Front Microbiol* **11**, 468 (2020).

48. Dehal, P. S. *et al.* MicrobesOnline: an integrated portal for comparative and functional genomics. *Nucleic Acids Res* **38**, D396-400 (2010).

49. Tsoy, O. V., Ravcheev, D. A., Čuklina, J. & Gelfand, M. S. Nitrogen Fixation and Molecular Oxygen: Comparative Genomic Reconstruction of Transcription Regulation in Alphaproteobacteria. *Front Microbiol* **7**, 1343 (2016).

50. Bonnet, M. *et al.* The Structure of *Bradyrhizobium japonicum* Transcription Factor FixK2 Unveils Sites of DNA Binding and Oxidation. *J Biol Chem* **288**, 14238–14246 (2013).

51. Grant, C. R. *et al.* Distinct gene clusters drive formation of ferrosome organelles in bacteria. *Nature* **606**, 160–164 (2022).

52. Pellicer Martinez, M. T. *et al.* Mechanisms of iron- and O2-sensing by the [4Fe-4S] cluster of the global iron regulator RirA. *eLife* **8**, e47804 (2019).

53. Todd, J. D. *et al.* RirA, an iron-responsive regulator in the symbiotic bacterium *Rhizobium leguminosarum*. *Microbiology* **148**, 4059–4071 (2002).

54. Rodionov, D. A., Gelfand, M. S., Todd, J. D., Curson, A. R. J. & Johnston, A. W. B. Computational Reconstruction of Iron- and Manganese-Responsive Transcriptional Networks in α-Proteobacteria. *PLOS Computational Biology* **2**, e163 (2006).

55. Singh, R., Ranaivoarisoa, T. O., Gupta, D., Bai, W. & Bose, A. Genetic Redundancy in Iron and Manganese Transport in the Metabolically Versatile Bacterium *Rhodopseudomonas palustris* TIE-1. *Applied and Environmental Microbiology* **86**, e01057-20 (2020).

56. Kim, M., Kim, W., Park, Y., Jung, J. & Park, W. Lineage-specific evolution of *Aquibium*, a close relative of *Mesorhizobium*, during habitat adaptation. *Appl Environ Microbiol* **90**, e02091-23.

57. Johnston, A. W. B. *et al.* Living without Fur: the subtlety and complexity of iron-responsive gene regulation in the symbiotic bacterium *Rhizobium* and other α-proteobacteria. *Biometals* **20**, 501–511 (2007).

58. Todd, J. D., Sawers, G. & Johnston, A. W. B. Proteomic analysis reveals the wide-ranging effects of the novel, iron-responsive regulator RirA in *Rhizobium leguminosarum* bv. viciae. *Mol Genet Genomics* **273**, 197–206 (2005).

59. Rudolph, G., Hennecke, H. & Fischer, H.-M. Beyond the Fur paradigm: iron-controlled gene expression in rhizobia. *FEMS Microbiology Reviews* **30**, 631–648 (2006).

60. O'Brian, M. R. Perception and Homeostatic Control of Iron in the Rhizobia and Related Bacteria. *Annu. Rev. Microbiol.* **69**, 229–245 (2015).

61. Sandu, C., Olariu, C.-T. & Sandu, R.-C. Technologies for Deviation of Asteroids and Cleaning of Earth Orbit by Space Debris. in *Planetology* (ed. Palaszewski, B.) Chapter 2 (IntechOpen, London, United Kingdom, 2020). doi:10.5772/intechopen.75213.

62. Roelfsema, C. M. *et al.* MRST: What is remote sensing? *A Web Based Toolkit for Using Remote Sensing to Map and Monitor Terrestrial, Marine and Atmospheric Environments* https://sees-rsrc.science.uq.edu.au/rstoolkit/en/html/marine/resources/what-is-remote-sensing.html (2017).

63. Smithsonian Environmental Research Center. Underwater Light and Seagrass. *Ecosystems on the Edge* https://ecosystemsontheedge.org/underwater-light-and-seagrass/ (2013).

64. van Niel, C. B. [1] Techniques for the enrichment, isolation, and maintenance of the photosynthetic bacteria. in *Methods in Enzymology* vol. 23 3–28 (Academic Press, 1971).

65. Reis, M. G. D. & Ribeiro, A. Conversion factors and general equations applied in agricultural and forest meteorology. *AgroM* **27**, (2020).

66. 29.3: Photon Energies and the Electromagnetic Spectrum. *Physics LibreTexts* https://phys.libretexts.org/Bookshelves/College_Physics/College_Physics_1e_(OpenStax)/29%3A_Introduction_to_Quantum_Physics/29.03%3A_Photon_Energies_and_the_Electromagnetic_Spectrum (2016).

67. Bailey, T. L., Johnson, J., Grant, C. E. & Noble, W. S. The MEME Suite. *Nucleic Acids Res* **43**, W39–W49 (2015).

68. Grant, C. E., Bailey, T. L. & Noble, W. S. FIMO: scanning for occurrences of a given motif. *Bioinformatics* **27**, 1017–1018 (2011).

69. Qi, Z. & O'Brian, M. R. Interaction between the Bacterial Iron Response Regulator and Ferrochelatase Mediates Genetic Control of Heme Biosynthesis. *Molecular Cell* **9**, 155–162 (2002).

70. Yang, J., Ishimori, K. & O'Brian, M. R. Two Heme Binding Sites Are Involved in the Regulated Degradation of the Bacterial Iron Response Regulator (Irr) Protein *. *Journal of Biological Chemistry* **280**, 7671–7676 (2005).

71. Todd, J. D., Sawers, G., Rodionov, D. A. & Johnston, A. W. B. The *Rhizobium leguminosarum* regulator IrrA affects the transcription of a wide range of genes in response to Fe availability. *Mol Genet Genomics* **275**, 564–577 (2006).

72. Green, J., Sharrocks, A. D., Green, B., Geisow, M. & Guest, J. R. Properties of FNR proteins substituted at each of the five cysteine residues. *Mol Microbiol* **8**, 61–68 (1993).

73. Wetmore, K. M. *et al.* Rapid Quantification of Mutant Fitness in Diverse Bacteria by Sequencing Randomly Bar-Coded Transposons. *mBio* **6**, 10.1128/mbio.00306-15 (2015).

74. Pechter, K. B., Gallagher, L., Pyles, H., Manoil, C. S. & Harwood, C. S. Essential Genome of the Metabolically Versatile Alphaproteobacterium *Rhodopseudomonas palustris*. *J Bacteriol* **198**, 867–876 (2016).

75. Jiao, Y., Kappler, A., Croal, L. R. & Newman, D. K. Isolation and Characterization of a Genetically Tractable Photoautotrophic Fe(II)-Oxidizing Bacterium, *Rhodopseudomonas palustris* Strain TIE-1. *Appl Environ Microbiol* **71**, 4487–4496 (2005).

76. Dehio, C. & Meyer, M. Maintenance of broad-host-range incompatibility group P and group Q plasmids and transposition of Tn5 in *Bartonella henselae* following conjugal plasmid transfer from *Escherichia coli*. *Journal of Bacteriology* **179**, 538–540 (1997).

77. Khan, S. R., Gaines, J., Roop, R. M. & Farrand, S. K. Broad-Host-Range Expression Vectors with Tightly Regulated Promoters and Their Use To Examine the Influence of TraR and TraM Expression on Ti Plasmid Quorum Sensing. *Appl Environ Microbiol* **74**, 5053–5062 (2008).

78. Quandt, J. & Hynes, M. F. Versatile suicide vectors which allow direct selection for gene replacement in gram-negative bacteria. *Gene* **127**, 15–21 (1993).

79. Jiao, Y. & Newman, D. K. The *pio* Operon Is Essential for Phototrophic Fe(II) Oxidation in *Rhodopseudomonas palustris* TIE-1. *Journal of Bacteriology* **189**, 1765–1773 (2007).

80. Gupta, D. *et al.* Photoferrotrophs Produce a PioAB Electron Conduit for Extracellular Electron Uptake. *mBio* **10**, e02668-19 (2019).

81. Potter, S. C. *et al.* HMMER web server: 2018 update. *Nucleic Acids Res* **46**, W200–W204 (2018).

82. Valasatava, Y., Rosato, A., Banci, L. & Andreini, C. MetalPredator: a web server to predict iron–sulfur cluster binding proteomes. *Bioinformatics* **32**, 2850–2852 (2016).

83. Bird, L. J. *et al.* Nonredundant Roles for Cytochrome c2 and Two High-Potential Iron-Sulfur Proteins in the Photoferrotroph *Rhodopseudomonas palustris* TIE-1. *Journal of Bacteriology* **196**, 850–858 (2014).

84. Dixon, R. & Kahn, D. Genetic regulation of biological nitrogen fixation. *Nat Rev Microbiol* **2**, 621–631 (2004).

85. Nomata, J., Kitashima, M., Inoue, K. & Fujita, Y. Nitrogenase Fe protein-like Fe-S cluster is conserved in L-protein (BchL) of dark-operative protochlorophyllide reductase from *Rhodobacter capsulatus*. *FEBS Lett* **580**, 6151–6154 (2006).

86. Boll, M. Key enzymes in the anaerobic aromatic metabolism catalysing Birch-like reductions. *Biochimica et Biophysica Acta (BBA) - Bioenergetics* **1707**, 34–50 (2005).
